# Synthesis of Cyclopropene-Modified Fatty Acids Allows Single Cell Quantification of Uptake By Immune Cells

**DOI:** 10.1101/2025.07.11.664320

**Authors:** Luuk Reinalda, Graham A Heieis, Diana Torres-García, Marouane el Boujadayni, Kas Steuten, Kristine Bertheussen, Laura I. Bogue, Jeroen M. Punt, Wouter P. F. Driever, Connor Corrigan, Xinyuan Wang, Bart Everts, David Finlay, Mario van der Stelt, Sander I. van Kasteren

## Abstract

The activation of T-cells is heavily shaped by the nutrients that are available during activation. Fatty acids can have highly pleiotropic effects in this process. On the one hand, they are essential for driving T-cell activation, yet on the other hand they can affect curtailed T-cell activation. These differences are likely dependent on the nature and absolute uptake of the fatty acids. Quantifying the uptake of specific FAs by individual cells and studying the effect on the phenotype of the cell is currently not possible with existing tools. Here we therefore use the live-cell compatible Inverse Electron-Demand Diels-Alder reaction combined with the synthesis of saturated, unsaturated and polyunsaturated fatty acids carrying the 1-carbon cyclopropene click group. Single cell uptake studies of these various clickable FAs in primary immune cell mixtures show highly divergent uptake behaviour between different immune cells, with polyunsaturated fatty acids markedly preferred by all immune cells tested.

## Introduction

Fatty acids (FAs) play essential roles in every mammalian cell type.[1, 2] They serve as structural building blocks of membranes[3], as fuel to highly energy demanding cells, such as activated immune cells and cancer cells[4], and as signalling mediators, particularly in immunological processes.[5-8] The identification of the (patho)physiological roles of FAs in immune activation and the shaping and resolution of the immune response is leading to the discovery of new therapeutic targets.[9-11]

FAs used by immune cells can be synthesized de novo by the enzymes amino acetyl carboxylase and fatty acid synthase.[12] However, many cell types, including immune cells also take up exogenous FAs to fuel their metabolic and signalling demands, shaping the specific immune response.[13, 14] Specific fatty acids can affect particular aspects of the immune response. For example, exogenous oleic acid is essential for T cell activation[15], but can also have more specific effects on subset skewing.[16, 17] The saturated fatty acids palmitic and stearic acid can be toxic to activating T cells[18], which can in part be rescued by other FAs.[19] Linoleic, palmitoleic, and arachidonic acid have been shown to both potentiate T-cell activation[20], but also to suppress it, perhaps in a concentration dependent manner.[6, 16, 20-22]

These contradicting biological findings highlight the need to be able to study the uptake of FAs in immune cells at the single cell level, as their effects on the target cell is likely dependent on both the nature and amount of the specific FA a cell has taken up. Fluorophore labelled fatty acid analogues (e.g. BODIPYTM-C12/16, TopFluor® oleic acid) were the first reagents to be developed with this aim in mind.[23, 24] They mimic long chain fatty acids by either replacing the end of the hydrophobic tail with, or adding a fluorophore on to it. The approach is live-cell compatible[25], but the large pendant aromatic fluorophore disrupts the native structure of the fatty acid, with the fluorophore likely having much more of an impact on FA structure than the chain length and/or the presence/absence of double bonds, limiting their suitability for studying uptake of lipids with subtle structural differences.[26]

Bioorthogonal click chemistry has been used to overcome this problem.[27, 28] Instead of using large pendant fluorophores for direct detection, fatty acid analogues bearing the 2-3 atom alkyne and azide groups were developed.[29] These small chemical functionalities allow for fluorophore labelling after completion of the biological time courses, whilst keeping structural fidelity to natural FA.[30-32] Alkyne analogues of saturated fatty acid (SFA) palmitic acid, and monounsaturated fatty acid (MUFA) oleic acid have, for example, been used to mark differences in the uptake of exogenous FAs in differently polarized bone marrow derived macrophages.[33] They were also employed to study the CD36-mediated trans-endothelial cells transport[34], and more recently to quantify the subcellular localization of different lipids after uptake, when combining alkynes with photo-crosslinking.[35] An alkyne analogue of arachidonic acid was also reported and used to reveal the critical role of this polyunsaturated fatty acid (PUFA) in megakaryocyte differentiation.[36] An alkyne-containing analogue of the ω-3 PUFA α-linolenic acid was used to unravel S. Aureus capacity to escape antimicrobial FAs.[37]

Whilst the Cu-catalysed azide-alkyne cycloaddition (CuAAC) is a very powerful approach for labelling lipids, one major downside is that the chemistry required for their labelling requires a toxic copper catalyst. To combat this, non-toxic click reactions, such as the strain-promoted azide-alkyne cycloaddition (SPAAC)[38] and the inverse demand Diels-Alder cycloaddition reaction (IEDDA) were developed.[39-41] A downside, specific to lipid labelling, is that these require the use of either polar click groups such as azides, or bulkier bioorthogonal groups, such as cyclooctynes, tetrazines, and trans-cyclooctenes in the lipid, which too – like fluorophore modification – result in major structural alterations to the lipids.[23]

One group that can combine the favourable properties of live cell labelling and stands out in this context: 1,2- and 1,3-cyclopropenes (Cps) are the smallest strained alkenes that can react in the IEDDA reaction with electron deficient dienes like tetrazines (Figure 1).[42-45] The cyclopropene is only moderately larger than the alkynes commonly used for fixed-cell FA-labelling. The reaction rate is similar to that of the CuAAC38, without the need for a toxic catalyst. This has led to a flurry of activity for using cyclopropenes in chemical biology. Cp-amino acid analogues have been applied to protein labelling[46-49], nucleotide analogues to label RNA/DNA[50, 51], and carbohydrate analogues for glycan imaging.[44, 52-54] Only a few examples of cyclopropenes used in the study of lipid uptake have been reported. Devaraj and co-workers attached a 1,3-cyclopropene tag to the polar head of phospholipids and incorporated those in membranes of a human breast cancer cell line, which was visualized by confocal microscopy.[43] Our group has recently reported the use of the naturally occurring cyclopropene containing lipid, sterculic acid, an analogue of oleic acid bearing an extra methylene bridging the double bond as a tool for imaging lipid uptake.[55, 56] This lipid could be incorporated in membranes of an immortalized cell line and be used for live-cell imaging and chemical proteomics.[55] However, no other naturally occurring cyclopropene lipids exist that can be used analogously to this oleic acid analogue.

**Figure 1.**
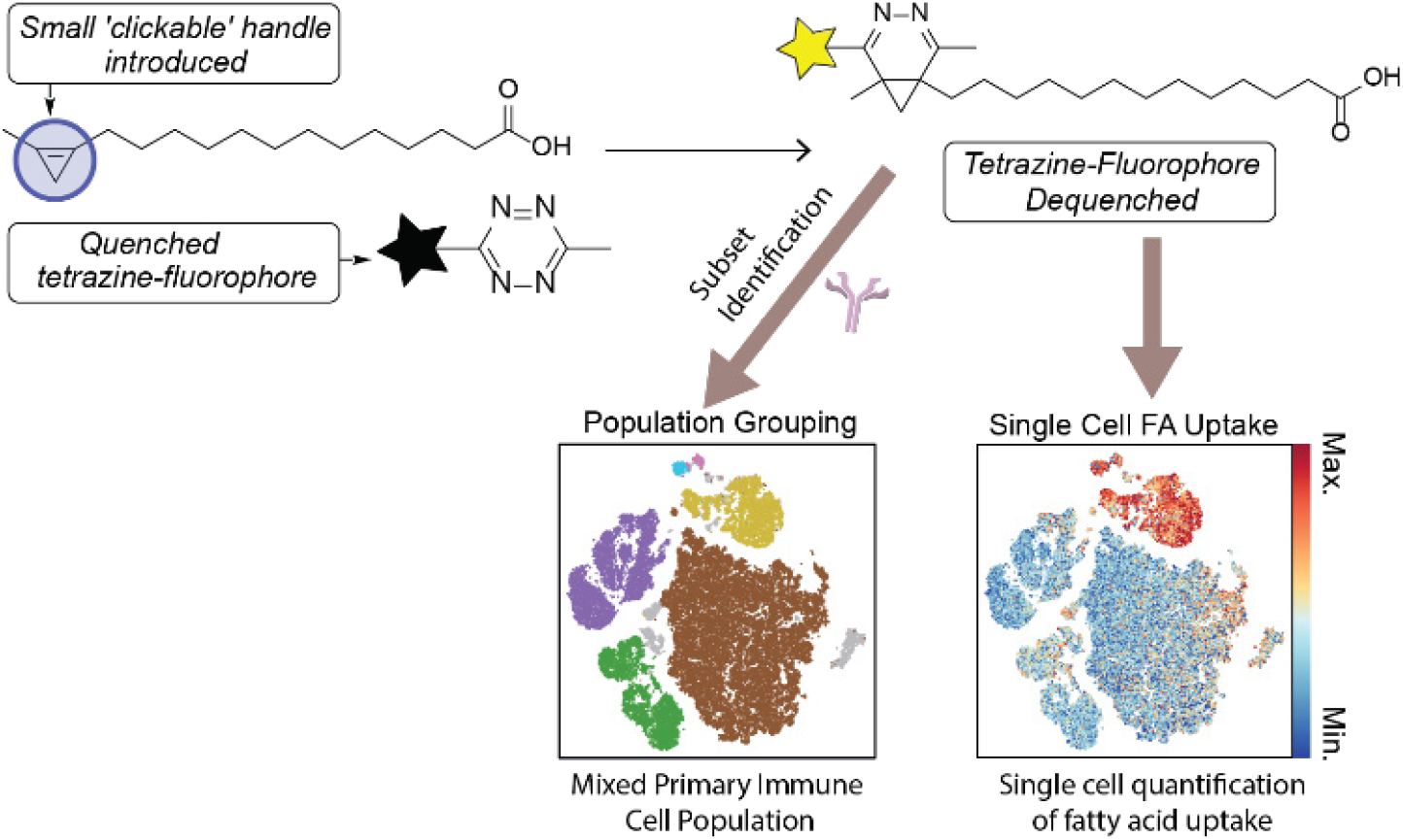
Overview of the approach and synthetic targets. Mixtures of primary immune cells were incubated with cyclopropene-modified fatty acids. These nominal 1-carbon click groups are reactive in the inverse electron-demand Diels-Alder reaction to allow the quantification of uptake of fatty acids in live primary immune cells at the single cell level. The mildness of the approach allows easy combination with complex flow cytometry and analysis of live cells.

Here we tackle this lack of IEDDA-reactive Cp-containing lipids, we here describe the synthesis of sterculic acid and five new synthetic cyclopropene fatty acid analogues (CpFAs) (Figure 2A, 1-6) and assess their uptake by T-cells in mixed immune populations. All 6 CpFAs showed equal reactivity with tetrazines, allowing us to show that activated T cells take up more polyunsaturated fatty acids compared to monounsaturated and saturated fatty acids.

**Figure 2.**
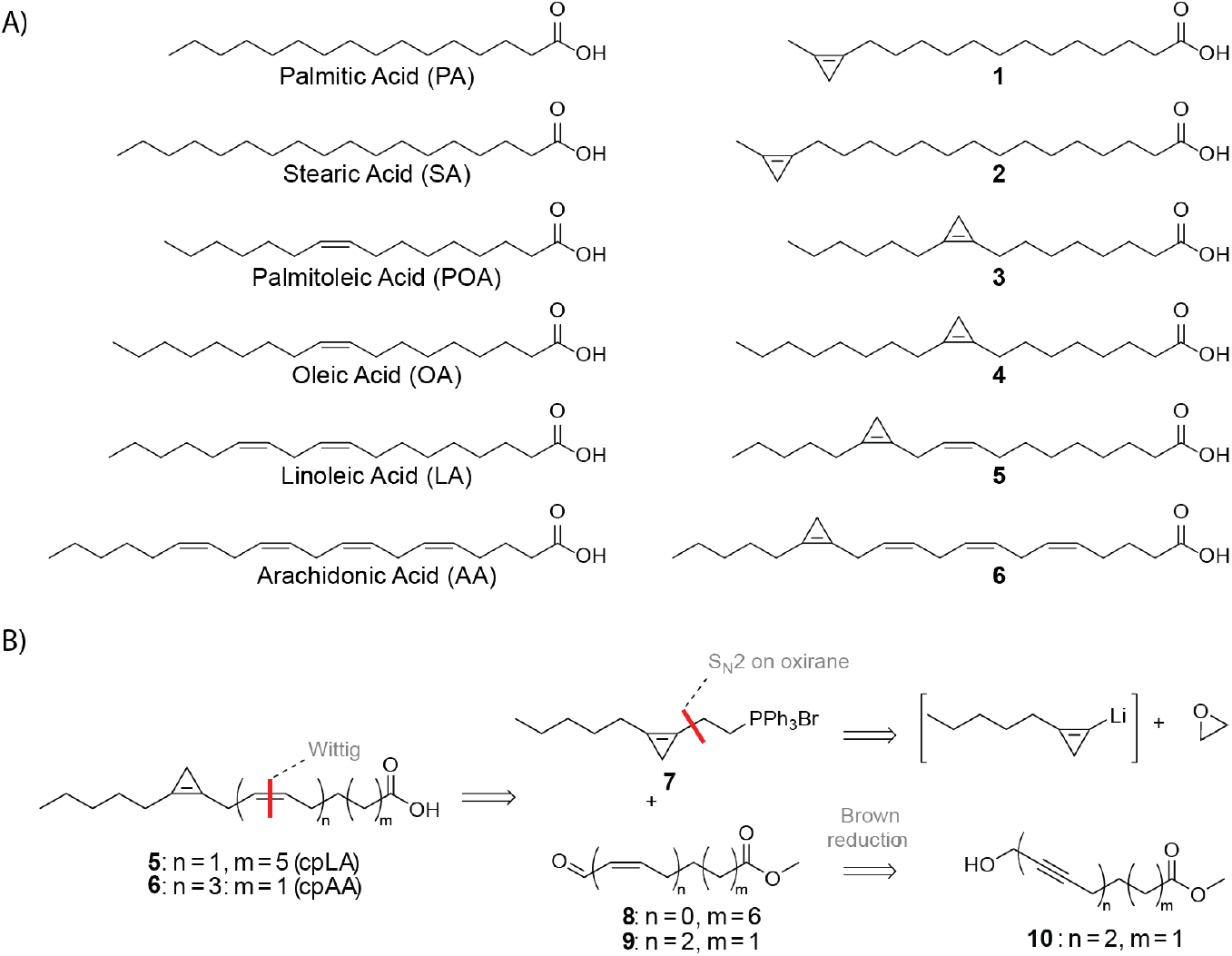
A) (Left side) Key fatty acids in T-cell metabolism and their cyclopropene containing counterparts made in this manuscript. B) Retrosynthetic analysis for PUFAs linoleic acid 5 and arachidonic acid 6.

## Results and Discussion

### Design and Synthesis of Cyclopropene Fatty Acids

Our first aim was to design Cp-containing analogues of six key fatty acids important in immunology with minimal structural modification. For the saturated fatty acids, we envisaged that fatty acids with an ω-terminal Cp-group would best mimic the saturated FAs palmitic and stearic acid, leading to the chosen design of 1, and 2. For any lipid bearing a single double bound, we envisaged a formal replacement of the double bond with the 1,2-cyclopropene group. Cp-palmitoleic acid 3 was thus designed following nature’s design of sterculic acid 4. For the polyunsaturated FAs, we chose to substitute the ω-6 double bonds of linoleic acid and arachidonic acid to yield designs 5 and 6 (Figure 2B). We chose to keep the linear chain length identical to that of the parent lipid, leading to a nominal size increase of one carbon in the Cp-analogues.

The polyunsaturated CpFAs were the most challenging synthetically. Retrosynthetic analysis (Figure 2B) had us disconnect across the double bond adjacent to the Cp-group, which can be formed using a Wittig reaction. This approach identified triphenylphosphonium bromide building block 7 as a key intermediate to be accessed by an SN2-reaction between a cyclopropyl-lithium species and oxirane.[57] For the synthesis of the aldehyde building block 9, required for the synthesis of CpAA 6, a Brown reduction of the bis-alkyne building block 10 to bis-cis-alkene 21 was proposed.

The synthesis of polyunsaturated CpFAs 5 and 6 (Scheme 1) was achieved via a phosphonium bromide was positioned on the same fragment as the cyclopropene.[58] Incidentally, the inverse approach, where the aldehyde was introduced on the Cp-containing fragment, did not work due to instability of the aldehyde (Scheme S2). Alcohol 12 was synthesized by ring-opening of ethylene oxide, catalysed by BF3·OEt in presence of the lithiated cyclopropene nucleophile from 11. Appel-like conditions using N-bromosuccinimide on our 1,2-substituted substrate afforded stable cyclopropene bromide 13 in 59% yield, as direct alkylation with dihaloalkanes did not work (Scheme S1). An overnight substitution reaction with triphenylphosphine under reflux in acetonitrile yielded the target cyclopropene phosphonium bromide 7 in 23% yield over 3 steps. Aldehyde 8 was synthesized in five steps from a diol by protecting the alcohol on one end with a silyl to yield compound 14, which was oxidized and protected to yield methyl ester 15. The silyl ether protecting group was removed and the alcohol 16 oxidized affording aldehyde 8. The phosphorus ylide of 7 was formed in situ using NaHMDS and reacted with aldehyde 8 to yield the cyclopropene linoleic methyl ester 18. Subsequent saponification yielded 5, to the best of our knowledge the first example of the synthesis of a cyclopropene-containing PUFA analogue. The synthesis of the polyunsaturated arachidonic acid-analogue 6 was again achieved using a late stage Wittig reaction to form the C-C-bond between aldehyde 9 and phosphonium bromide 7, but instead uses the bis-alkyne 10 as key intermediate (Scheme 1C) that could later be reduced stereoselectively to the bis-cis-alkene[59-61], as efforts to perform the Wittig using the bis-alkene had failed (Scheme S3).

**Scheme 1.**
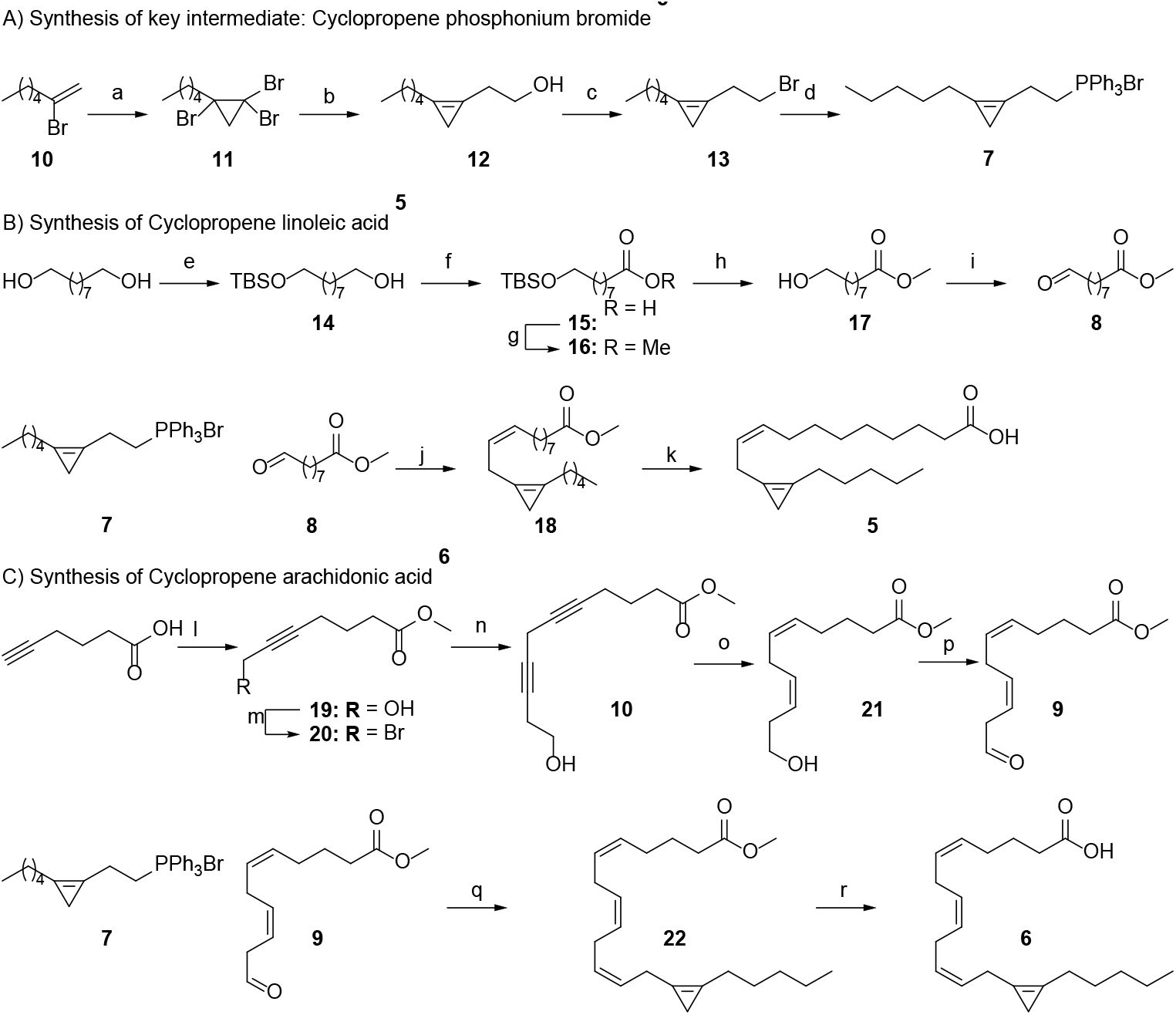
Synthesis of PUFAs linoleic acid (5) and arachidonic acid (6). A) Synthesis of key intermediate cyclopropene phosphonium bromide **7**. Reagents and conditions: a) BBr_3_, DCM, - 78 °C to rt, 3 h, 52% yield. b) *n*-Buli, BF_3_·OEt_2_, oxirane, THF, −78 °C to 0 °C, 2 h, 49% yield. c) NBS, PPh_3_, DCM, 0 °C to rt, 2 h, 59% yield. d) PPh_3_, MeCN, 82 °C, 18 h, 82% yield. B) Synthesis of cyclopropene linoleic acid **5**. Reagents and conditions: e) TBSCl, imidazole, DCM, rt, 3 h, 64% yield. f) TEMPO / BAIB, 2:1 MeCN / water, rt, 3 h, 92% yield. g) MeI, K_2_CO_3_, DMF, rt, 18 h, 86% yield. h) TBAF, DCM, 0 °C to rt, 5 h, 93% yield. i) DMP, DCM, 0 °C to rt, 1 h, 78% yield. j) NaHMDS, THF, −78 °C to 0 °C, 2.5 h, 44% yield. k) NaOH (aq.), 1:1 THF / EtOH, 70 °C, 2 h, 73% yield. C) Synthesis of cyclopropene arachidonic acid **6**. Reagents and conditions: l) EtMgBr, PFA, THF, 0 °C to 75 °C, 18 h, then: SOCl_2_, MeOH, 0 °C to rt, 1 h, 51% yield. m) NBS, PPh_3_, DCM, 0 C to rt, 1 h, 82% yield. n) CuI, NaI, K_2_CO_3_, but-3-yn-1-ol, DMF, rt, 18 h, 72% yield. o) NaBH_4_, Ni(OAc)_2_ · 4 H_2_O, ethylenediamine, H_2_ (g), MeOH, rt, 2 h, 27% yield. p) DMP, DCM, 0 °C to rt, 1 h, 37% yield. q) NaHMDS, THF, −78 °C to 0 °C, 2.5 h, 54% yield. r) NaOH (aq.), 1:1 THF / EtOH, 70 °C, 2 h, quant. yield.

Alkynylation of paraformaldehyde with 5-hexynoic acid using an excess of EtMgBr followed by esterification yielded methyl ester 19, which was converted into the bromide 20 with an Appel-like reaction. Then, a copper(I) mediated cross-coupling with but-3-yn-1-ol afforded skipped diyne 10. A selective reduction of the alkynes to alkenes was attempted using poisoned P-2 Nickel, as Lindlar’s catalyst yielded inseparable E/Z mixtures.[62, 63] HPLC-purification was required to separate diene 21 from its by-products. A Wittig reaction between aldehyde 9 and phosphonium bromide 7 yielded cyclopropene arachidoyl methyl ester 22 that could be saponified to yield 6 (Scheme 1C).

Various routes towards the synthesis of sterculic acid have been reported.[64-67] Due to the structural similarity of Cp-palmitoleic acid 3 to sterculic acid, we opted for the most-used synthetic route which involves a rhodium catalysed [2+1] cycloaddition to form the cyclopropene moiety (Scheme S4). Nevertheless, this route was deemed unpractical due to experimental difficulty and poor reproducibility.[68] Saturated CpFAs 1 and 2 and MUFA 4 were therefore synthesized using a tribromocyclopropane 23 as a cyclopropene precursor (Scheme 2).[69] Treatment of 26 and 27 with two equivalents of n-butyllithium (n-BuLi) yielded a lithiated cyclopropene nucleophile via a double lithium-halogen exchange and an elimination[70, 71], that could be reacted with a diiodoalkane to form the iodocyclopropenes. In the first step, bromination of the terminal alkyne with BBr3 yielded bromoalkene 24. A dibromocarbene-species was formed from bromoform by NaOH in a phase transfer catalytic system, which was reacted with the readily available alkene 23 and compound 24 to yield tribromocyclopropanes 25 and 26. The reaction of the lithiated cyclopropene nucleophile with an excess of diiodoalkane afforded iodocyclopropenes 27-29. These were converted to nitriles 30-32 by reacting them with sodium cyanide followed by basic hydrolysis to give the cyclopropene fatty acids 1, 2 and 4 in 2.5 – 11.5% yield over 5 steps. Although the yield was still low, the experimental simplicity of the individual steps and higher reproducibility made this the route of choice for the synthesis of compounds 1, and 2.

**Scheme 2.**
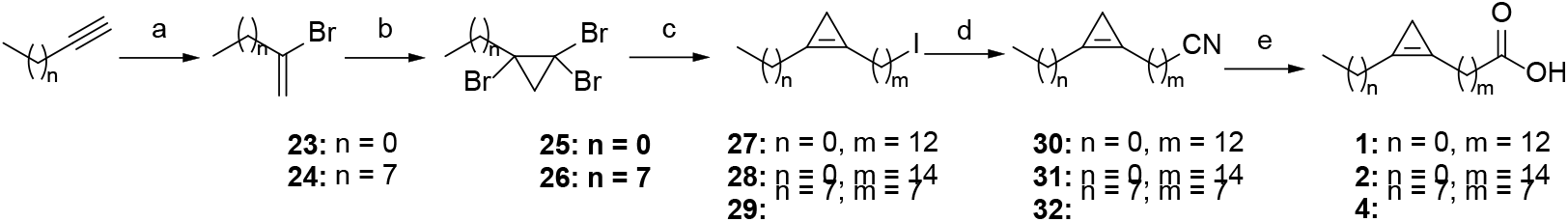
Synthesis of SFAs and MUFA 1, 2 and 4. Reagents and conditions: a) BBr_3_, DCM, - 78 °C to rt, 3 h, 78% yield. b) NaOH, CTAB, CHBr_3_, DCM, 0 °C to rt, 18 h, 47% – 57% yield. c) n-BuLi, diiodoalkane, THF, −78 °C to 0 °C, 4 h, 27 – 56% yield. d) NaCN, DMSO, 90 °C, 1 h, 68 – 89% yield. e) NaOH, EtOH / water, reflux, 18 h, 37 – 63% yield.

### CpFA Uptake in Mouse Immune Cells

With a library of six cyclopropene fatty acid analogues of FAs that have been implicated in T-cell activation biology[72, 73], we next assessed the suitability of these analogues to study fatty acid uptake in live immune cell populations. We opted against the use of cell lines, as their neoplastic nature heavily impacts their nutrient uptake and use.[74] Instead, we chose to optimise the uptake method and IEDDA reaction for CpFAs 1-6 in primary splenocytes which were obtained by mechanical homogenization of mouse spleens.[75] After red blood cell lysis, this population consists of a mixture of many key immune cells in various states of activation, such as dendritic cells, macrophages, T-cells and B-cells. The flow cytometric identification with well-characterized antibodies is routine (Table S1) and would allow for the facile quantification of uptake of 1-6 at the single cell level.[76]

After pulsing splenocytes with the CpFAs, we first assessed uptake by performing the click reaction with tetrazine-AF488. All six CpFA showed active uptake at 37 °C compared to cells that were incubated with CpFAs on ice as a negative control (Figure 2A/B, S1A). However, the background of this reaction detected in the ice-controls was very high. We attempted to increase the signal-to-noise ratios by titrating both the levels of sterculic acid (Figure 2B) and the concentration of AF488-Tz used (Figure 2B/C). The concentration of tetrazine had the largest impact on detection levels, as performing the reaction without CpFA treatment led to a high background fluorescence (Figure 2B). To resolve the high non-specific binding of the tetrazine-AF488, we tested tetrazine-BODIPY FL (Figure 2D) as an alternative, which is quenched until reaction with Cp.[55] Using this method, the background of the click reaction in the absence of CpFAs was comparable to splenocytes with no reaction done (Figure 2D-F), and similar values were calculated for both tetrazine-conjugates after background subtraction (Figure 2C, F). Importantly, the same uptake pattern was observed for both reactions across specific immune cell populations (Figure S1B). The improved signal-to-noise ratio as indicated by the calculated stain-index, however, makes the BODIPY more suitable for flow-cytometry based assays (Figure S1C). Moving forward we opted to use 25 µM of CpFA, which falls in the range of physiological concentration in all parent lipids[77], and 1 µM of tetrazine-BODIPY FL to prevent risk of fluorescent spillover.

One factor to consider in these assays, is the potential difference in dequenching upon reaction of BODIPY FL with each of the CpFAs. We assessed this by performing an in vitro fluorescence turn-on assay where the six CpFAs (25 µM) were incubated with tetrazine-BODIPY FL (1 µM) in PBS for 30 min and the fluorescent signal (λex = 447 nm, λem = 530 nm) measured (Table S2) to create an adjusted value (relative to linoleic acid). Applying the adjusted values to the intensities of pulsed splenocytes did not alter the overall pattern of uptake across CpFAs (Figure S1E). To assess whether the CpFAs are suitable analogues, we carried out a competition assay where CpFAs (25 µM) were simultaneously pulsed with 25, 50 or 100 µM of the parent FA (Figure S1F). Competition of palmitic acid and arachidonic acid were complete (i.e. only 20% of signal remained in presence of 100 µM – a 4-fold excess of parent lipid). Uptake inhibition of stearic acid, palmitoleic acid, oleic acid and linoleic acid was not complete, with 40% to 70% of signal remaining in presence of 100 µM of the parent lipid, which was also observed for alkyne lipids.[78] Application of this optimized protocol to the study of the uptake of the six CpFAs showed active uptake at 37 °C that was reduced when cells were left on ice (Figure 3A,B). Cell viability was not affected by the presence of the CpFAs, as determined by staining with a live exclusion dye following uptake, which was detected by flow cytometry (Figure S1D).

**Figure 3.**
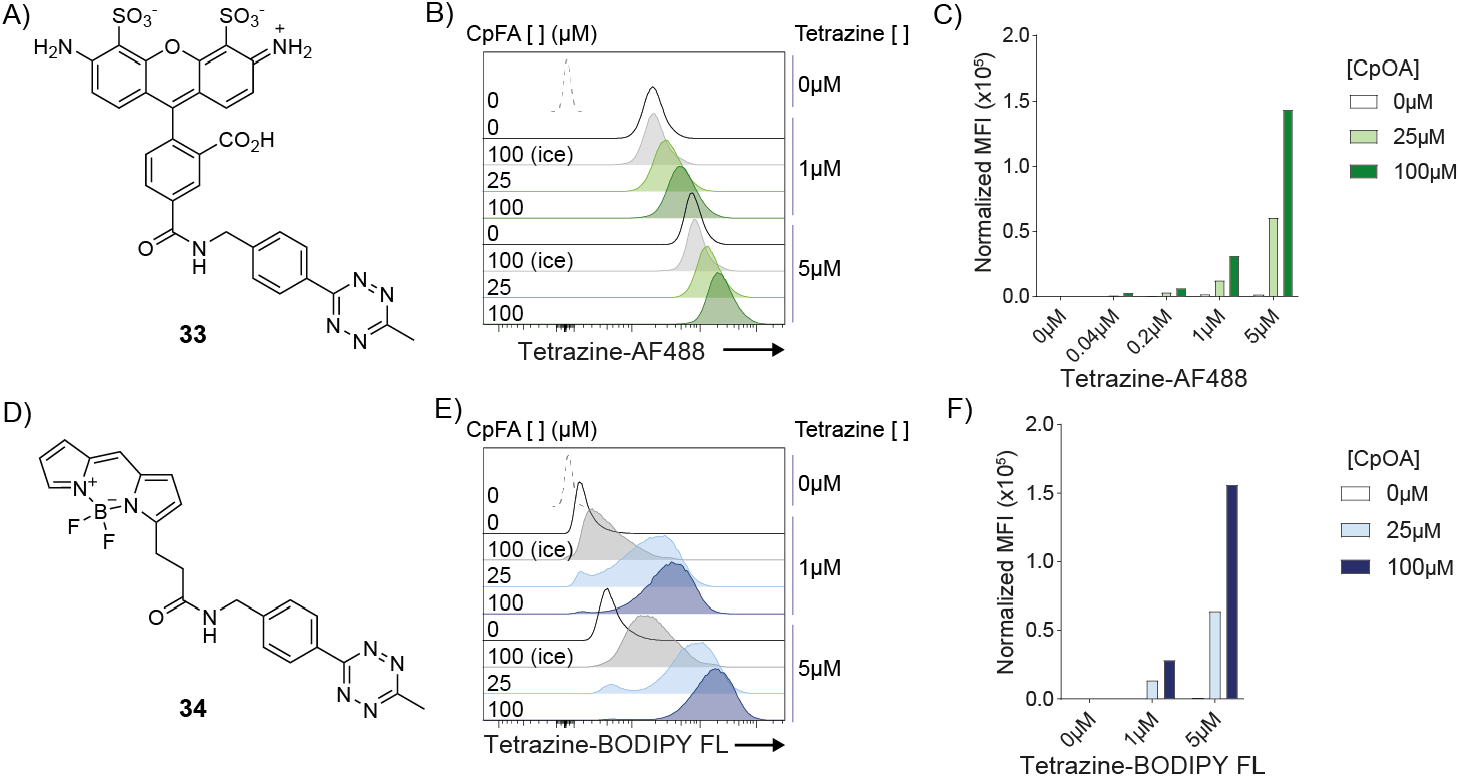
Optimization of ex-vivo uptake of sterculic acid 4 in mouse splenocytes. Splenocytes were incubated with sterculic acid (0, 25 or 100 µM) for 30 minutes prior to fixation and subsequent click-reaction with either AF488-tetrazine **33** (0.04, 0.2, 1 or 5 µM) or BODIPY-tetrazine (0, 1 or 5 µM). A, D) Chemical structures of AF488-tetrazine **33** and tetrazine-BODIPY FL **34**. B) Histograms of signal after uptake. Green: 37 °C, grey: cold control. C) Bar chart of the data shown in B. E) Signal resulting from reaction between **4** after uptake and tetrazine-BODIPY FL **34**. Green: 37 °C, grey: cold control. F) Bar chart of the data shown in E.

Having optimized the protocol and validated our CpFAs as analogues for their native counterparts, we aimed to determine how FA were differently utilized across immune populations from the mouse spleen. Following the uptake assay and click reaction, cells were therefore stained using fluorescent-conjugated antibodies targeting receptors unique to specific immune populations, allowing their identification by flow cytometry. To assess which immune cells take up the various FAs, we performed an unsupervised dimensional reduction to visualize the correlation between immune cell type and FA-uptake, using palmitic acid as an example (Figure 3C). The resulting t-distributed Stochastic Neighbour Embedding (tSNE) plot indicated particularly high uptake in cells from the myeloid lineage, including monocytes and dendritic cells, whereas lymphoid cells, such as T and B cells, demonstrated comparatively low uptake. We confirmed these findings by manually gating and determining the detection of CpFA for each major immune subset (Figure 3D) as we found a similar pattern of uptake for all FA species tested across the different cell types (Figure 3E). Notably, all immune cells showed a preference for the PUFAs, Cp-linoleic and Cp-arachidonic acid, followed by the SFA Cp-palmitic acid. Therefore, our library of novel CpFAs has the potential to reveal specific needs for particular fatty acid species across diverse primary immune cell populations.

Figure 4. Uptake of CpFA library 1-6 in splenocytes. A) Representative histograms of the uptake of six CpFAs (37 °C) including autofluorescence (No click), non-specific binding of tetrazine-BODIPY FL (Background click) and passive uptake (ice). B) Change in normalized MFI of CpFA (25 µM) uptake with cold control included. C) Dimensionality reduction using tSNE was performed on splenocytes for Cp-palmitic acid uptake. Right panel shows splenocytes population including CD19+/CD3 B cells, CD11b+/Ly6Chi monocytes, Ly6Chi neutrophils, CD3+/CD19 CD4+ T cells, CD3+/CD19 CD8+ T cells and MHCII+/CD11c+ dendritic cells. Bottom panel shows the corresponding Cp-palmitic acid uptake. D) Corresponding manual gating strategy. E) Change in for passive uptake corrected MFI of CpFA uptake in different immune cell subsets. Data shown in this figure represents uptake performed in splenocytes harvested from individual mice (N = 3) with no technical replicates (n = 1).

**Figure 4.**
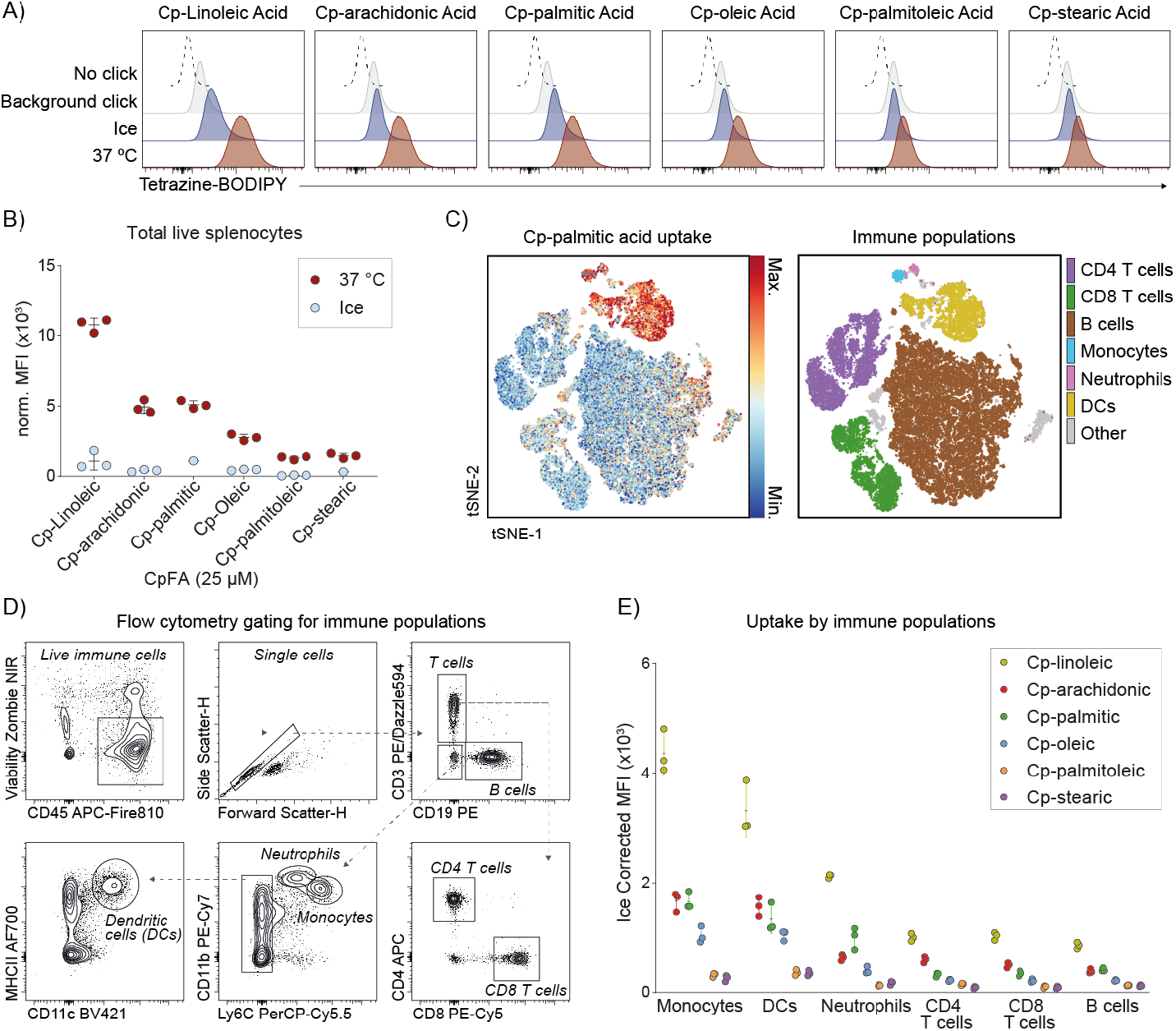
Uptake of CpFA library 1-6 in splenocytes. A) Representative histograms of the uptake of six CpFAs (37 °C) including autofluorescence (No click), non-specific binding of tetrazine-BODIPY FL (Background click) and passive uptake (ice). B) Change in normalized MFI of CpFA (25 µM) uptake with cold control included. C) Dimensionality reduction using tSNE was performed on splenocytes for Cp-palmitic acid uptake. Right panel shows splenocytes population including CD19^+^/CD3 B cells, CD11b^+^/Ly6C^hi^ monocytes, Ly6C^hi^ neutrophils, CD3^+^/CD19 CD4^+^ T cells, CD3^+^/CD19 CD8^+^ T cells and MHCII^+^/CD11c^+^ dendritic cells. Bottom panel shows the corresponding Cp-palmitic acid uptake. D) Corresponding manual gating strategy. E) Change in for passive uptake corrected MFI of CpFA uptake in different immune cell subsets. Data shown in this figure represents uptake performed in splenocytes harvested from individual mice (N = 3) with no technical replicates (n = 1).

**Figure 5.**
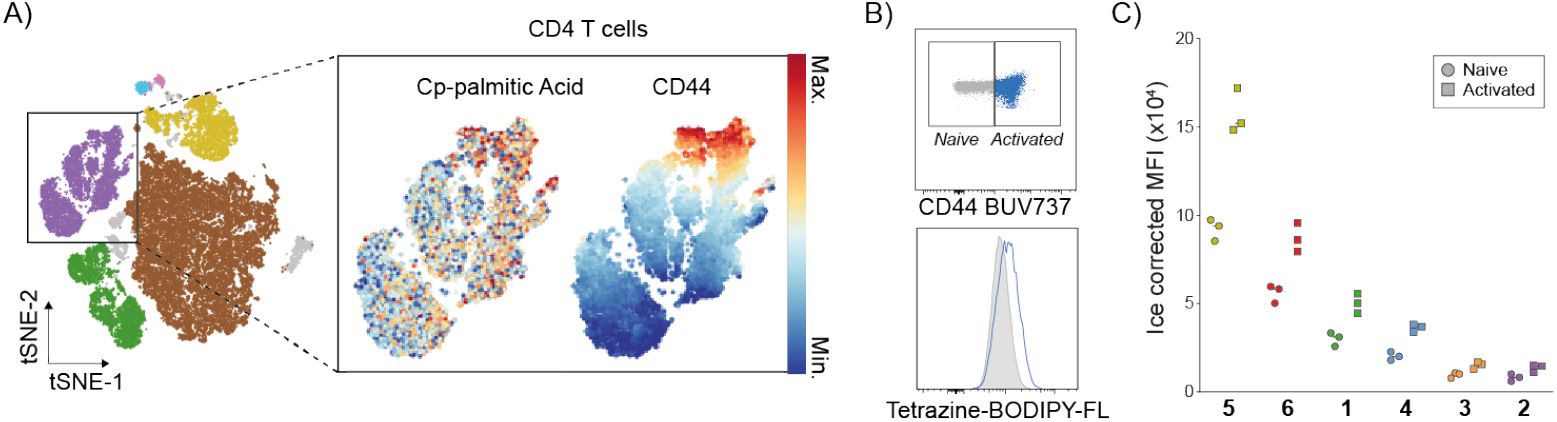
A) Dimensionality reduction using tSNE on CD4^+^ T cells expressing activation marker CD44 for Cp-palmitic acid uptake. B) Top panel shows the manual gating strategy for naïve CD44^-^ T cells and activated CD44^+^ T cells, bottom panel the representative histogram. C) Difference in corrected MFI of CpFA uptake between naïve and activated T cells. Data shown in this figure represents uptake performed in splenocytes harvested from individual mice (N = 3) with no technical replicates (n = 1).

FA uptake in immune cells is rapidly regulated during activation, or within the same cell type can differ depending on their functional state. CD4+ T cells, for instance, rapidly increase their need for environmental FA when stimulated by the matched antigen to their specific T cell receptor, allowing them to divide and produce effector molecules. Expression of the T cell activation marker CD44 appeared to correlate with palmitic acid uptake, which we further confirmed with manual gating and analysis (Figure 4A,B) Indeed, all analogues showed an increase in uptake in activated compared to naïve CD4+ T cells, however this was less evident for palmitoleic and stearic acid (Figure 4C).

## Conclusion

We here report the use of a cyclopropene long chain fatty acid analogue library, covering the three main types of saturation, to investigate fatty acid uptake of individual immune cells in complex cell populations. An optimized synthetic methodology was applied to synthesize SFA and MUFA analogues and we developed new synthetic methodology to synthesize the first ever reported cyclopropene PUFA analogues. An initial uptake experiment revealed a high background was detected when using tetrazine-AF488 due to non-specific binding. Hence, the quenched tetrazine-BODIPY FL was employed resulting in a much better resolution. The uptake of palmitic acid was visualized in a tSNE plot revealing a higher FA uptake for cells from the myeloid, for instance monocytes and dendritic cells, compared to lymphoid cells like T and B cells. Within all different immune cell types a similar pattern was observed where immune cells have a preference for PUFAs, followed by MUFA oleic acid and SFA palmitic acid. Furthermore, we showed that different cell states (e.g. activated or naïve T cells) lead to a quantitative change in exogenous FA uptake. With this we established a platform which can be applied to study relationships between nutrient uptake and immune cell function on a new level.

## Supporting Information

Supplementary figure S1-2, table S1 and schemes S1-4 Supplementary figure S1, tables S1-2, and Schemes S1-4 and experimental information are associated with this manuscript. The authors have cited additional references within the Supporting Information.[79-96]

## Acknowledgements

SvK was funded by an ERC Consolidator Grant (Grant number 865175) and internal funding from the NWO-Gravitation-funded Institute of Chemical Immunology and Institute for Chemical Neuroscience. MvdS was funded by a Netherlands Organization for Scientific Research VICI-grant (Grant number 724.017.002) and by the Institute for Chemical Neuroscience.

## Notes

### Competing Interest Statement

The authors have declared no competing interest.

### Summary of Updates

The accent of the chemistry had been switched towards the method development and synthesis of the key compounds and away from the immunology.

