## Supplementary Information for "Synthesis of Cyclopropene-Modified Fatty Acids Allows Single Cell Quantification of Uptake By Immune Cells"

Classified as Internal | Intern

Classified as Internal | Intern

### Supplementary Schemes and Figures

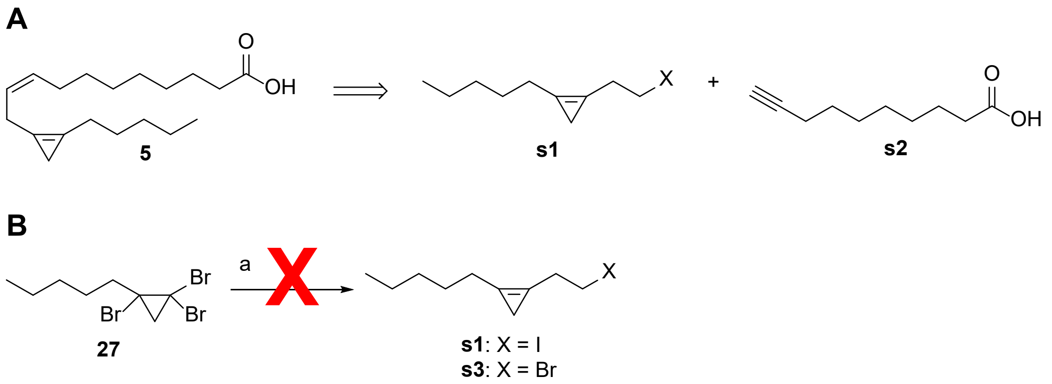

**Scheme S1. Synthetic strategy for PUFA synthesis**. A) Retrosynthetic analysis of compound **5**. B) Attempted route to form cyclopropene halides **s1** or **s3**. Reagents and conditions: a) *n*-Buli, THF, 1,2-diiodoethane or 1-bromo-2-iodoethane, -78 °C to 0 °C, 4 h.

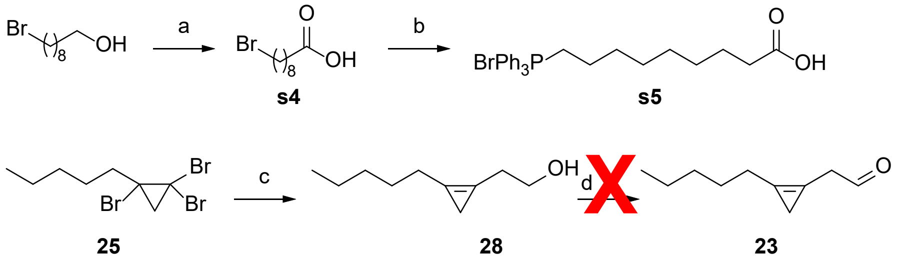

**Scheme S2. Synthetic route for phosphonium bromide s5 and attempted route towards cyclopropene aldehyde 23**. Reagents and conditions: a) TEMPO, BAIB, 2:1 MeCN / water, rt, 4h, 94% yield. b) $PPh_3$, MeCN, reflux, 18 h, quant. yield. c) *n*-Buli, $BF_3{\cdot}OEt_2$, oxirane, THF, -78 °C to 0 °C, 2 h, 49% yield. d) DMP, DCM, 0 °C to rt, 2 h. OR DMSO, $C_2O_2Cl_2$, DCM, $Et_3N$, -78 °C to rt, 2 h. OR DMSO, $Ac_2O$, DCM, rt, 2 h.

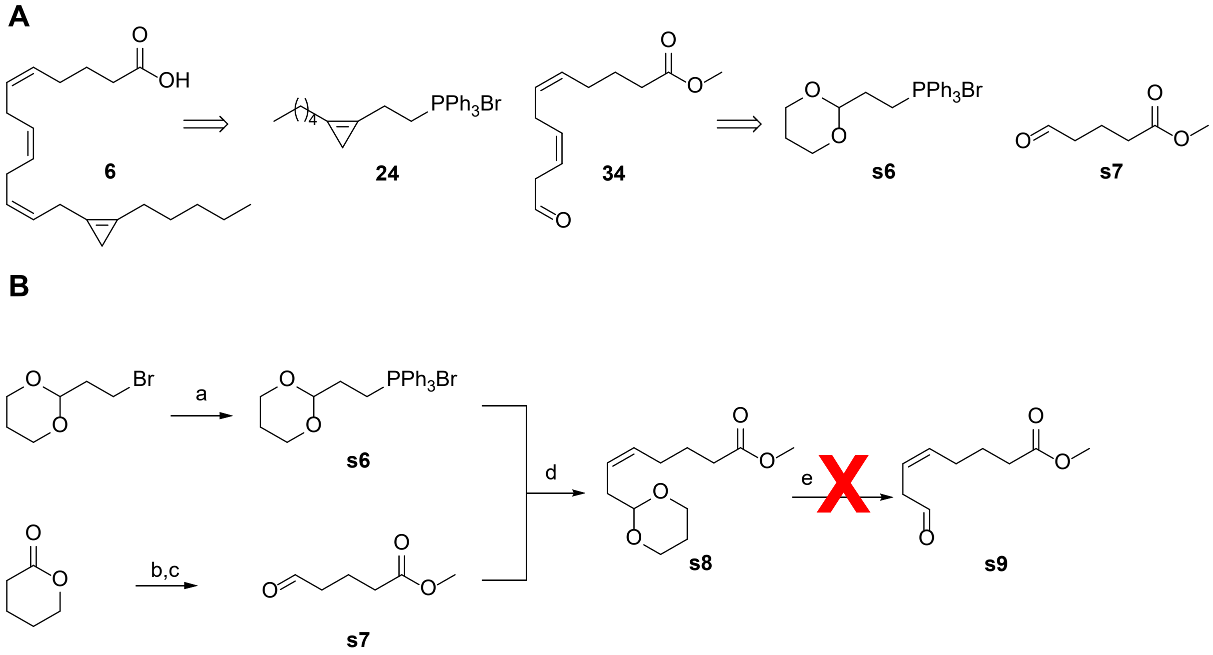

**Scheme S3. Synthetic strategy for PUFA with 3 or more double bonds**. A) Retrosynthetic analysis of fatty acid **6**. B) Reagents and conditions: a) $PPh_3$, MeCN, 82 °C, 18 h, 15% yield. b) $H_2SO_4$, MeOH, reflux, 18 h, 72% yield. c) DMP, DCM, 0 °C to rt, 2 h, 40% yield. d) NaHMDS, THF, -78 °C to rt, 2 h, 65% yield (95% Z-config.). e) HCl (aq.), THF, rt, 20 min OR AcOH, water, 85 °C, 4 h

Classified as Internal | Intern

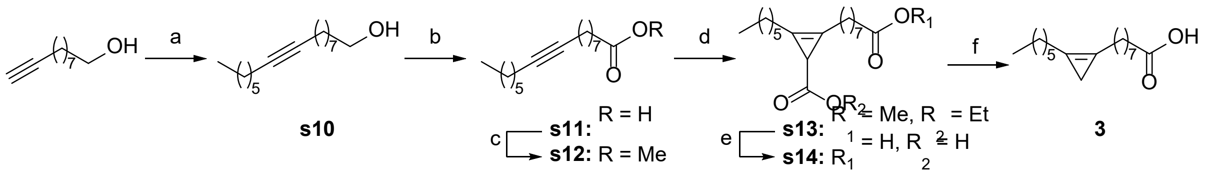

**Scheme S4. Synthesis route for cpPOA 3.** Reagents and conditions: a) *n*-BuLi, 1-bromohexane, NaI, 4:1 THF / HMPA, -78 °C to rt, 18 h, 44% yield. b) TEMPO, BAIB, 2:1 MeCN / water, 0 °C to rt, 2.5 h, 84% yield. c) $SOCl_2$, MeOH, 0 °C, 2.5 h, 88% yield. d) EDA, $Rh_2(OAc)_4$, DCM, 0 °C, 3.5 h, 76% yield. e) KOH, MeOH, reflux, 2.5 h, 97% yield. e) $SOCl_2$, $Et_2O$, 0 °C, 1 h, then $ZnCl_2$, DCM, rt, 3 h, then $NaBH_4$, NaOH, MeOH, -30 °C to rt, 1 h, 10% yield.

Classified as Internal | Intern

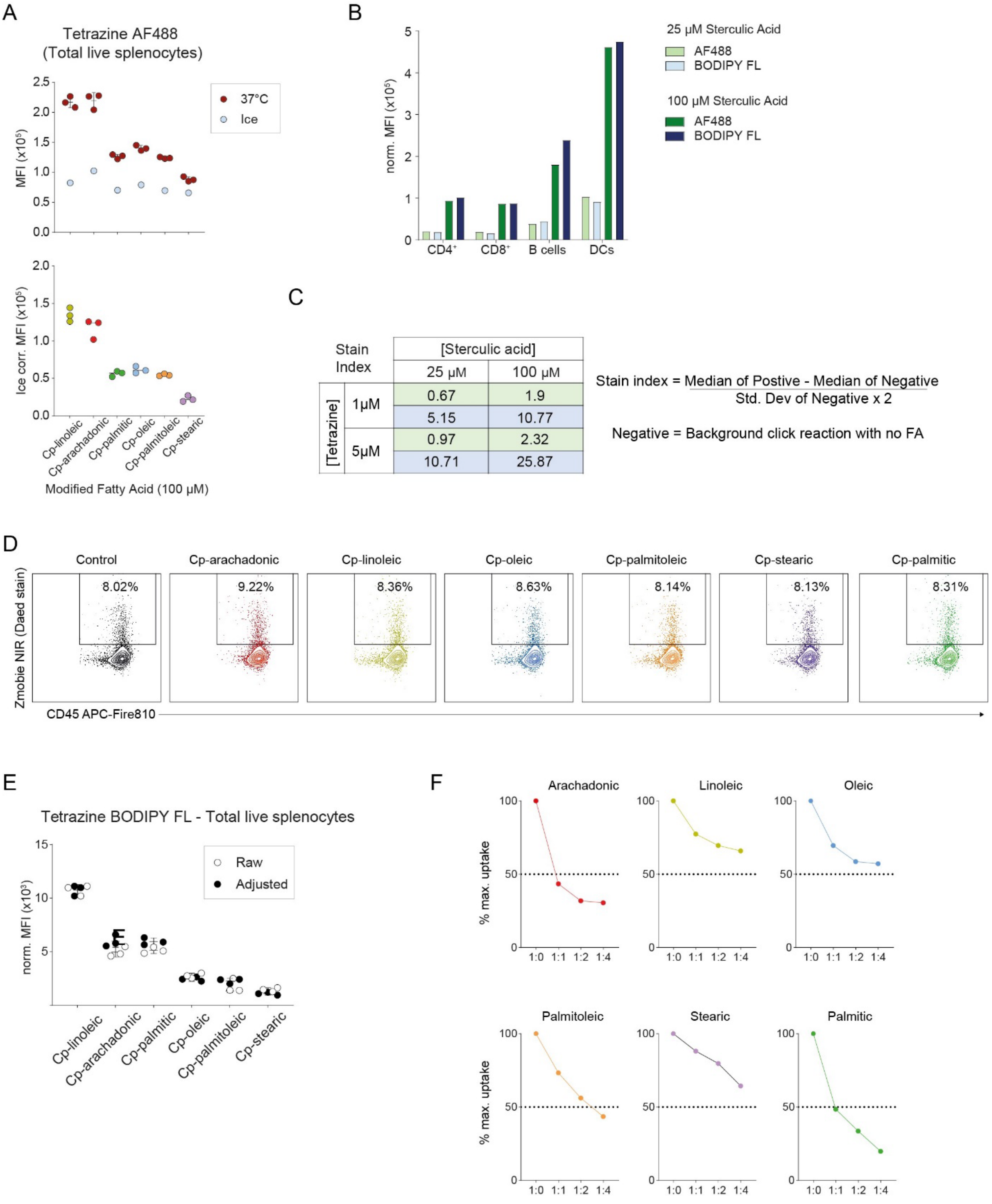

**Figure S1. Supporting Figure on the application of Cp-FAs in splenocytes.** A) Median fluorescent intensity of Tetrazine-AF488 (5 µM) following click reaction with 100 µM cpFA pulsed splenocytes (top) and normalized values after subtracting the MFI for splenocytes pulsed on ice (bottom). B) Ice-subtracted MFI of immune populations from the spleen, using 1 µM of either tetrazine-AF488 or tetrazine-BODIPY FL. C) The estimated stain index for tetrazine-AF488 (green) or tetrazine-BODIPY FL (blue), calculated as shown. D) Frequency of dead cells within the CD45+ single cell population, determined by amine dye exclusion and flow cytometry, following cpFA pulse (25 µM). E) Raw and adjusted data for each cpFA according to relative fluorescent intensity in a cell-free assay, set relative to linoleic acid. F) Competition assay showing the relative cpFA uptake with increasing concentration of the corresponding native FA.

Classified as Internal | Intern

### Supplementary Methods Immunology

#### Isolation of splenocytes

Spleens were isolated from naïve C57BL/6 mice bred under specific pathogen free (SPF) conditions. To obtain a single cell suspension, spleens were minced with scissors in a 2 mL Eppendorf and shaken at 37 °C, 200 rpm for 30 minutes in 1 mg/mL of Collagenase IV reconstituted in RPMI medium. Tissue digestion was stopped by adding 1 mL of cold PBS containing 0.5% BSA and 2 mM EDTA and placed on ice. The remaining homogenate was crushed and washed through a 40 µM strainer into a 50 mL tube, spun at 400g for 5 minutes and resuspended in 3 mL of ACK lysis buffer to remove red blood cells. After 5 minutes in lysis solution, 7 mL of cold PBS was added and cells were spun, washed through a second 40 µM strainer and counted for plating using a hemocytometer.

#### Preparation of fatty acid stocks

Solid cpFA stocks were first constituted in DMSO to make a 50 mM solution. Two 10x Step-wise dilutions were then done in RPMI medium containing 0.5% fatty acid free BSA to make a 5 mM and subsequent 0.5 mM working stock. The 0.5 mM stock was then diluted to the appropriate 1x concentration to be added to isolated splenocytes.

#### Uptake assay

Working FA stocks were first pre-warmed in a 37 °C water bath. 1-2 x $10^6$ splenocytes were plated in a 96-well V-bottom plate and kept on ice. Plated cells were washed twice in 200 µL of cold PBS. After washing, 100 µL of pre-diluted FA stocks (in the appropriate ratio with native FA for the competition experiment) were added directly to the splenocytes, which were immediately re-suspended by gentle pipetting. Resuspended cells were either placed in a 37 °C 5% $CO_2$ incubator or left on ice for 30 minutes, followed by addition of 100 µL cold PBS (0.5% BSA, 2mM EDTA) and spun at 400g for 5 minutes. Cells were washed an additional 2 times with cold PBS and left on ice for cell staining.

#### Cell staining and click-reaction for flow cytometry

Following FA uptake, splenocytes were stained in 50 µl of Zombie NIR viability dye diluted 500x and anti-XCR1 antibody in PBS for 15 minutes at room temperature. Cells were then washed 2x with 150 µL of cold PBS (0.5% BSA, 2mM EDTA), fixed with 2% methanol-free formaldehyde for 10 minutes at room temperature, and washed twice with PBS. Cells were subsequently permeabilized by incubation in 50 µL of PBS containing 0.1% saponin and 1% BSA for 15 minutes. Permeabilized cells were washed with PBS and the tetrazine-cyclopropene click reaction was performed by diluting the indicated tetrazine-fluorophore in PBS and incubating in 50 µL at room temperature for 1 hour in the dark. Cells were washed 2 times in PBS and either left in the fridge until staining could be performed or immediately stained with 50 µL of antibody cocktail for the remaining surface markers for 30 minutes, washed twice with PBS, and resuspended in 150 µL PBS for acquisition. A full list of antibodies used can be found in Supplementary Table S1. Fully stained samples were acquired on a Cytek 5-laser Aurora and unmixed using SpectroFlo v3. Analysis was done using FlowJo v10, GraphPad Prism and OMIQ.ai.

#### Fluorescence assay

cpFA stocks were diluted to 50 µM in a similar way to the uptake assay. In a black 96-well flat-bottom plate, 100 µL of tetrazine-BODIPY FL (2 µM) was added to 100 µL of cpFA stock (50 µM) and left to react at room temperature for 1 hour. The plate was scanned for fluorescence on a CLARIOstar plate reader (BMG LABTECH) with excitation/emission at 477-14/530- 40 and dichroic filter 497 showing the turn-on ratio between the sample and negative control (1 µM tetrazine-BODIPY FL), as an average of the three samples.

Classified as Internal | Intern

**Table S1.** Flow cytometry reagents.

| Marker | Conjugate | Dilution | Supplier | Cat. # |
|---|---|---|---|---|
| CD45 | APC-Fire810 | 1000x | BioLegend | 103174 |
| CD3 | PE/Dazzle594 | 1000x | BioLegend | 100246 |
| CD4 | APC | 1000x | ThermoFisher | 17-0042-83 |
| CD44 | BUV737 | 1000x | BD Biosciences | 612799 |
| CD8 | PE-Cy5 | 1000x | ThermoFisher | 15-0081-82 |
| CD19 | PE | 200x | ThermoFisher | 12-0191-82 |
| MHCII | AF700 | 400x | ThermoFisher | 56-5321-82 |
| CD11c | BV421 | 400x | BioLegend | 117330 |
| XCR1 | BV650 | 400x | BioLegend | 148220 |
| CD172a | BUV805 | 200x | BD Biosciences | 741997 |
| CD11b | PE-Cy7 | 10000x | ThermoFisher | 25-0112-82 |
| Ly6C | PerCP-Cy5.5 | 1000x | BioLegend | 128028 |

**Table S2.** Turn-on Ratio of cpFAs in fluorescence assay with tetrazine-BODIPY FL.

| cpFA | Turn-on Ratio | Relative ToR |
|---|---|---|
| Cp-palmitic acid | 4.9 | 0.56 |
| Cp-stearic acid | 1.8 | 1.58 |
| Cp-palmitoleic acid | 5.3 | 0.52 |
| Cp-oleic acid | 2.2 | 1.29 |
| Cp-linoleic acid | 2.8 | 1 |
| Cp-arachidonic acid | 3.2 | 0.88 |

Classified as Internal | Intern

### Supplementary Methods Chemistry

#### General information

All commercially available reagents and solvents were used as received unless stated otherwise. Anhydrous solvents were prepared by storage over activated 3 Å (only for MeOH) or 4 Å molecular sieves. Reaction progress was determined with thin layer chromatography (TLC) (Sigma, TLC Silica gel 60 F254) via UV visualisation (λ = 254 nm) and potassium permanganate stain (6 g $KMnO_4$ and 40 g, $K_2CO_3$ in 600 mL water). All column chromatography purifications were performed using solvents as received and silica gel (Macherey-Nagel, Kieselgel 60 M, 0.04 – 0.63 mm). Analytical LC-MS analysis was performed on a Finnigan Surveyor HPLC system (detection at 200-600 nm) with an analytical C18 column (Gemini, 50 x 4.6 mm, 3 µm particle size, Phenomenex) coupled to a Thermo LCQ Fleet Ion mass spectrometer (ESI+). Solvent system: A: water (gradient, 10-90%), B: $CH_3CN$ (gradient, 90-10%), C: (constant, 10%) 1% TFA (aq). Recorded data was interpreted and analysed using Xcalibur software. $^1$H and $^{13}$C NMR spectra were recorded using a Brüker AV-400 (400/101 MHz) or AV-500 (500/126 MHz). Chemical shift values are reported in ppm (δ) relative to the residual solvent signal of $CDCl_3$ (δ = 7.26 ppm for $^1$H and δ = 77.16 ppm for $^{13}$C NMR). Multiplicities are given as singlet (s), doublet (d), triplet (t), quartet (q), pentet (p), doublet of doublets (dd), broad singlet (br s) and multiplet (m). Recorded data was interpreted and analysed using MestReNova software.

#### General procedure A: Oxidation of alcohol to carboxylic acid

(Diacetoxyiodo)benzene (3 eq) and TEMPO (0.3 eq) were added to a solution of the alcohol (1. eq) in 2:1 MeCN / water (0.2 M) at 0 °C and stirred for 3 h at rt. The solution was quenched with sat. aq. $Na_2S_2O_3$, diluted with water and extracted with EtOAc (3x). The combined organic layers were washed with brine, dried over $MgSO_4$ and concentrated under reduced pressure. The crude product was purified by silica gel column chromatography to afford the carboxylic acid.

#### General procedure B: Hydrobromination of alkyne

In a slightly modified procedure from the literature[1], terminal alkyne (1 eq) was added dropwise to a diluted solution of $BBr_3$ (0.5 M in anh. DCM) (0.5 eq) at -78 °C under $N_2$ atmosphere and the reaction was slowly warmed to rt and stirred for 3 hours. The solution was cooled to 0 °C and acetic acid (30 eq) was added and stirred for another 1 h. The solution was neutralized carefully with sat. aq. $NaHCO_3$ and extracted with DCM (3x). The combined organic layers were washed with brine, dried over $MgSO_4$ and concentrated under reduced pressure (900 mbar). The crude product was purified by silica gel column chromatography (pentane) to afford 2-bromoalkene.

#### General procedure C: Cyclopropanation of 2-bromoalkene

In a slightly modified procedure from the literature[2], a sat. aq. NaOH solution (5 eq) was added dropwise to a solution of 2-bromo alkene (1 eq) and cetyltrimethylammonium bromide (0.1 eq) in bromoform (3 eq) and DCM (1.0 M) at 0 °C and stirred for 18 h. The solution was diluted with water and extracted with DCM (3x). The combined organic layers were washed with brine, dried over $MgSO_4$ and concentrated under reduced pressure. The crude product was purified by silica gel column chromatography (pentane) to afford the tribromocyclopropane.

Classified as Internal | Intern

**General procedure D: Substiution of tribromocyclopropane on diiodide**

In a slightly modified procedure from the literature[3], a solution of *n*-butyllithium (2.2 eq) was added dropwise to a solution of tribromocyclopropane (1 eq) in anhydrous THF (0.25 M) at -78 °C under $N_2$ atmosphere and was slowly warmed to 0 °C and stirred for 1 hour. A solution of diiodoalkane (2 eq) in anhydrous THF (1.0 M) was added and the reaction was stirred for 3 h at rt. The reaction was quenched with water, extracted with 1:1 $Et_2O$ / pentane (3x). The combined organic layers were washed with brine, dried over $MgSO_4$ and concentrated under reduced pressure. The crude product was purified by silica gel column chromatography (pentane) to afford the iodocyclopropene.

**General procedure E: Substitution of nitrile on iodocycloprpene**

In a slightly modified procedure from the literature[1], NaCN (2 eq) was added in one portion to a solution of iodocyclopropene (1 eq) in anhydrous DMSO (0.2 M) under $N_2$ atmosphere and stirred at 90 °C for 1 h. The reaction was diluted with water and extracted with $Et_2O$ (3x). The combined organic layers were washed with brine, dried over $MgSO_4$ and concentrated under reduced pressure. The crude product was purified by silica gel column chromatography ($Et_2O$/pentane 1:9) to afford the nitrilecyclopropene.

**General procedure F: Hydrolysis of nitrilecyclopropene**

In a slightly modified procedure from the literature[1], a sat. aq. NaOH solution (10 eq) was added to a solution of nitrilecyclopropene (1 eq) in 2:1 EtOH / water (0.2 M) and refluxed overnight. The reaction was acidified with aq. 1 M HCl and extracted with $Et_2O$ (3x). The combined organic layers were washed with brine, dried over $MgSO_4$ and concentrated under reduced pressure. The crude product was purified by silica gel column chromatography ($Et_2O$/pentane 1:9, 1% AcOH) to afford the cyclopropene fatty acid.

**General procedure G: Synthesis of phosphonium bromides**

In a slightly modified procedure from the literature[4], triphenylphosphine (1.1 eq) was added to a solution of alkylbromide (1 eq) in anh. MeCN at rt under $N_2$ atmosphere and the reaction mixture was stirred for 18 h at 100 °C. The solvents were removed under reduced pressure and the crude product was purified by silica gel column chromatography (DCM/MeOH 99:1 to 19:1) to afford the phosphonium bromide.

**General procedure H: Hydrolysis of cyclopropene fatty acid methyl ester**

An aq. 2 M NaOH solution (5 eq) was added to a solution of protected cyclopropene fatty acid methyl ester (1 eq) in 1:1 THF / EtOH (0.2 M) and stirred at 70 °C for 2 h. The reaction mixture was acidified with aq. 1 M HCl and extracted with $Et_2O$ (3x). The combined organic layers were washed with brine, dried over $MgSO_4$ and concentrated under reduced pressure. The crude product was purified by silica gel column chromatography ($Et_2O$/pentane 1:9, 1% AcOH) to afford the cyclopropene fatty acid.

Classified as Internal | Intern

**Synthetic procedures**

**2-bromohept-1-ene (10)**

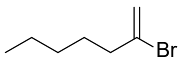

General procedure B was used with hept-1-yne (7.33 g, 76.22 mmol, 1.0 eq) as starting material to afford the title compound **10** (7.05 g, 39.81 mmol, 52% yield) as a colorless liquid. $^1$H NMR (400 MHz, $CDCl_3$) δ 5.55 (dt, *J* = 1.6, 1.2 Hz, 1H), 5.38 (d, *J* = 1.6 Hz, 1H), 2.41 (td, *J* = 7.4, 1.2 Hz, 2H), 1.65 – 1.48 (m, 2H), 1.39 – 1.15 (m, 4H), 0.90 (t, *J* = 7.0 Hz, 3H). Spectral data was in accordance with the literature.[9]

**1,1,2-tribromo-2-pentylcyclopropane (11)**

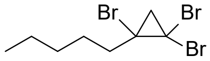

General procedure C was used with compound **10** (7.05 g, 39.81 mmol, 1.0 eq) as starting material to afford the title compound **11** (7.88 g, 22.57 mmol, 57% yield) as a colorless liquid. $^1$H NMR (400 MHz, $CDCl_3$) δ 2.13 – 1.99 (m, 2H), 1.98 (dd, *J* = 9.1, 0.8 Hz, 1H), 1.85 (d, *J* = 9.2 Hz, 1H), 1.82 – 1.58 (m, 2H), 1.48 – 1.29 (m, 4H), 1.10 – 0.84 (m, 3H). Spectral data was in accordance with the literature.[10]

**2-(2-pentylcycloprop-1-en-1-yl)ethan-1-ol (12)**

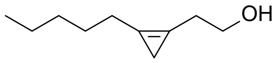

A solution of *n*-Buli (1.6 M in hexanes, 7.80 mL, 12.48 mmol, 2.2 eq) was added dropwise to a solution of compound **11** (1.98 g, 5.67 mmol, 1 eq) at -78 °C under $N_2$ atmosphere and stirred for 30 min. A solution of oxirane (2.5 – 3.3 M in THF, 4.54 mL, 11.35 mmol, 2 eq) and BF3·OEt (7.8 ml, 12.48 mmol, 2.2 eq) were added carefully and the reaction mixture was stirred for 15 min at -78 °C and for 1 h at 0 °C. The reaction was quenched with sat. aq. $NH_4Cl$ and extracted with $Et_2O$ (3x). The combined organic layers were washed with brine, dried over $MgSO_4$ and concentrated under reduced pressure. The crude product was purified by silica gel column chromatography ($Et_2O$ / pentane 1:9 to 1:4) to afford compound **12** (428 mg, 2.77 mmol, 49% yield) as a light yellow oil. **$^1$H NMR** (400 MHz, $CDCl_3$) δ 3.84 (t, *J* = 6.3 Hz, 2H), 2.68 (tt, *J* = 6.2, 1.6 Hz, 2H), 2.41 (tt, *J* = 7.3, 1.6 Hz, 2H), 1.61 – 1.52 (m, 2H), 1.50 (br s, 1H), 1.41 – 1.23 (m, 4H), 0.95 – 0.86 (m, 3H), 0.84 (s, 2H).

**1-(2-bromoethyl)-2-pentylcycloprop-1-ene (13)**

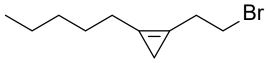

N-bromosuccinimide (506 mg, 3.28 mmol, 1 eq) and triphenylphosphine (1.03 mg, 3.94 mmol, 1.2 eq) were added in one portion to a solution of compound **12** (700 mg, 3.94 mmol, 1.2 eq) in anh. DCM at 0 °C under $N_2$ atmosphere and stirred at rt for 1 h. The reaction mixture was quenched with sat. aq. $NaHCO_3$ and extracted with DCM (3x). The combined organic layers were washed with brine, dried over $MgSO_4$ and concentrated under reduced pressure. The crude product was purified by silica gel column chromatography (pentane) to afford compound **13** (417 mg, 1.92 mmol, 59% yield) as a light yellow oil. **$^1$H NMR** (400 MHz, $CDCl_3$) δ 3.54 (t, *J* = 7.0 Hz, 2H), 2.99 (tt, *J* = 7.1, 1.6 Hz, 2H), 2.42 (tt, *J* =

Classified as Internal | Intern

7.3, 1.5 Hz, 2H), 1.62 – 1.51 (m, 2H), 1.39 – 1.21 (m, 4H), 0.93 – 0.87 (m, 3H), 0.86 (s, 2H). **$^{13}$C NMR** (101 MHz, $CDCl_3$) δ 112.9, 106.7, 31.7, 30.2, 30.2, 27.0, 26.2, 22.6, 14.2, 7.6.

**(2-(2-pentylcycloprop-1-en-1-yl)ethyl)triphenylphosphonium bromide (7)**

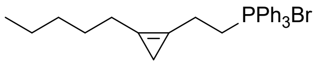

General procedure G was used with compound **13** (49 mg, 0.23 mmol, 1.0 eq) as starting material to afford the title compound **7** (89 mg, 0.19 mmol, 82% yield) as a light yellow oil. **$^{1}$H NMR** (400 MHz, $CDCl_3$) δ 7.99 – 7.64 (m, 15H), 3.96 (dt, *J* = 12.1, 7.0 Hz, 2H), 3.00 (dtt, *J* = 17.4, 7.1, 1.5 Hz, 2H), 2.23 (tt, *J* = 7.3, 1.6 Hz, 2H), 1.47 – 1.35 (m, 2H), 1.35 – 1.14 (m, 4H), 0.85 (t, *J* = 7.0 Hz, 3H), 0.60 (s, 2H). **$^{13}$C NMR (**101 MHz, $CDCl_3$) δ 135.3, 135.2, 133.6, 133.5, 130.6, 130.5, 118.2, 117.3, 113.5, 105.7, 105.6, 31.4, 26.7, 25.5, 22.3, 21.5, 21.0, 19.5, 19.5, 14.0, 7.9.

**9-((tert-butyldimethylsilyl)oxy)nonan-1-ol (14)**

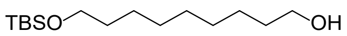

Imidazole (510 mg, 7.49 mmol, 1.2 eq) and TBSCl (940 mg, 6.24 mmol, 1 eq) were added to a solution of nonane-1,9-diol (2.0 g, 12.48 mmol, 2 eq) in anh. DCM at rt under $N_2$ atmosphere and stirred for 3 h. The reaction mixture was diluted with water and extracted with DCM (3x). The combined organic layers were washed with brine, dried over $MgSO_4$ and concentrated under reduced pressure. The crude product was purified by silica gel column chromatography ($Et_2O$ / pentane 1:4 to 1:2) to afford compound **14** (1.10 g, 4.0 mmol, 64% yield) as a clear oil. **$^{1}$H NMR** (400 MHz, $CDCl_3$) δ 3.64 (td, *J* = 6.7, 5.2 Hz, 2H), 3.59 (t, *J* = 6.6 Hz, 2H), 1.64 – 1.44 (m, 4H), 1.40 – 1.27 (m, 10H), 0.89 (s, 9H), 0.04 (s, 6H). $^{1}$H NMR (400 MHz, $CDCl_3$) δ 3.63 (t, *J* = 6.6 Hz, 2H), 3.59 (t, *J* = 6.6 Hz, 2H), 1.61 – 1.44 (m, 4H), 1.37 – 1.22 (m, 10H), 0.89 (s, 9H), 0.04 (s, 6H). **$^{13}$C NMR** (101 MHz, $CDCl_3$) δ 63.5, 63.2, 33.0, 32.9, 29.7, 29.5, 26.1, 25.9, 25.9, 18.5, -5.1. Spectral data was in accordance with the literature.[11]

**9-((tert-butyldimethylsilyl)oxy)nonanoic acid (15)**

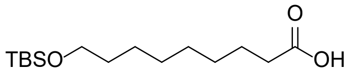

General procedure A was used with compound **14** (1.16 g, 4.02 mmol, 1.0 eq) as starting material, the crude product was purified by silica gel column chromatography ($Et_2O$ / pentane 1:4, 1% AcOH) to afford compound **15** (1.07 g, 3.70 mmol, 92% yield) as a light yellow oil. **$^{1}$H NMR** (400 MHz, $CDCl_3$) δ 3.59 (t, *J* = 6.6 Hz, 2H), 2.34 (t, *J* = 7.5 Hz, 2H), 1.63 (p, *J* = 7.5 Hz, 2H), 1.54 – 1.45 (m, 2H), 1.38 – 1.24 (m, 8H), 0.89 (s, 9H), 0.04 (s, 6H). **$^{13}$C NMR** (101 MHz, $CDCl_3$) δ 180.3, 63.4, 34.2, 33.0, 29.4, 29.1, 26.1, 25.9, 24.8, 18.5, -5.1. Spectral data was in accordance with the literature.[11]

**methyl 9-((tert-butyldimethylsilyl)oxy)nonanoate (16)**

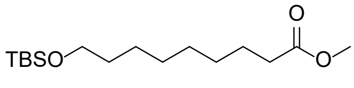

Classified as Internal | Intern

Methyl iodide (0.43 mL, 6.93 mmol, 2 eq) and $K_2CO_3$ (1.44 g, 10.4 mmol, 3 eq) were added to a solution of compound **15** (1.0 g, 3.47 mmol, 1 eq) in anh. DMF at rt under $N_2$ atmosphere and stirred for 18 h. The reaction mixture was diluted (10x) with sat. aq. $NaHCO_3$ and extracted with $Et_2O$ (3x). The combined organic layers were washed with brine, dried over $MgSO_4$ and solvents removed under reduced pressure. The crude product was purified by silica gel column chromatography ($Et_2O$ / pentane 1:9) to afford compound **16** (898 mg, 2.97 mmol, 86% yield). **$^1$H NMR** (400 MHz, $CDCl_3$) δ 3.66 (s, 3H), 3.59 (t, *J* = 6.6 Hz, 2H), 2.30 (t, *J* = 7.6 Hz, 2H), 1.70 – 1.56 (m, 2H), 1.55 – 1.43 (m, 2H), 1.35 – 1.23 (m, 8H), 0.89 (s, 9H), 0.04 (s, 6H). **$^{13}$C NMR** (101 MHz, $CDCl_3$) δ 174.5, 63.4, 51.6, 34.2, 33.0, 29.4, 29.2, 26.1, 25.9, 25.1, 18.5, -5.1. Spectral data was in accordance with the literature.[12]

**methyl 9-hydroxynonanoate (17)**

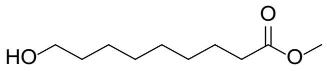

A solution of TBAF (1.0 M in THF, 8.73 mL, 8.73 mmol, 3 eq) was added dropwise to a solution of compound **16** (880 mg, 2.91 mmol, 1 eq) in anh. THF (0.3 M) at 0 °C under $N_2$ atmosphere and stirred for 5 h at rt. The reaction mixture was diluted with aq. 1 M HCl and extracted with DCM (3x). The combined organic layers were washed with brine, dried over $MgSO_4$ and solvents removed under reduced pressure. The crude product was purified by silica gel column chromatography (EtOAc/pentane 1:4 to 1:1) to afford compound **17** (512 mg, 2.72 mmol, 93% yield) **$^1$H NMR** (400 MHz, $CDCl_3$) δ 3.65 (s, 3H), 3.61 (t, *J* = 6.6 Hz, 2H), 2.28 (t, *J* = 7.5 Hz, 2H), 1.68 – 1.47 (m, 4H), 1.38 – 1.23 (m, 8H). **$^{13}$C NMR** (101 MHz, $CDCl_3$) δ 174.5, 63.1, 51.6, 34.2, 32.8, 29.3, 29.1, 25.8, 25.0. Spectral data was in accordance with the literature. [13]

**methyl 9-oxononanoate (8)**

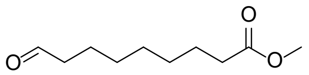

Dess-Martin periodinane (1.41 g, 3.32 mmol, 1.3 eq) was added to a solution of compound **17** (481 mg, 2.55 mmol, 1 eq) in anh. DCM (0.2 M) at 0 °C under $N_2$ atmosphere and stirred for 1 h at rt. The reaction mixture was quenched with sat. aq. $NaHCO_3$ and extracted with $Et_2O$ (3x). The combined organic layers were washed with brine, dried over $MgSO_4$ and solvents removed under reduced pressure. The crude product was purified by silica gel column chromatography ($Et_2O$ / pentane 1:4) to afford compound **8** (372 mg, 2.0 mmol, 78% yield) as a light yellow oil. **$^1$H NMR** (400 MHz, $CDCl_3$) δ 9.75 (t, *J* = 1.6, 1H), 3.65 (s, 3H), 2.41 (td, *J* = 7.3, 1.8 Hz, 2H), 2.29 (t, *J* = 7.5 Hz, 2H), 1.70 – 1.52 (m, 4H), 1.39 – 1.22 (m, 6H). **$^{13}$C NMR** (101 MHz, $CDCl_3$) δ 202.9, 174.3, 51.6, 43.9, 34.1, 29.1, 29.0, 29.0, 24.9, 22.1. Spectral data was in accordance with the literature.[13]

**methyl (Z)-11-(2-pentylcycloprop-1-en-1-yl)undec-9-enoate (18)**

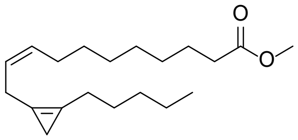

A solution of NaHMDS (1 M, 0.27 mL, 0.27 mmol, 1 eq) was added dropwise to a solution of compound **7** (128 mg, 0.12 mmol, 1 eq) in anh. THF (0.25 M) at -78 °C under $N_2$ atmopshere and stirred for 1 h. A

Classified as Internal | Intern

solution of compound **8** (60 mg, 0.32 mmol, 1.2 eq) in anh. THF (0.5 M) was added dropwise to the orange brown suspension and stirred for 1.5 h. The reaction mixture was quenched with sat. aq. $NH_4Cl$ and extracted with $Et_2O$ (3x). The combined organic layers were washed with brine, dried over $MgSO_4$ and solvents removed under reduced pressure. The crude product was purified by silica gel column chromatography ($Et_2O$ / pentane 1:99 to 1:49) to afford compound **18** (36 mg, 0.12 mmol, 44% yield) as a clear colorless oil. **$^1$H NMR** (500 MHz, $CDCl_3$) δ 5.55 – 5.41 (m, 2H), 3.66 (s, 3H), 3.12 (d, *J* = 6.3 Hz, 2H), 2.37 (tt, *J* = 7.3, 1.6 Hz, 2H), 2.29 (t, *J* = 7.6 Hz, 2H), 2.05 (q, *J* = 7.1, 6.7 Hz, 2H), 1.66 – 1.56 (m, 2H), 1.57 – 1.49 (m, 2H), 1.39 – 1.24 (m, 12H), 0.89 (t, *J* = 6.9 Hz, 3H), 0.81 (s, 2H). **$^{13}$C NMR** (126 MHz, $CDCl_3$) δ 174.4, 131.3, 125.3, 110.1, 108.1, 51.6, 34.2, 31.7, 29.6, 29.3, 29.2, 29.2, 27.3, 27.3, 26.1, 25.1, 24.8, 22.6, 14.2, 7.8.

**(Z)-11-(2-pentylcycloprop-1-en-1-yl)undec-9-enoic acid (5)**

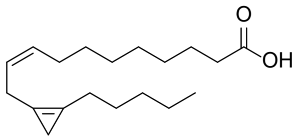

General procedure H was used with compound **18** (36 mg, 0.12 mmol, 1.0 eq) as starting material to afford the title compound **5** (25 mg, 85 µmol, 73% yield) as a light yellow oil. **$^1$H NMR** (400 MHz, $CDCl_3$) δ 11.02 (br s, 1H), 5.66 – 5.31 (m, 2H), 3.13 (d, *J* = 6.4 Hz, 2H), 2.43 – 2.29 (m, 4H), 2.12 – 2.00 (m, 2H), 1.70 – 1.49 (m, 4H), 1.43 – 1.23 (m, 12H), 0.92 – 0.87 (m, 3H), 0.81 (s, 2H). **$^{13}$C NMR** (101 MHz, $CDCl_3$) δ 180.4, 131.3, 125.3, 110.1, 108.1, 34.2, 31.8, 29.6, 29.3, 29.2, 29.2, 27.3, 27.3, 26.1, 24.8, 24.8, 22.6, 14.2, 7.8.

**methyl 7-hydroxyhept-5-ynoate (19)**

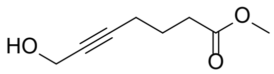

In a slightly modified procedure from the literature[14], a solution of EtMgBr (3.0 M in $Et_2O$, 45.5 mL, 136.5 mmol, 3 eq) was added to a solution of hex-5-ynoic acid (5.10 g, 45.5 mmol, 1 eq) in anh. THF (0.2 M) at 0 °C under $N_2$ atmosphere and stirred for 1 h slowly warming to rt. Then, paraformaldehyde (2.73 g, 91.0 mmol, 2 eq) was added portionwise to the grey suspension and the reaction mixture was stirred at 75 °C for 18 h. The reaction mixture was quenched with aq. 1 M HCl and extracted with DCM (3x). The combined organic layers were washed with brine, dried over $MgSO_4$ and solvents removed under reduced pressure. Thionyl chloride (6.60 mL, 91.0 mmol, 2 eq) was added carefully to a solution of the crude product in anh. MeOH (0.2 M) at 0 °C under $N_2$ atmosphere and stirred for 1 h at rt. The reaction mixture was concentrated under reduced pressure and the crude product was purified by silica gel column chromatography ($Et_2O$ / pentane 1:2) to afford compound **19** (3.60 g, 23.1 mmol, 51% yield) as a clear oil. **$^1$H NMR** (400 MHz, $CDCl_3$) δ 4.23 (t, *J* = 2.2 Hz, 2H), 3.67 (s, 3H), 2.43 (t, *J* = 7.4 Hz, 2H), 2.28 (tt, *J* = 6.9, 2.2 Hz, 2H), 1.82 (p, *J* = 7.1 Hz, 2H). **$^{13}$C NMR** (101 MHz, $CDCl_3$) δ 173.8, 85.1, 79.4, 51.8, 51.4, 32.9, 23.8, 18.3. Spectral data was in accordance with the literature.[15]

**methyl 7-bromohept-5-ynoate (20)**

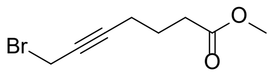

Classified as Internal | Intern

N-bromosuccinimide (1.16 g, 6.53 mmol, 1.2 eq) and triphenylphosphine (1.71 g, 6.53 mmol, 1.2 eq) were added to a solution of compound **19** (855 mg, 5.44 mmol, 1 eq) in anh. DCM (0.2 M) at 0 °C under $N_2$ atmosphere and stirred for 1 h at rt. The reaction mixture was quenched with sat. aq. $NaHCO_3$ and extracted with DCM (3x). The combined organic layers were washed with brine, dried over $MgSO_4$ and solvents removed under reduced pressure. The crude product was purified by silica gel column chromatography ($Et_2O$ / pentane 1:19) to afford compound **20** (980 mg, 4.47 mmol, 82% yield) as a light yellow oil. **$^1$H NMR** (400 MHz, $CDCl_3$) δ 3.90 (t, *J* = 2.4 Hz, 2H), 3.67 (s, 3H), 2.42 (t, *J* = 7.4 Hz, 2H), 2.31 (tt, *J* = 6.9, 2.4 Hz, 2H), 1.82 (p, *J* = 7.1 Hz, 2H). **$^{13}$C NMR** (101 MHz, $CDCl_3$) δ 173.6, 86.8, 76.3, 51.7, 32.8, 23.6, 18.5, 15.5. Spectral data was in accordance with the literature.[15]

**methyl 11-hydroxyundeca-5,8-diynoate (10)**

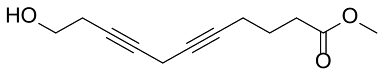

Copper(I)iodide (447 mg, 2.92 mmol, 2 eq), Sodium iodide (438 mg, 2.92 mmol, 2 eq) and $K_2CO_3$ (404 mg, 2.92 mmol, 2 eq) were added to a solution of but-3-yn-1-ol (0.22 mL, 2.92 mmol, 2 eq) in anh. DMF (0.2 M) at rt under $N_2$ atmosphere and stirred for 30 min at rt. Then, a solution of compound **20** (320 mg, 1.46 mmol, 1 eq) in anh. DMF (1.0 M) was added dropwise to the reaction mixture and stirred for 18 h. The suspension was diluted with sat. aq. $NH_4Cl$ and extracted with $Et_2O$ (3x). The combined organic layers were washed with sat. aq. $Na_2S_2O_3$ and brine, dried over $MgSO_4$ and solvents removed under reduced pressure. The crude product was purified by silica gel column chromatography ($Et_2O$ / pentane 1:1) to afford compound **10** (220 mg, 1.06 mmol, 72% yield) as a light yellow oil. **$^1$H NMR** (400 MHz, $CDCl_3$) δ 3.67 (t, *J* = 6.3 Hz, 2H), 3.65 (s, 3H), 3.10 (p, *J* = 2.4 Hz, 2H), 2.46 – 2.36 (m, 4H), 2.21 (tt, *J* = 6.9, 2.4 Hz, 2H), 2.17 – 2.04 (m, 1H), 1.79 (p, *J* = 7.1 Hz, 2H). **$^{13}$C NMR** (101 MHz, $CDCl_3$) δ 173.8, 79.5, 77.1, 76.6, 75.2, 61.2, 51.7, 33.0, 23.9, 23.2, 18.3, 9.8. Spectral data was in accordance with the literature.[15]

**methyl (5Z,8Z)-11-hydroxyundeca-5,8-dienoate (21)**

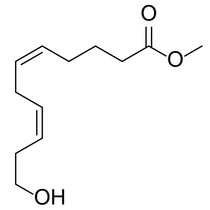

In a slightly modified procedure from the literature[16], a solution of $NaBH_4$ (111 mg, 2.94 mmol, 2 eq) in degassed anh. MeOH (1.5 M) was added to a solution of Nickel(II) acetate hydrate (658 mg, 2.64 mmol, 1.8 eq) in MeOH (1.5 M) at rt under $N_2$ atmosphere, the solution turned dark black immediately and was purged with $H_2$ for 5 min. Ethylenediamine (0.18 ml, 2.64 mmol, 1.8 eq) and a solution of compound **10** (306 mg, 1.47 mmol, 1 eq) in MeOH (1.5 M) were added and the reaction mixture was stirred under positive $H_2$ pressure for 2 h at rt and monitored using LCMS. The suspension was diluted with $Et_2O$, filtered over celite, washed with brine and dried over $MgSO_4$. The crude product was purified by silica gel column chromatography (Et2O / pentane 1:1) to afford a mixture of under / over reduced or even cis / trans products (200 mg, 0.94 mmol, 64 % yield). This was purified by HPLC (Solvent system: A: water (gradient, 10-90%), B: $CH_3CN$ (gradient, 90-10%), C: (constant, 10%) 1% TFA (aq.) to afford compound **21** (84 mg, 0.40 mmol, 27% yield) as a clear colorless oil. **$^1$H NMR** (400 MHz, $CDCl_3$) δ 5.52 – 5.25 (m, 4H), 3.62 (s, 3H), 3.59 (t, *J* = 6.6 Hz, 2H), 2.81 – 2.72 (m, 2H), 2.36 – 2.24 (m, 4H), 2.13 (s, 1H), 2.10 – 2.01 (m, 2H), 1.65 (p, *J* = 7.5 Hz, 2H). **$^{13}$C NMR** (101 MHz, $CDCl_3$) δ 174.3, 130.8,

Classified as Internal | Intern

129.0, 128.8, 125.7, 62.2, 51.6, 33.4, 30.9, 26.6, 25.8, 24.7. Spectral data was in accordance with the literature.[17]

**methyl (5Z,8Z)-11-oxoundeca-5,8-dienoate (9)**

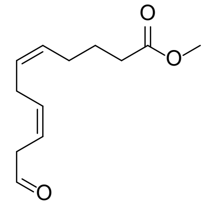

Dess-Martin periodinane (201 mg, 0.47 mmol, 1.2 eq) was added to a solution of compound **21** (84 mg, mmol, 1 eq) in anh. DCM (0.2 M) at 0 °C under $N_2$ atmosphere and stirred for 1 h at rt. The reaction mixture was quenched with sat. aq. $NaHCO_3$ and extracted with $Et_2O$ (3x). The combined organic layers were washed with brine, dried over $MgSO_4$ and solvents removed under reduced pressure. The crude product was purified by silica gel column chromatography ($Et_2O$ / pentane 1:2) to afford compound **22** (31 mg, 0.15 mmol, 37% yield) as a clear oil. **$^1$H NMR** (400 MHz, $CDCl_3$) δ 9.66 (t, *J* = 1.9 Hz, 1H), 5.72 – 5.49 (m, 2H), 5.46 – 5.30 (m, 2H), 3.65 (s, 3H), 3.26 – 3.16 (m, 2H), 2.80 – 2.73 (m, 2H), 2.30 (t, *J* = 7.4 Hz, 2H), 2.13 – 2.03 (m, 2H), 1.74 – 1.62 (m, 2H). **$^{13}$C NMR** (101 MHz, $CDCl_3$) δ 199.5, 174.1, 133.3, 129.7, 127.9, 118.7, 51.6, 42.6, 33.5, 26.7, 26.0, 24.8. Spectral data was in accordance with the literature.[18]

**methyl (5Z,8Z,11Z)-13-(2-pentylcycloprop-1-en-1-yl)trideca-5,8,11-trienoate (22)**

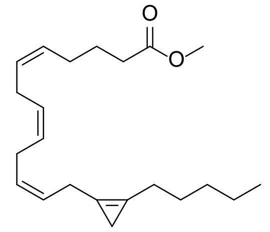

A solution of NaHMDS (1 M, 0.29 mL, 0.29 mmol, 2 eq) was added dropwise to a solution of compound **7** (140 mg, 0.29 mmol, 2 eq) in anh. THF (0.25 M) at -78 °C under $N_2$ atmopshere and stirred for 1 h. A solution of compound **9** (31 mg, 0.15 mmol, 1 eq) in anh. THF (0.5 M) was added dropwise to the orange brown suspension and stirred for 1.5 h. The reaction mixture was quenched with sat. aq. $NH_4Cl$ and extracted with $Et_2O$ (3x). The combined organic layers were washed with brine, dried over $MgSO_4$ and solvents removed under reduced pressure. The crude product was purified by silica gel column chromatography ($Et_2O$ / pentane 1:99 to 1:49) to afford compound **22** (26 mg, 78 µmol, 54% yield) as a clear colorless oil. **$^1$H NMR** (500 MHz, $CDCl_3$) δ 5.60 – 5.40 (m, 2H), 5.42 – 5.29 (m, 4H), 3.66 (s, 3H), 3.17 (d, *J* = 7.0 Hz, 2H), 2.85 (t, *J* = 6.5 Hz, 2H), 2.80 (t, *J* = 5.8 Hz, 2H), 2.38 (t, *J* = 7.3 Hz, 2H), 2.32 (t, *J* = 7.5 Hz, 2H), 2.11 (q, *J* = 7.0 Hz, 2H), 1.70 (p, *J* = 7.5 Hz, 2H), 1.54 (p, *J* = 7.0 Hz, 2H), 1.37 – 1.26 (m, 4H), 0.89 (t, *J* = 6.7 Hz, 3H), 0.82 (s, 2H). **$^{13}$C NMR** (126 MHz, $CDCl_3$) δ 174.2, 129.2, 129.1, 129.0, 128.4, 128.1, 125.8, 110.4, 107.8, 51.6, 33.6, 31.7, 27.2, 26.7, 26.0, 25.8, 25.7, 24.9, 24.8, 22.6, 14.2, 7.8.

**(5Z,8Z,11Z)-13-(2-pentylcycloprop-1-en-1-yl)trideca-5,8,11-trienoic acid (6)**

Classified as Internal | Intern

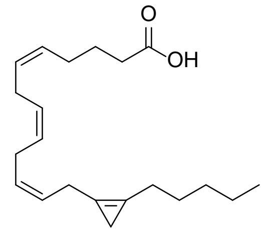

General procedure H was used with compound **22** (26 mg, 78 µmol, 1.0 eq) as starting material to afford the title compound **6** (25 mg, 78 µmol, quantitative yield) as a clear colorless oil. **$^1$H NMR** (500 MHz, $CDCl_3$) δ 11.24 (br s, 1H), 5.60 – 5.41 (m, 2H), 5.44 – 5.30 (m, 4H), 3.17 (d, *J* = 7.0 Hz, 2H), 2.85 (t, *J* = 6.6 Hz, 2H), 2.80 (t, *J* = 6.1 Hz, 2H), 2.38 (q, *J* = 8.2 Hz, 4H), 2.13 (q, *J* = 7.1 Hz, 2H), 1.72 (p, *J* = 7.5 Hz, 2H), 1.55 (p, *J* = 7.2 Hz, 2H), 1.37 – 1.25 (m, 4H), 0.89 (t, *J* = 6.5 Hz, 3H), 0.83 (s, 2H). **$^{13}$C NMR** (126 MHz, $CDCl_3$) δ 180.0, 129.2, 128.9, 128.4, 128.2, 125.8, 110.4, 107.9, 33.5, 31.8, 27.3, 26.6, 26.1, 25.8, 25.8, 24.8, 24.6, 22.6, 14.2, 7.9.

**8-(2-hexylcycloprop-1-en-1-yl)octanoic acid (3)**

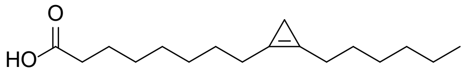

In a slightly modified procedure from the literature[7], thionylchloride (0.48 mL, 6.62 mmol, 8.1 eq) was added to a solution of compound **s14** (254 mg, 0.819 mmol, 1.0 eq) in anhydrous $Et_2O$ (0.2 M) under $N_2$ atmosphere at 0 °C and stirred for 1 h. The reaction mixture was concentrated under reduced pressure and a solution of the crude acid chloride in anhydrous DCM (0.2 M) was added to anhydrous $ZnCl_2$ (221 mg, 1.622 mmol, 2.0 eq) under $N_2$ atmosphere. The suspension was stirred for 1 h and periodically flushed with $N_2$ gas, after which additional anhydrous $ZnCl_2$ (271 mg, 1.988 mmol, 2.4 eq) was added and stirred for another 2 hours. Then, methanol (0.035 mL, 0.87 mmol, 1.06 eq) was added dropwise and stirred for 30 min. The reaction mixture was added to a solution of $NaBH_4$ (167 mg, 4.41 mmol, 5.4 eq) and NaOH (74 mg, 1.85 mmol, 2.25 eq) in anhydrous meOH (0.1 M) at -30 °C and stirred for 1 h gradually warming to rt. The reaction was quenched with water, acidified with aq. 1 M HCl and extracted with DCM (3x). The combined organic layers were washed with water, sat. aq. $NaHCO_3$ and brine, dried over $MgSO_4$ and concentrated under reduced pressure. The crude product was purified by silica gel column chromatography ($Et_2O$ / pentane 1:9) to afford compound **3** (21 mg, 0.079 mmol, 10% yield). **$^1$H NMR** (400 MHz, $CDCl_3$) δ 2.37 (t, *J* = 7.2Hz, 4H), 2.35 (t, *J* = 7.68Hz, 2H), 1.71 – 1.58 (m, 2H), 1.59 – 1.46 (m, 4H), 1.42 – 1.15 (m, 12H), 0.88 (t, *J* = 6.8 Hz, 3H), 0.76 (s, 2H). **$^{13}$C NMR** (101 MHz, $CDCl_3$) δ 179.7, 109.5, 109.2, 34.0, 31.7, 29.2, 29.1, 29.0, 29.0, 27.4, 27.3, 26.1, 26.0, 24.7, 22.6, 14.1, 7.4.

**1,1,2-tribromo-2-methylcyclopropane (25)**

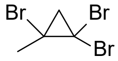

General procedure C was used with 2-bromoprop-1-ene **23** (4.50 g, 37.20 mmol, 1.0 eq) as starting material to afford the title compound **25** (5.10 g, 17.42 mmol, 47% yield) as a colorless liquid. **$^1$H NMR** (300 MHz, $CDCl_3$) δ 2.08 (s, 3H), 1.98 (d, *J* = 9.2 Hz, 1H), 1.84 (d, *J* = 9.2 Hz, 1H). **$^{13}$C NMR** (75 MHz, $CDCl_3$) δ 39.7, 38.6, 33.3, 30.0. Spectral data was in accordance with the literature.[8]

**1-(12-iodododecyl)-2-methylcycloprop-1-ene (27)**

Classified as Internal | Intern

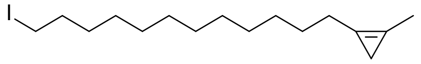

General procedure D was used with compound **25** (410 mg, 1.40 mmol, 1.0 eq) and 1,12-diiodododecane (1.48 g, 3.50 mmol, 2.5 eq) as starting materials to afford the title compound **27** (134 mg, 0.38 mmol, 27 % yield). **$^1$H NMR** (300 MHz, $CDCl_3$) δ 3.18 (t, *J* = 7.1 Hz, 2H), 2.36 (tq, *J* = 7.3, 1.7 Hz, 2H), 2.03 (t, *J* = 1.6 Hz, 3H), 1.82 (p, *J* = 7.1 Hz, 2H), 1.58 – 1.48 (m, 2H), 1.43 – 1.24 (m, 16H), 0.77 (s, 2H). **$^{13}$C NMR** (75 MHz, $CDCl_3$) δ 110.2, 105.3, 33.7, 30.7, 29.7, 29.7, 29.6, 29.5, 28.7, 27.5, 26.1, 11.7, 8.3, 7.5.

**13-(2-methylcycloprop-1-en-1-yl)tridecanenitrile (30)**

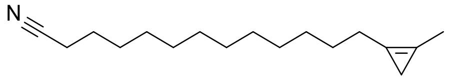

General procedure E was used with compound **27** (134 mg, 0.38 mmol, 1.0 eq) as starting material to afford the title compound **30** (76 mg, 0.31 mmol, 80 % yield). **$^1$H NMR** (300 MHz, $CDCl_3$) δ 2.40 – 2.32 (m, 2H), 2.33 (t, *J* = 7.1 Hz, 2H), 2.03 (t, *J* = 1.6 Hz, 3H), 1.73 – 1.57 (m, 2H), 1.59 – 1.38 (m, 4H), 1.37 – 1.18 (m, 14H), 0.77 (s, 2H). **$^{13}$C NMR** (75 MHz, $CDCl_3$) δ 120.0, 110.2, 105.3, 29.7, 29.6, 29.5, 29.5, 29.4, 28.9, 28.8, 27.5, 26.1, 25.5, 17.3, 11.7, 8.2.

**13-(2-methylcycloprop-1-en-1-yl)tridecanoic acid (1)**

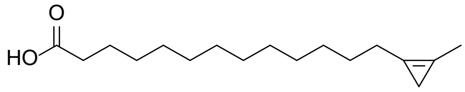

General procedure F was used with compound **30** (76 mg, 0.31 mmol, 1.0 eq) as starting material to afford the title compound **1** (30 mg, 0.11 mmol, 37% yield). **$^1$H NMR** (300 MHz, $CDCl_3$) δ 2.41 – 2.29 (m, 4H), 2.03 (t, *J* = 1.5 Hz 3H), 1.63 (p, *J* = 7.4 Hz, 2H), 1.59 – 1.46 (m, 2H), 1.38 – 1.16 (m, 16H), 0.77 (s, 2H). $^{13}$C NMR (75 MHz, $CDCl_3$) δ 179.8, 110.1, 105.2, 34.0, 29.6, 29.6, 29.6, 29.4 29.4, 29.4, 29.3, 29.1, 27.4, 26.0, 24.7, 11.5, 8.1.

**1-(14-iodotetradecyl)-2-methylcycloprop-1-ene (28)**

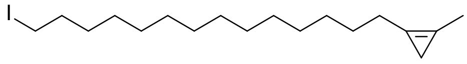

General procedure D was used with compound **25** (488 mg, 1.67 mmol, 1.0 eq) and 1,14-diiodotetradecane (1.58 g, 3.34 mmol, 2.2 eq) as starting materials to afford the title compound **28** (280 mg, 0.74 mmol, 45 % yield). **$^1$H NMR** (300 MHz, $CDCl_3$) δ 3.18 (t, *J* = 7.1 Hz, 2H), 2.48 – 2.28 (m, 2H), 2.03 (t, *J* = 1.6 Hz, 3H), 1.82 (p, *J* = 7.1 Hz, 2H), 1.60 – 1.47 (m, 2H), 1.43 – 1.17 (m, 20H), 0.77 (s, 2H). **$^{13}$C NMR** (75 MHz, $CDCl_3$) δ 110.2, 105.3, 33.7, 30.7, 29.8, 29.8, 29.7, 29.6, 29.5, 28.7, 27.5, 26.1, 11.7, 8.3, 7.3.

**15-(2-methylcycloprop-1-en-1-yl)pentadecanenitrile (31)**

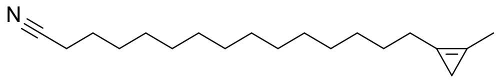

Classified as Internal | Intern

General procedure E was used with compound **28** (280 mg, 0.74 mmol, 1.0 eq) as starting material to afford the title compound **31** (140 mg, 0.51 mmol, 68 % yield). **$^1$H NMR** (300 MHz, $CDCl_3$) δ 2.49 – 2.28 (m, 4H), 2.03 (t, *J* = 1.6 Hz 3H), 1.75 – 1.40 (m, 6H), 1.39 – 1.18 (m, 18H), 0.79 (s, 2H). **$^{13}$C NMR** (75 MHz, $CDCl_3$) δ 120.0, 110.2, 105.3, 29.8, 29.7, 29.6, 29.6, 29.5, 29.4, 28.9, 28.8, 27.5, 26.1, 25.5, 17.3, 11.7, 8.2.

**15-(2-methylcycloprop-1-en-1-yl)pentadecanoic acid (2)**

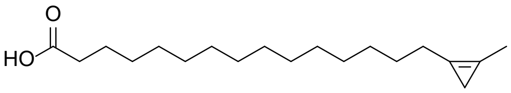

General procedure F was used with compound **31** (140 mg, 0.51 mmol, 1.0 eq) as starting material to afford the title compound **2** (89 mg, 0.30 mmol, 60% yield). **$^1$H NMR** (300 MHz, $CDCl_3$) δ 10.38 (br s, 1H), 2.43 – 2.29 (m, 4H), 2.03 (t, *J* = 1.6 Hz, 3H), 1.63 (p, *J* = 7.4 Hz, 2H), 1.59 – 1.46 (m, 2H), 1.38 – 1.17 (m, 20H), 0.77 (s, 2H). **$^{13}$C NMR** (75 MHz, $CDCl_3$) δ 180.6, 110.2, 105.3, 34.3, 29.8, 29.8, 29.8, 29.8, 29.7, 29.6, 29.6, 29.6, 29.4, 29.2, 27.5, 26.1, 24.8, 11.7, 8.3.

**2-bromodec-1-ene (24)**

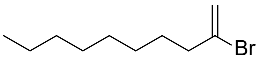

General procedure B was used with dec-1-yne (0.90 mL, 5.0 mmol, 1.0 eq) as starting material to afford the title compound **24** (854 mg, 3.90 mmol, 78% yield) as a colorless liquid. **$^1$H NMR** (400 MHz, $CDCl_3$): δ 5.55 (dt, *J* = 1.6 Hz, 1.2 Hz, 1H), 5.38 (d, *J* = 1.6 Hz, 1H), 2.49 – 2.34 (dt, *J* = 2.8, 1.2, 3H), 1.65 – 1.49 (m, 3H), 0.95 – 0.82 (t, *J* = 7.0 Hz, 3H). **$^{13}$C NMR** (101 MHz, $CDCl_3$): δ 135.1, 116.3, 41.6, 32.0, 29.4, 29.4, 28.6, 28.1, 22.8, 14.3. Spectral data was in accordance with the literature.[8]

**1,1,2-tribromo-2-octylcyclopropane (26)**

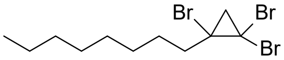

General procedure C was used with compound **24** (3.22 g, 14.7 mmol, 1.0 eq) as starting material to afford the title compound **26** (2.69 g, 6.88 mmol, 47% yield) as a colorless liquid. **$^1$H NMR** (400 MHz, $CDCl_3$): δ 2.11-1.93 (m, 2H) 1.97 – 1.93 (d, *J* = 9.2 Hz 1H), 1.82 (d, *J* = 9.2 Hz, 1H), 1.79 – 1.53 (m, 1H), 1.43 – 1.16 (m, 14H), 0.95 – 0.85 (t, *J* = 7.0 Hz , 3H). **$^{13}$C NMR** (101 MHz, $CDCl_3$): δ 46.0, 41.9, 38.2, 33.3, 32.0, 29.6, 29.4, 29.1, 27.8, 22.8, 14.3. Spectral data was in accordance with the literature.[3]

**1-(7-iodoheptyl)-2-octylcycloprop-1-ene (29)**

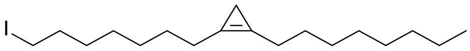

General procedure D was used with compound **26** (3.91 g, 10.0 mmol, 1.0 eq) and 1,7-diiodoheptane (8.10 g, 23.0 mmol, 2.3 eq) as starting materials to afford the title compound **29** (2.10 g, 5.59 mmol, 56 % yield). **$^1$H NMR** (400 MHz, $CDCl_3$): δ 3.18 (t, *J* = 7.0 Hz, 2H), 2.41 – 2.33 (m, 4H), 1.82 (p, *J* = 7.1 Hz, 2H), 0.92-0.85 (m, 3H), 0.77 (s, 2H). ). **$^{13}$C NMR** (101 MHz, $CDCl_3$): δ 109.6, 109.3, 33.7, 32.0, 30.6, 29.6, 29.4, 29.3, 28.5, 27.5, 27.4, 26.2, 26.1, 22.8, 14.3, 7.5, 7.3.

Classified as Internal | Intern

**8-(2-octylcycloprop-1-en-1-yl)octanenitrile (32)**

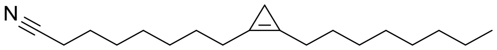

General procedure E was used with compound **29** (2.10 g, 5.59 mmol, 1.0 eq) as starting material to afford the title compound **32** (1.39 g, 5.00 mmol, 89 % yield) as a clear oil. **$^1$H NMR** (300 MHz, $CDCl_3$): δ 0.92 – 0.85 (m, 3H), 0.76 (s, 2H). **$^{13}$C NMR** (75 MHz, $CDCl_3$): δ 119.9, 109.7, 109.2, 32.0, 29.5, 29.4, 29.2, 28.8, 28.7, 27.5, 27.4, 26.2, 26.0, 25.5, 22.8, 17.3, 14.3, 7.5.

**8-(2-octylcycloprop-1-en-1-yl)octanoic acid (4)**

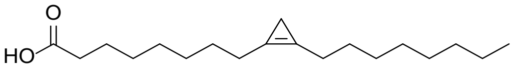

General procedure F was used with compound **32** (1.35 g, 4.92 mmol, 1.0 eq) as starting material to afford the title compound **4** (909 mg, 3.09 mmol, 63% yield) as a light yellow oil. **$^1$H NMR** (300 MHz, $CDCl_3$): δ 2.24-2.28 (m, 6H) 0.93 – 0.83 (m, 3H), 0.77 (s, 2H). **$^{13}$C NMR** (75 MHz, $CDCl_3$): δ 180.3, 109.6, 109.4, 34.2, 32.0, 29.6, 29.4, 29.3, 29.2, 27.5, 27.5, 26.2, 26.1, 24.8, 22.8, 14.3, 7.5.

**9-bromononanoic acid (s4)**

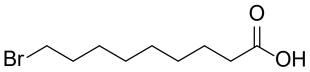

General procedure A was used with compound 9-bromononan-1-ol (3.05 g, 13.67 mmol, 1.0 eq) as starting material, the crude product was purified by silica gel column chromatography ($Et_2O$ / pentane 1:19, 1% AcOH) to afford compound **s4** (3.09 g, 13.0 mmol, 95% yield) as a yellow oil. **$^1$H NMR** (500 MHz, $CDCl_3$) δ 11.64 (s, 1H), 3.39 (t, *J* = 6.8 Hz, 2H), 2.34 (t, *J* = 7.5 Hz, 2H), 1.89 – 1.79 (m, 2H), 1.67 – 1.58 (m, 2H), 1.46 – 1.38 (m, 2H), 1.37 – 1.27 (m, 6H).**$^{13}$C NMR** (126 MHz, $CDCl_3$) δ 180.6, 34.2, 34.1, 32.9, 29.1, 29.0, 28.7, 28.2, 24.7**.** Spectral data was in accordance with the literature.[19]

**(8-carboxyoctyl)triphenylphosphonium bromide (s5)**

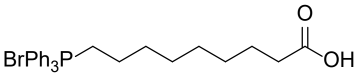

General procedure G was used with compound **s4** (3.09 g, 13.0 mmol, 1.0 eq) as starting material, the crude product was purified by silica gel column chromatography (DCM / MeOH 99:1 to 1:19, 1% AcOH) to afford compound **s5** (6.45 g, 12.9 mmol, quant. yield) as a yellow oil. **$^1$H NMR** (500 MHz, $CDCl_3$) δ 7.80 – 7.72 (m, 9H), 7.70 – 7.65 (m, 6H), 3.62 – 3.50 (m, 2H), 2.28 (t, *J* = 7.4 Hz, 2H), 1.56 (dd, *J* = 10.4, 6.2 Hz, 4H), 1.48 (q, *J* = 7.3 Hz, 2H), 1.26 – 1.11 (m, 6H). **$^{13}$C NMR** (126 MHz, $CDCl_3$) δ 177.2, 135.2, 135.2, 133.7, 133.6, 130.7, 130.6, 118.6, 117. 9, 34.5, 30.2, 30.0, 28.5, 28.5, 24.6, 22.9, 22.5, 22.5. Spectral data was in accordance with the literature.[20]

**(2-(1,3-dioxan-2-yl)ethyl)triphenylphosphonium bromide (s6)**

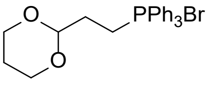

Classified as Internal | Intern

General procedure A was used with 2-(2-bromoethyl)-1,3-dioxane (1.0 mL, 7.34 mmol, 1.0 eq) as starting material to afford compound **s6** (500 mg, 1.09 mmol, 15% yield) as a yellow oil. $^{1}$**H NMR** (300 MHz, $CDCl_3$) δ 7.74 – 7.39 (m, 15H), 4.79 (t, *J* = 4.7 Hz, 1H), 3.83 – 3.73 (m, 2H), 3.60 (td, *J* = 12.2, 2.4 Hz, 2H), 3.54 – 3.38 (m, 2H), 1.86 – 1.74 (m, 1H), 1.73 – 1.59 (m, 2H), 1.16 – 1.05 (m, 1H). $^{13}$**C NMR** (75 MHz, $CDCl_3$) δ 134.8, 134.8, 133.1, 133.0, 130.3, 130.1, 118.0, 116.8, 99.0, 98.8, 66.3, 27.5, 27.5, 25.1, 17.6, 16.9.

**methyl 5-oxopentanoate (s7)**

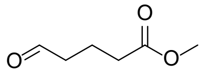

Triethylamine (22.6 mL, 122 mmol, 5 eq) was added too a solution of δ-valerolactone (3.0 mL, 32.4 mmol, 1 eq) in MeOH (0.5 M) at rt and the solution was stirred for 1 h. The solvents were removed and coevaporated with toluene to afford the crude alcohol. Dess-Martin periodinane (16.3 g, 38.4 mmol, 1.2 eq) was added to a solution of the crude alcohol (5.13 g, 38.9 mmol, 1 eq) in anh. DCM at 0 °C under $N_2$ atmosphere and stirred at rt for 2 h. The reaction mixture was quenched with sat. aq. $NaHCO_3$ and extracted with $Et_2O$ (3x). The combined organic layers were washed with brine, dried over $MgSO_4$ and solvents removed under reduced pressure. The crude product was purified by silica gel column chromatography ($Et_2O$ / pentane 1:4) to afford compound **s7** (1.96 g, 15.0 mmol, 47% yield) as a light yellow oil. $^{1}$**H NMR** (400 MHz, $CDCl_3$) δ 9.75 (t, *J* = 1.2 Hz, 1H), 3.65 (d, *J* = 0.9 Hz, 3H), 2.51 (tt, *J* = 7.2, 1.1 Hz, 2H), 2.36 (t, *J* = 7.3 Hz, 2H), 2.06 – 1.84 (m, 2H). $^{13}$**C NMR** (101 MHz, $CDCl_3$) δ 201.6, 173.5, 51.7, 43.0, 33.0, 17.4. Spectral data was in accordance with the literature.[21]

**methyl (Z)-7-(1,3-dioxan-2-yl)hept-5-enoate (s8)**

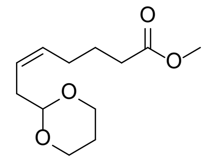

A solution of NaHMDS (1 M, 11.5 mL, 11.5 mmol, 1.3 eq) was added dropwise to a solution of compound **s6** (5.25 g, 11.5 mmol, 1.3 eq) in anh. THF (0.25 M) at -78 °C under $N_2$ atmopshere, slowly warmed to rt and stirred for 1 h. The reaction was cooled back to -78 °C and a solution of compound **s7** (1.15 g, 8.83 mmol, 1 eq) in anh. THF (0.5 M) was added dropwise to the orange suspension and stirred for 1.5 h. The reaction mixture was quenched with sat. aq. $NH_4Cl$ and extracted with $Et_2O$ (3x). The combined organic layers were washed with brine, dried over $MgSO_4$ and solvents removed under reduced pressure. The crude product was purified by silica gel column chromatography (acetone / pentane 1:49 to 1:19) to afford compound **s8** (1.31 g, 5.75 mmol, 65% yield) as a clear colorless oil. $^{1}$**H NMR** (300 MHz, $CDCl_3$) δ 5.58 – 5.33 (m, 2H), 4.51 (t, *J* = 5.2 Hz, 1H), 4.18 – 3.99 (m, 2H), 3.80 – 3.69 (m, 2H), 3.64 (s, 3H), 2.39 – 2.21 (m, 4H), 2.16 – 1.94 (m, 3H), 1.69 (dt, *J* = 8.3, 7.1 Hz, 2H), 1.32 (ddt, *J* = 13.5, 2.6, 1.3 Hz, 1H). $^{13}$**C NMR** (75 MHz, $CDCl_3$) δ 174.1, 131.3, 124.3, 101.9, 67.0, 51.5, 33.5, 26.8, 25.8, 24.8. Spectral data was in accordance with the literature.[22]

**hexadec-9-yn-1-ol (s10)**

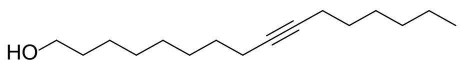

In a slightly modified procedure from the literature[5], *n*-BuLi (2.5 M in hexanes, 8.80 mL, 22 mmol, 2.2 eq) was added dropwise to a solution of dec-9-yn-1-ol (1.54 g, 10.0 mmol, 1 eq) in 4:1 THF / HMPA

Classified as Internal | Intern

(0.1 M) at -78 °C under $N_2$ atmosphere and the reaction mixture was stirred for 1 h gradually warming to rt. The formed suspension was cooled back to -78 °C, 1-bromohexane (1.35 ml, 9.6 mmol, 0.96 eq) and NaI (75 mg, 0.50 mmol, 0.05 eq) were added dropwise and the reaction was stirred for 18 h slowly warming to rt. The solution was quenched with water and extracted with 1:1 diethyl ether / pentane (3x). The combined organic layers were washed with brine, dried over $MgSO_4$ and concentrated under reduced pressure. The crude product was purified by silica gel column chromatography ($Et_2O$ / pentane 1:19 to 1:9) to afford compound **s10** (1.04 g, 4.38 mmol, 43.8 % yield) as a colorless oil. **$^1$H NMR** (400 MHz, $CDCl_3$) δ 3.47 (t, *J* = 6.8 Hz, 2H), 3.22 (s, 1H), 2.02 (t, *J* = 7.0 Hz, 4H), 1.56 – 1.09 (m, 20H), 0.78 (t, *J* = 7.0 Hz, 3H). $^{13}$C NMR (101 MHz, $CDCl_3$) δ 80.1, 80.0, 62.4, 32.6, 31.3, 29.4, 29.1, 29.1, 29.1, 28.8, 28.5, 25.7, 22.5, 18.7, 18.7, 14.0.

**hexadec-9-ynoic acid (s11)**

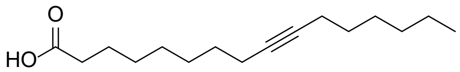

General procedure A was used with compound **s10** (402 mg, 1.69 mmol, 1.0 eq) as starting material, the crude product was purified by silica gel column chromatography ($Et_2O$ / pentane 1:9) to afford compound **s11** (357 mg, 1.41 mmol, 84% yield) as a light yellow crystalline solid. **$^1$H NMR** (400 MHz, $CDCl_3$) δ 2.33 (t, *J* = 7.5 Hz, 2H), 2.12 (t, *J* = 6.6 Hz, 4H), 1.69 – 1.57 (m, 2H), 1.51 – 1.17 (m, 16H), 0.87 (t, *J* = 6.9 Hz, 3H). **$^{13}$C NMR** (101 MHz, $CDCl_3$) δ 180.6, 80.4, 80.1, 34.2, 31.5, 29.2, 29.2, 29.1, 28.9, 28.7, 28.7, 24.7, 22.7, 18.9, 18.8, 14.2. Spectral data was in accordance with the literature.[6]

**methyl hexadec-9-ynoate (s12)**

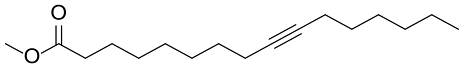

Thionylchloride (0.41 mL, 5.65 mmol, 4.0 eq) was added to a solution of compound **s11** (357 mg, 1.41 mmol, 1.0 eq) in methanol (0.2 M) at 0 °C under nitrogen atmosphere and stirred for 2.5 h at rt. The reaction mixture was concentrated under reduced pressure and the crude product was purified by silica gel column chromatography ($Et_2O$ / pentane 1:49) to afford compound **s12** (333 mg, 1.25 mmol, 88% yield) as a clear oil. **$^1$H NMR** (300 MHz, $CDCl_3$) δ 3.62 (s, 3H), 2.26 (t, *J* = 7.5 Hz, 2H), 2.13 – 2.04 (m, 4H), 1.65 – 1.51 (m, 2H), 1.49 – 1.15 (m, 16H), 0.85 (t, *J* = 6.7 Hz, 3H). **$^{13}$C NMR** (75 MHz, $CDCl_3$) δ 174.3, 80.4, 80.1, 51.4, 34.1, 31.5, 29.2, 29.1, 29.1, 28.9, 28.7, 28.6, 25.0, 22.7, 18.8, 18.8, 14.1

**ethyl 2-hexyl-3-(8-methoxy-8-oxooctyl)cycloprop-2-ene-1-carboxylate (s13)**

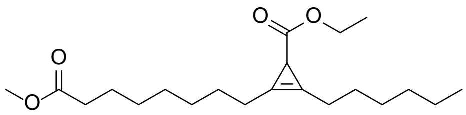

A solution of ethyl diazoacetate (15 wt. % in DCM, 0.9 mL, 1.29 mmol, 1.0 eq) was added dropwise over 3 hours to a suspension of compound **s12** (333 mg, 1.25 mmol, 1.0 eq) and rhodium(II) acetate

Classified as Internal | Intern

dimer (28 mg, 0.063 mmol, 0.05 eq) in DCM (0.2 M) at 0 °C under $N_2$ and stirred for another 30 minutes. The reaction mixture was concentrated under reduced pressure and the crude product was purified by silica gel column chromatography ($Et_2O$ / pentane 1:49 to 1:9) to afford the compound **s13** (327 mg, 0.928 mmol, 76% yield) as a light yellow oil. $^1$**H NMR** (400 MHz, $CDCl_3$) δ 4.06 (q, *J* = 7.1 Hz, 2H), 3.62 (s, 3H), 2.35 (t, *J* = 7.3 Hz, 4H), 2.26 (t, *J* = 7.5 Hz, 2H), 1.98 (s, 1H), 1.62 – 1.55 (m, 2H), 1.55 – 1.45 (s, 4 Hz), 1.35 – 1.16 (m, 12H), 1.20 (t, *J* = 7.2 Hz, 3H), 0.85 (t, *J* = 6.8 Hz, 3H). $^{13}$**C NMR** (101 MHz, $CDCl_3$) δ 177.1, 174.3, 105.8, 105.6, 59.8, 51.5, 34.1, 31.6 29.1, 29.1, 29.0, 29.0, 27.0, 27.0, 25.0, 24.6, 24.5, 22.6, 22.3, 14.5, 14.1.

**2-(7-carboxyheptyl)-3-hexylcycloprop-2-ene-1-carboxylic acid (s14)**

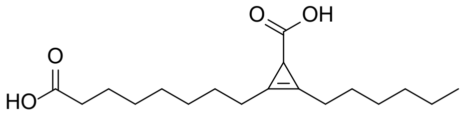

An aqueous solution of KOH (2.0 M, 4.2 mmol, 2.1 mL, 5.0 eq) was added to a solution of compound **s13** (296 mg, 0.840 mmol, 1.0 eq) in ethanol (0.2 M) and refluxed for 2.5 h. The reaction mixture was acidified with aq. 1 M HCl and extracted with $Et_2O$ (5x). The combined organic layers were washed with brine, dried over $MgSO_4$ and concentrated under reduced pressure. The crude product was purified by silica gel column chromatography ($Et_2O$ / pentane 1:3 + 1% AcOH) to afford the compound **s14** (254 mg, 0.819 mmol, 97% yield) as a clear oil. $^1$**H NMR** (400 MHz, $CDCl_3$) δ 11.07 (s, 2H), 2.38 (t, *J* = 7.2 Hz, 4H), 2.30 (t, *J* = 7.5 Hz, 2H), 1.99 (s, 1H), 1.64 – 1.46 (m, 6H), 1.38 – 1.18 (m, 12H), 0.85 (t, *J* = 6.7 Hz, 3H).

Classified as Internal | Intern

**2-bromohept-1-ene (11)**

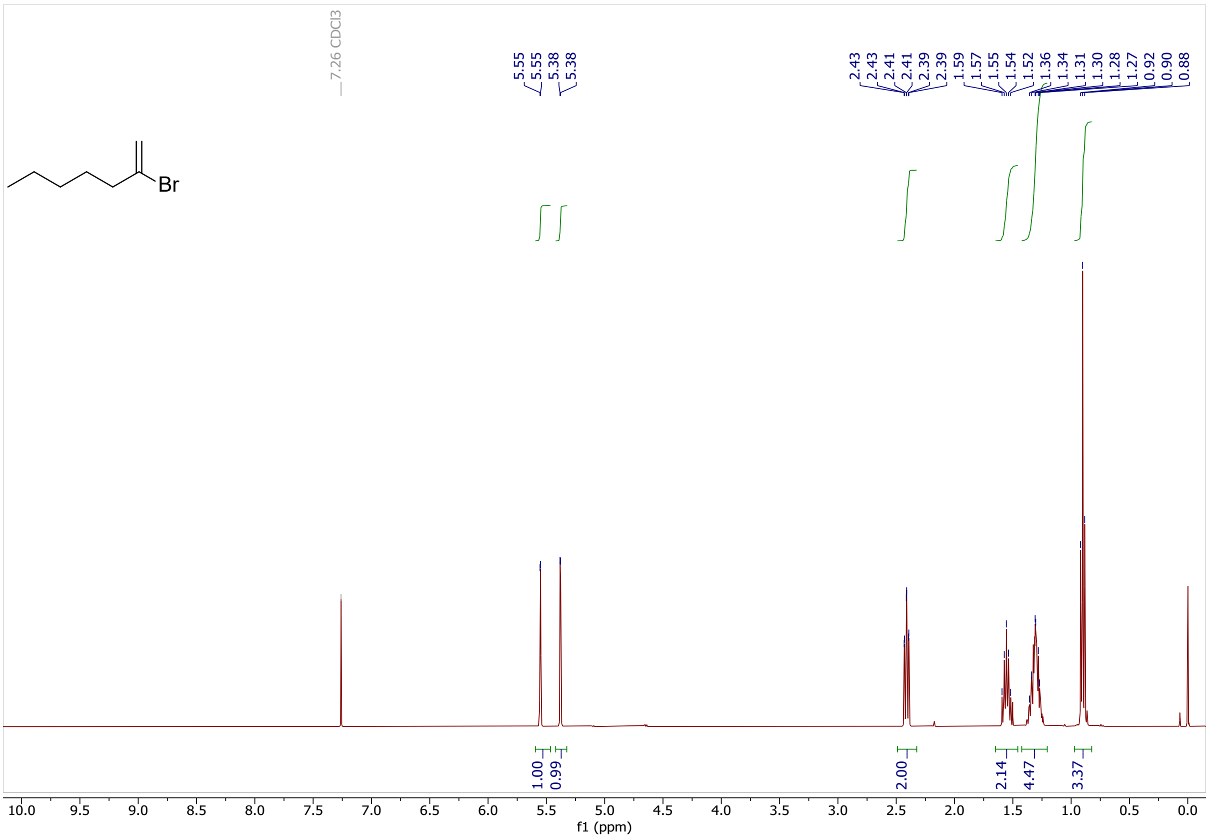

Classified as Internal | Intern

### 1,1,2-tribromo-2-pentylcyclopropane (12)

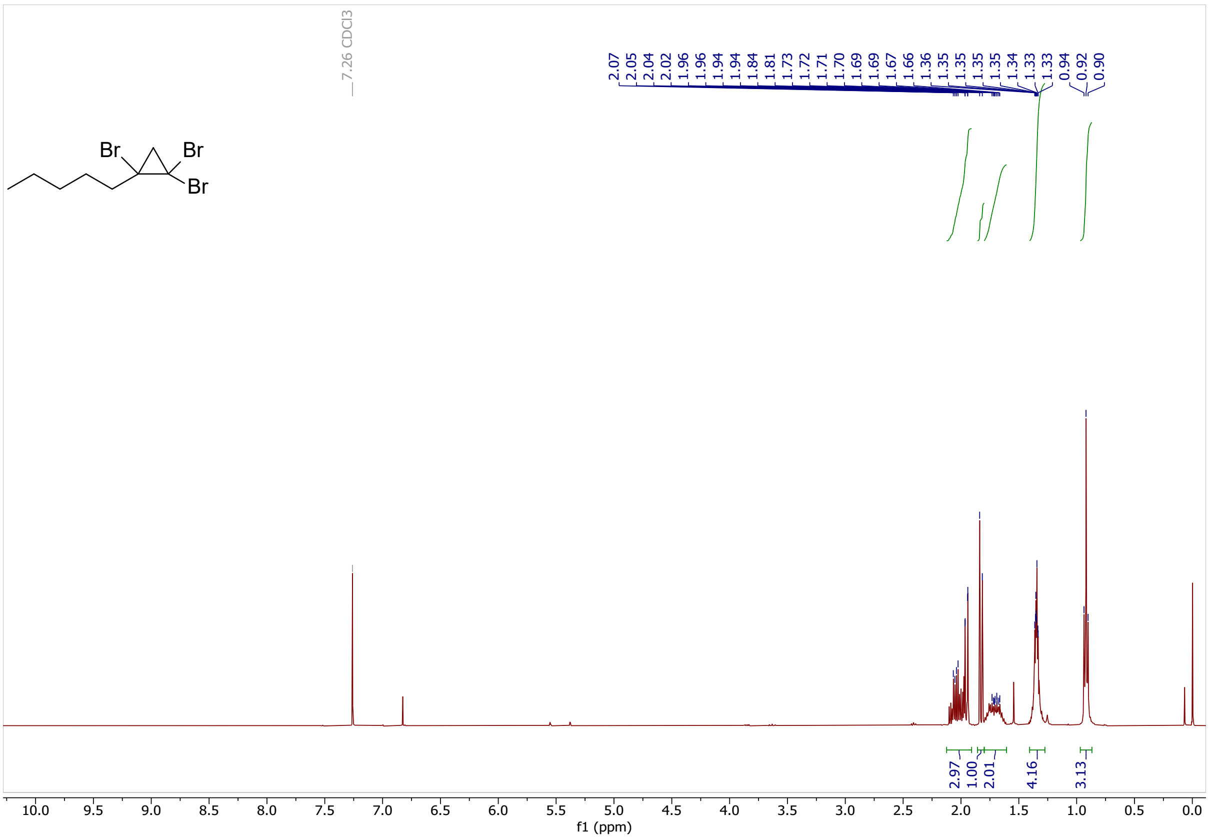

Classified as Internal | Intern

**2-(2-pentylcycloprop-1-en-1-yl)ethan-1-ol (13)**

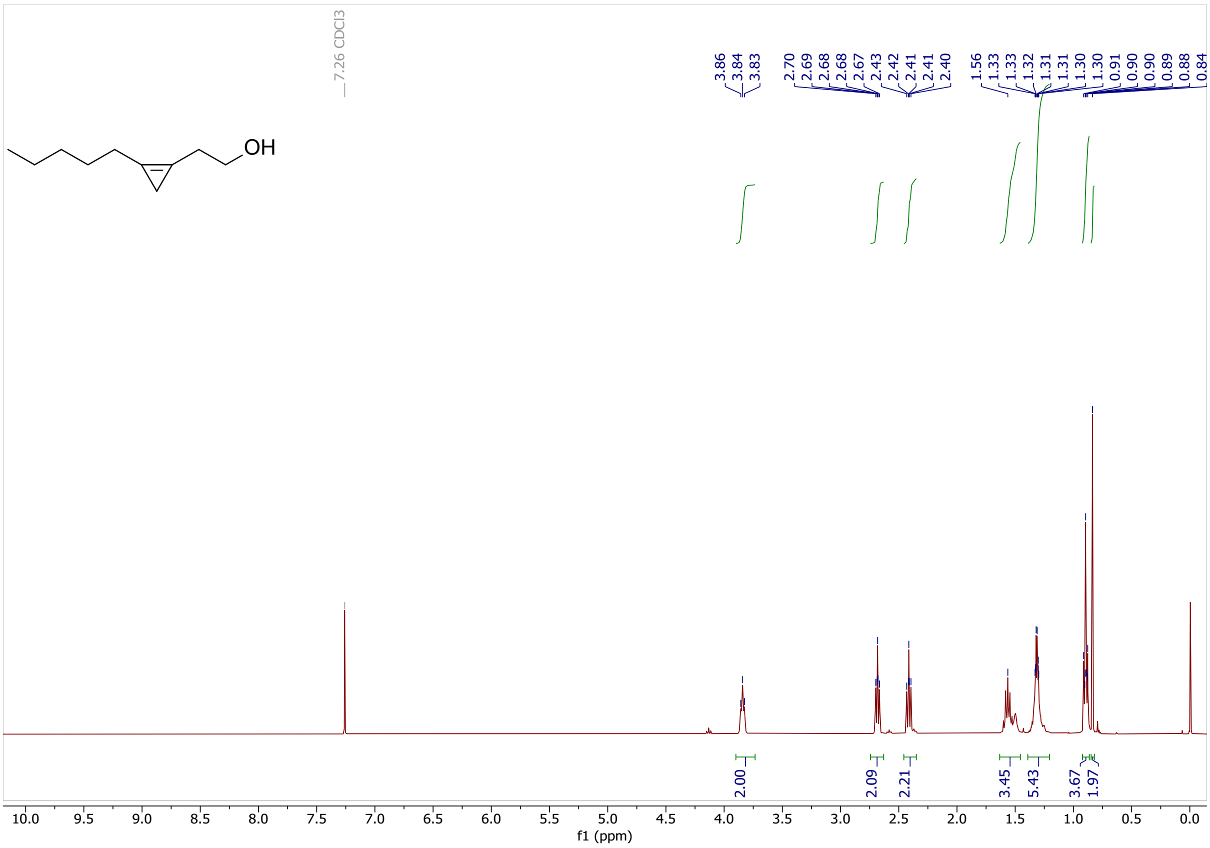

Classified as Internal | Intern

### 1-(2-bromoethyl)-2-pentylcycloprop-1-ene (14)

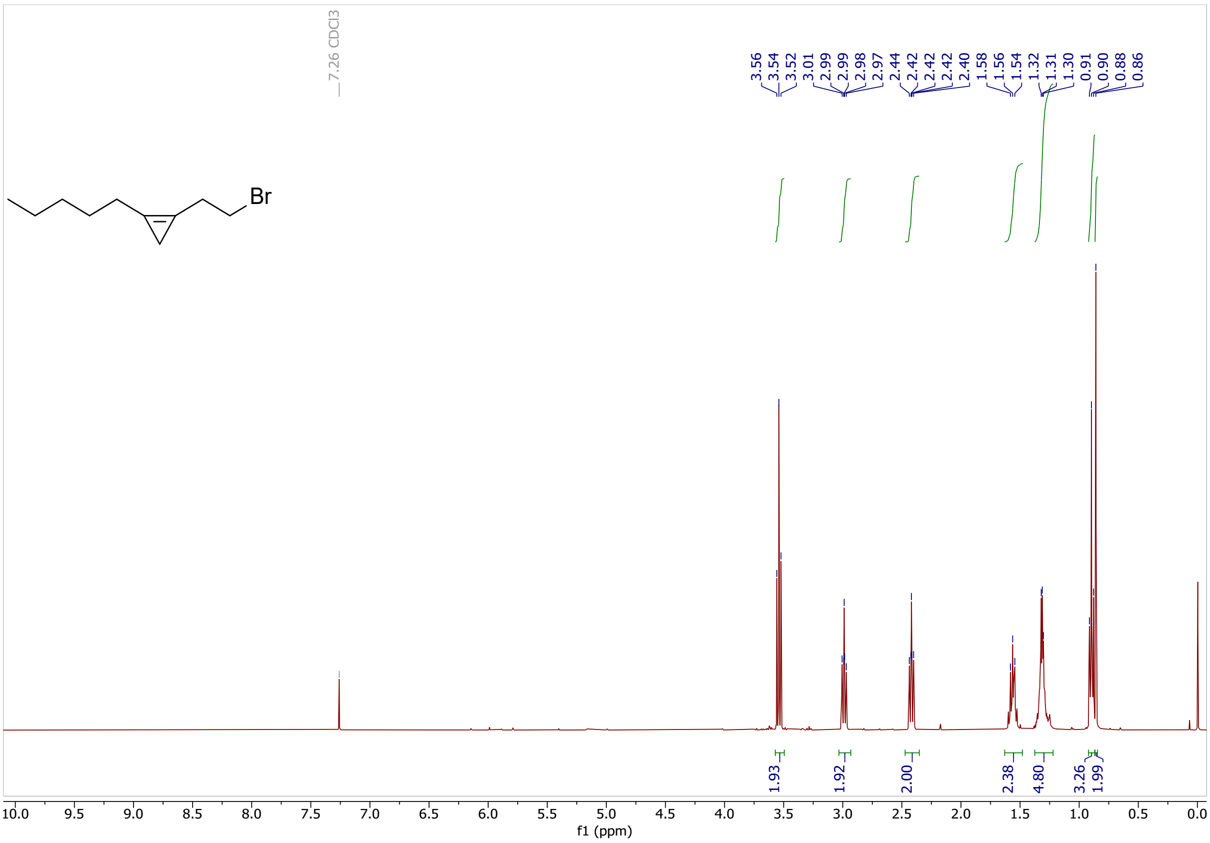

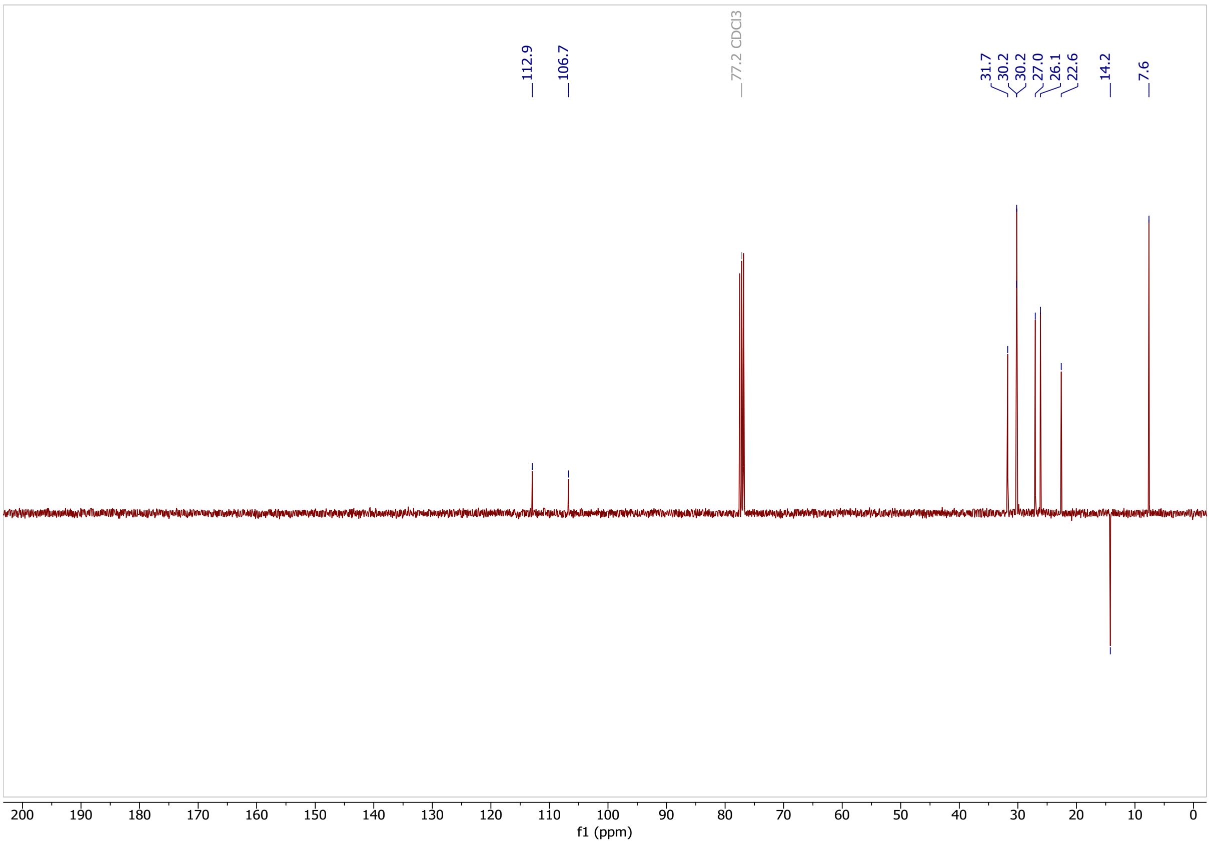

Classified as Internal | Intern

**(2-(2-pentylcycloprop-1-en-1-yl)ethyl)triphenylphosphonium bromide (7)**

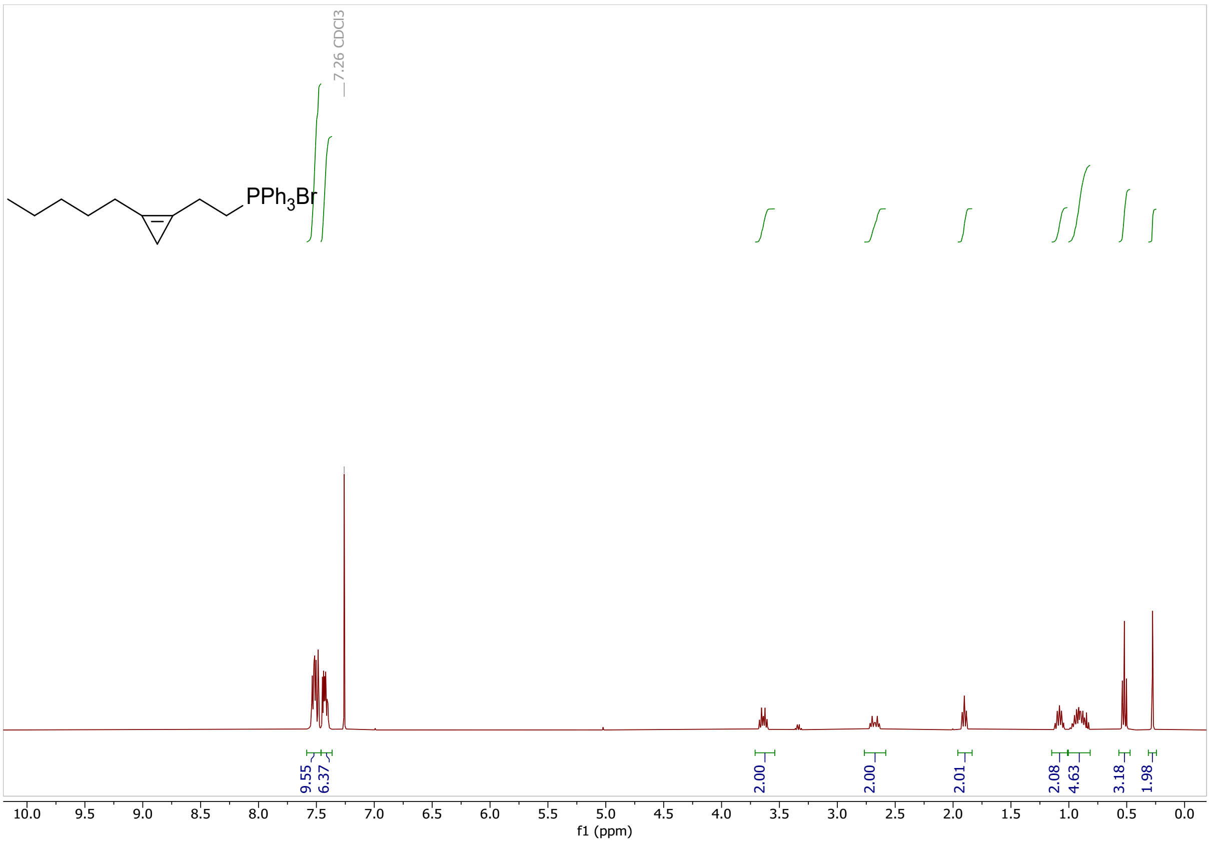

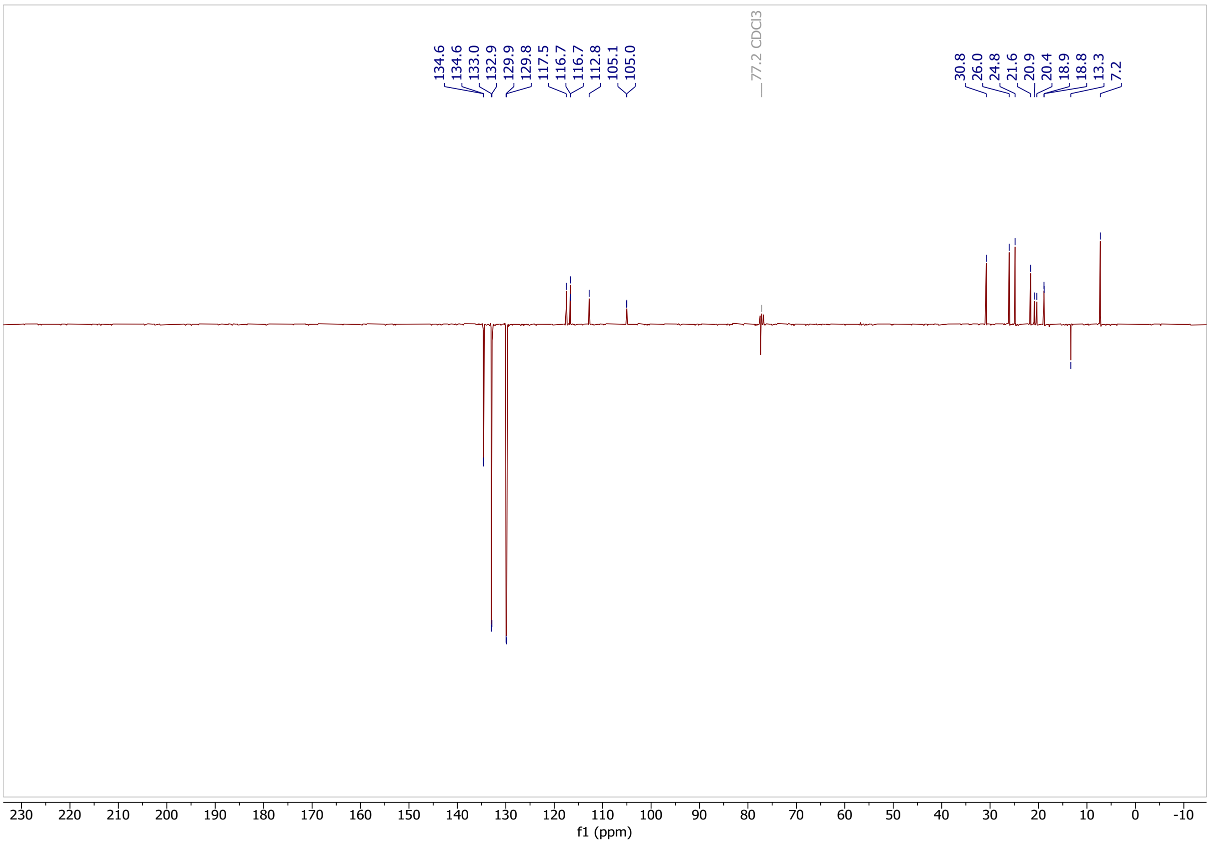

Classified as Internal | Intern

### 9-((tert-butyldimethylsilyl)oxy)nonan-1-ol (14)

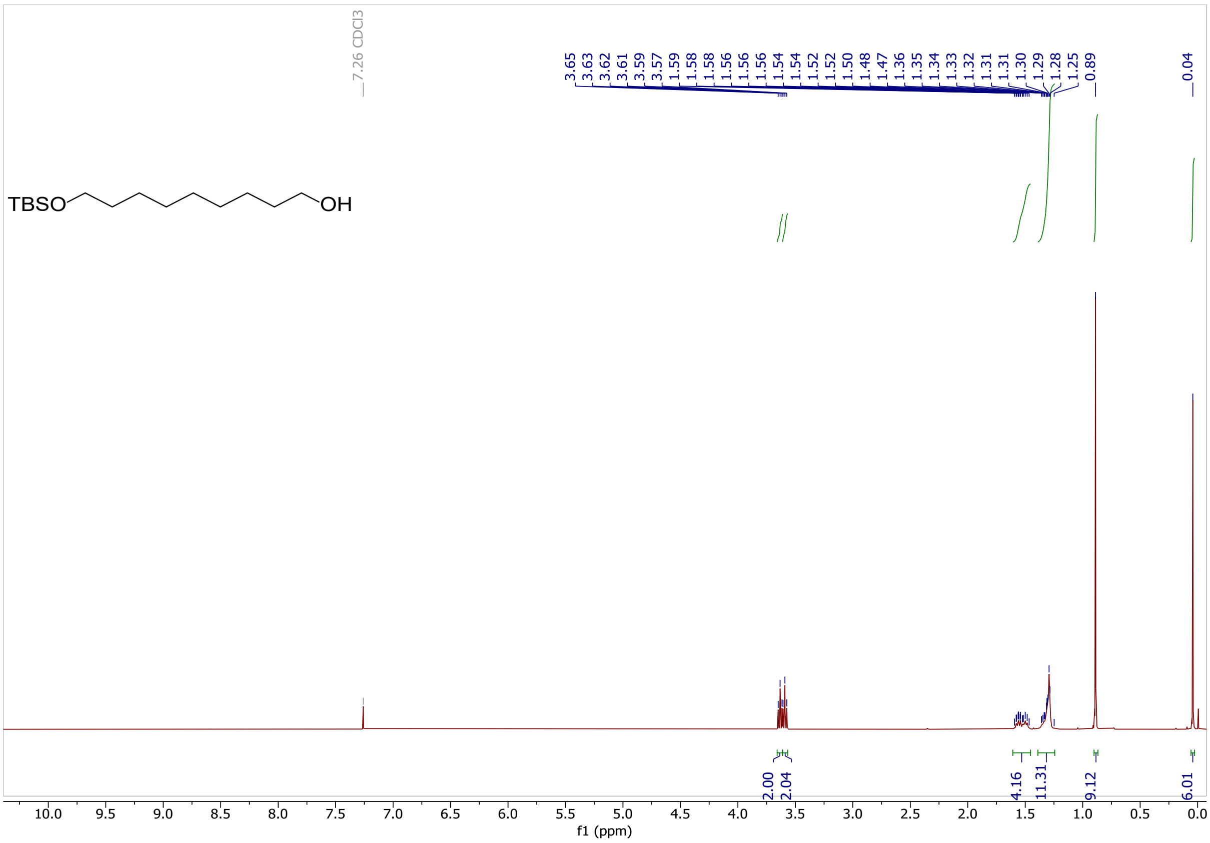

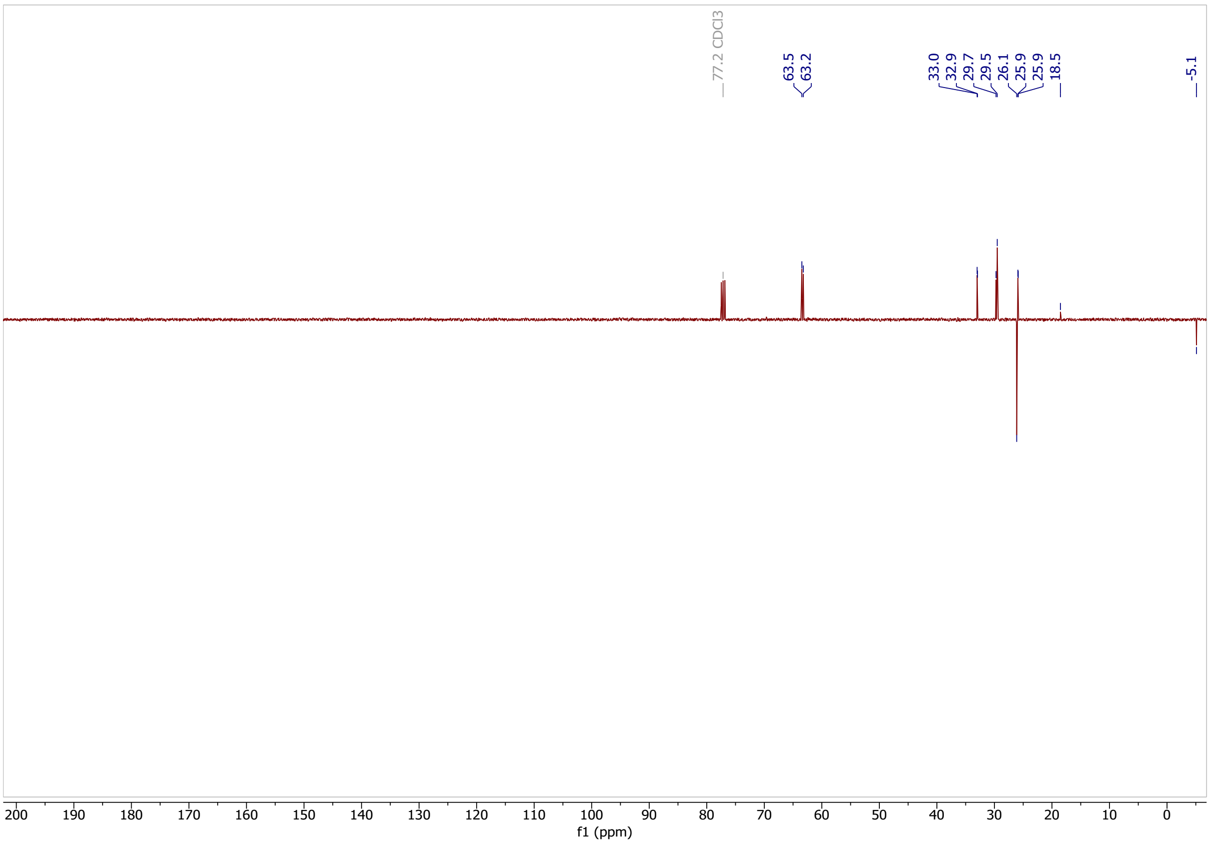

Classified as Internal | Intern

**9-((tert-butyldimethylsilyl)oxy)nonanoic acid (15)**

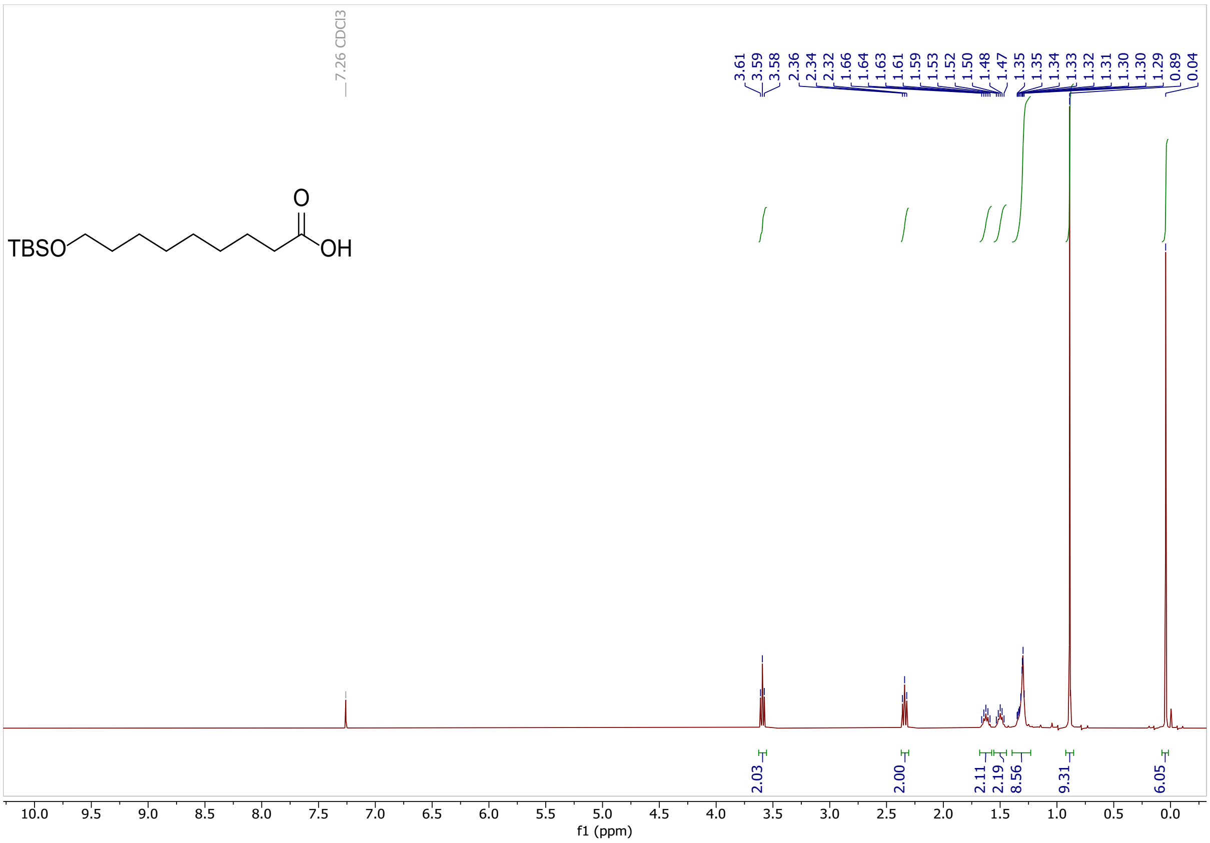

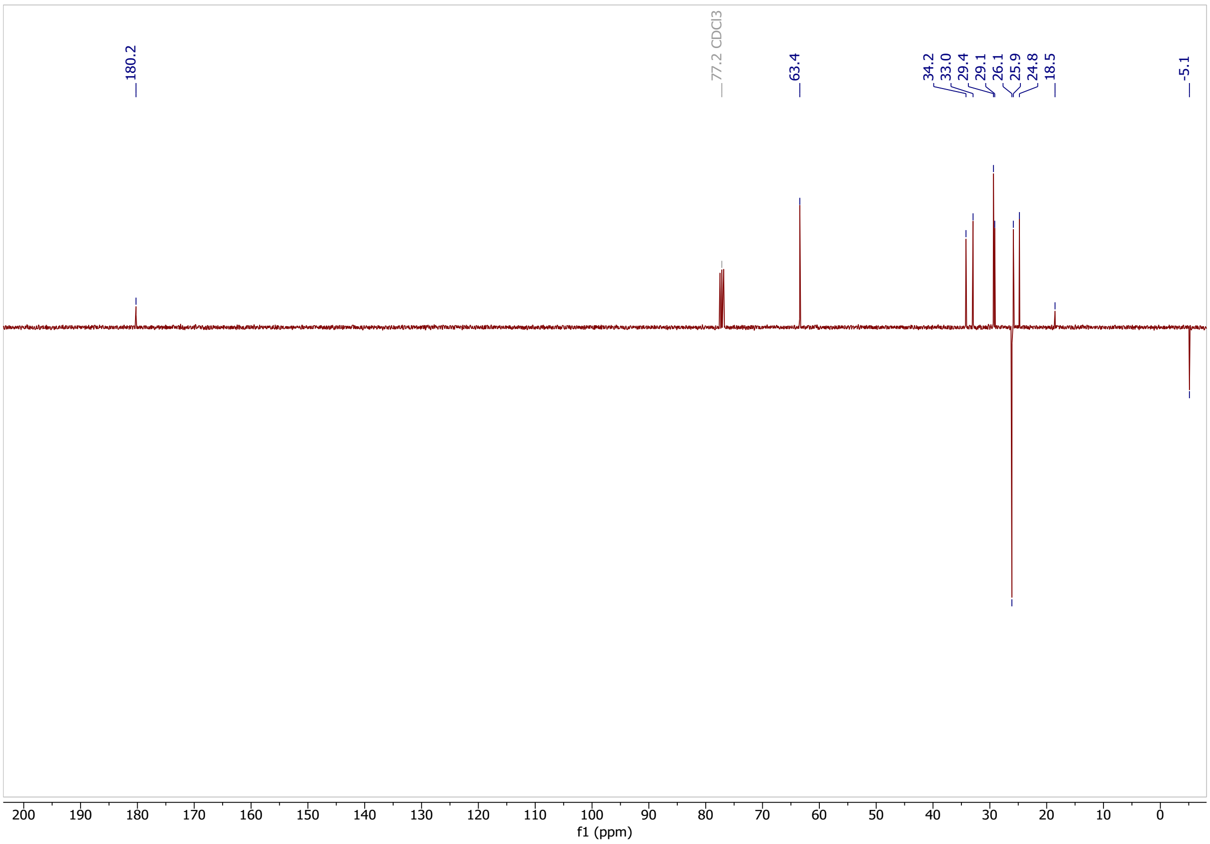

Classified as Internal | Intern

**methyl 9-((tert-butyldimethylsilyl)oxy)nonanoate (16)**

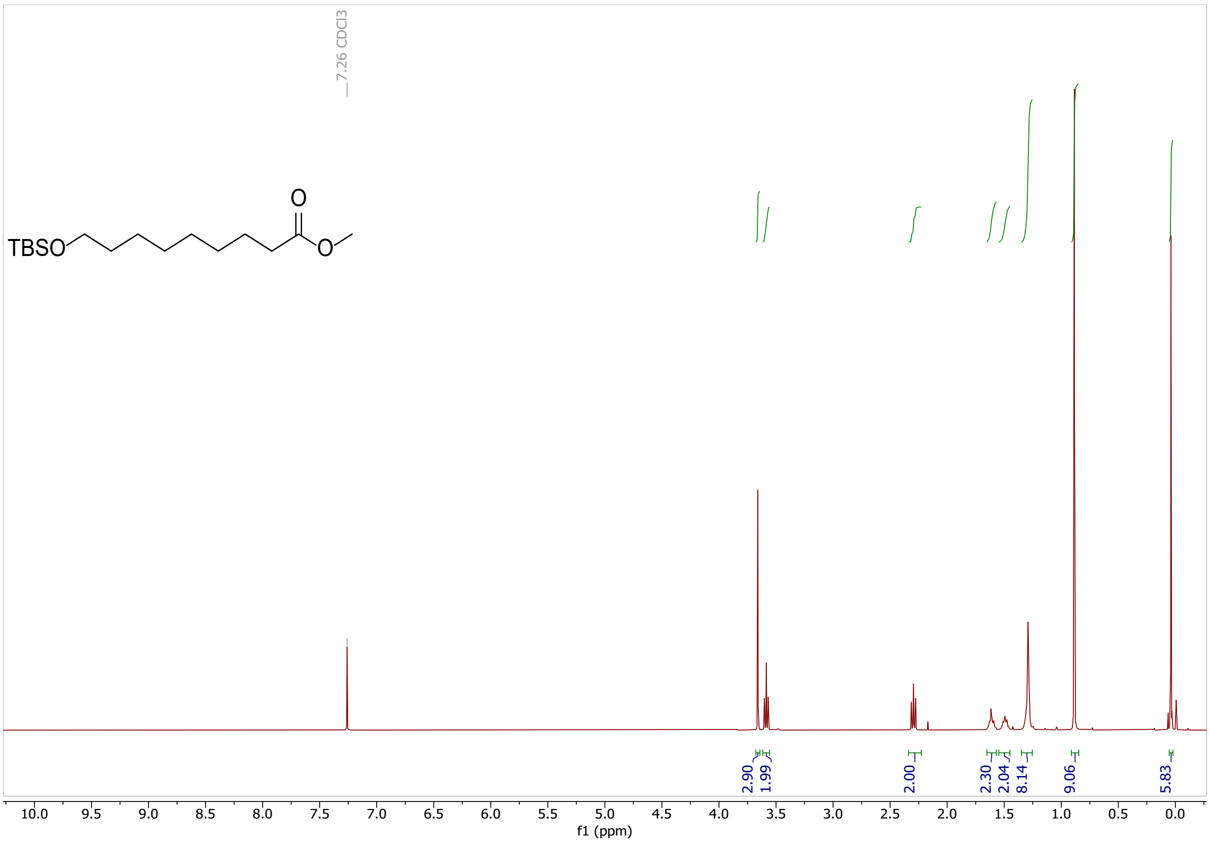

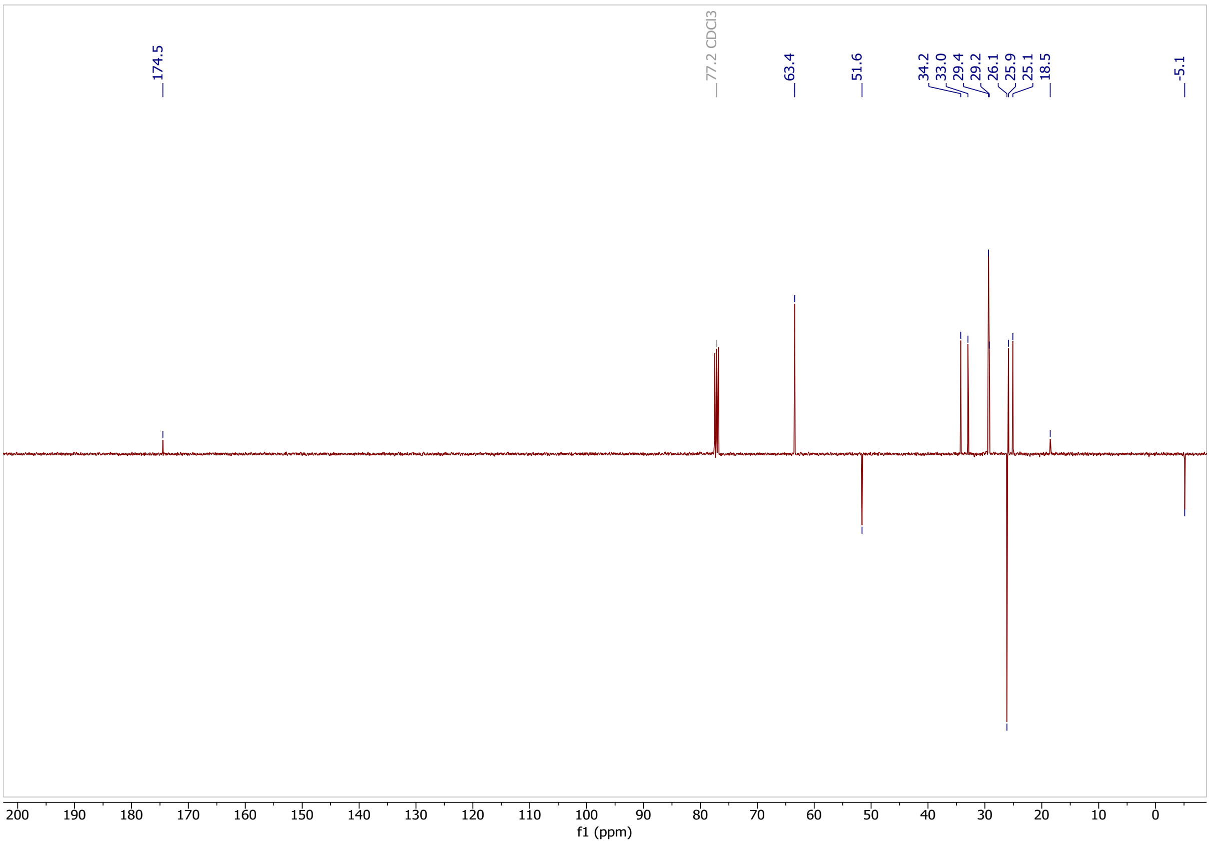

Classified as Internal | Intern

**methyl 9-hydroxynonanoate (17)**

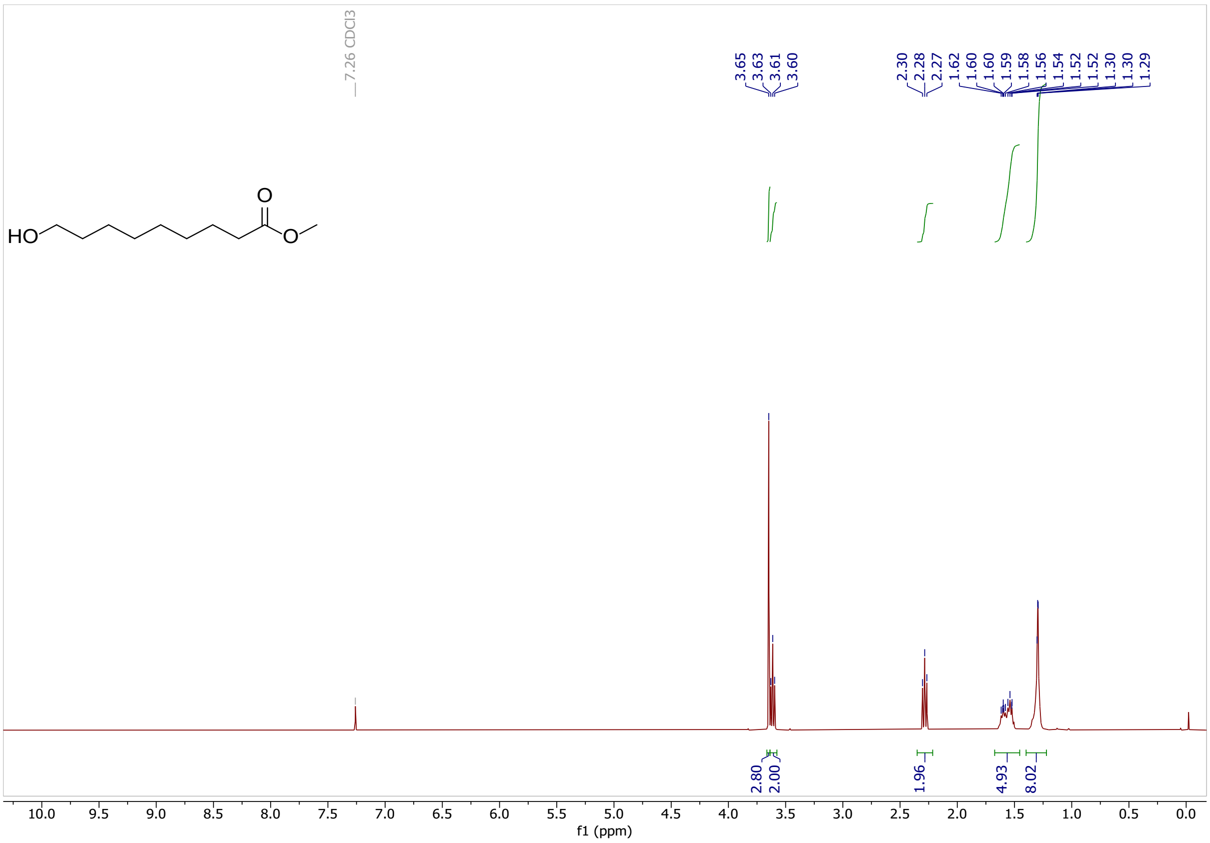

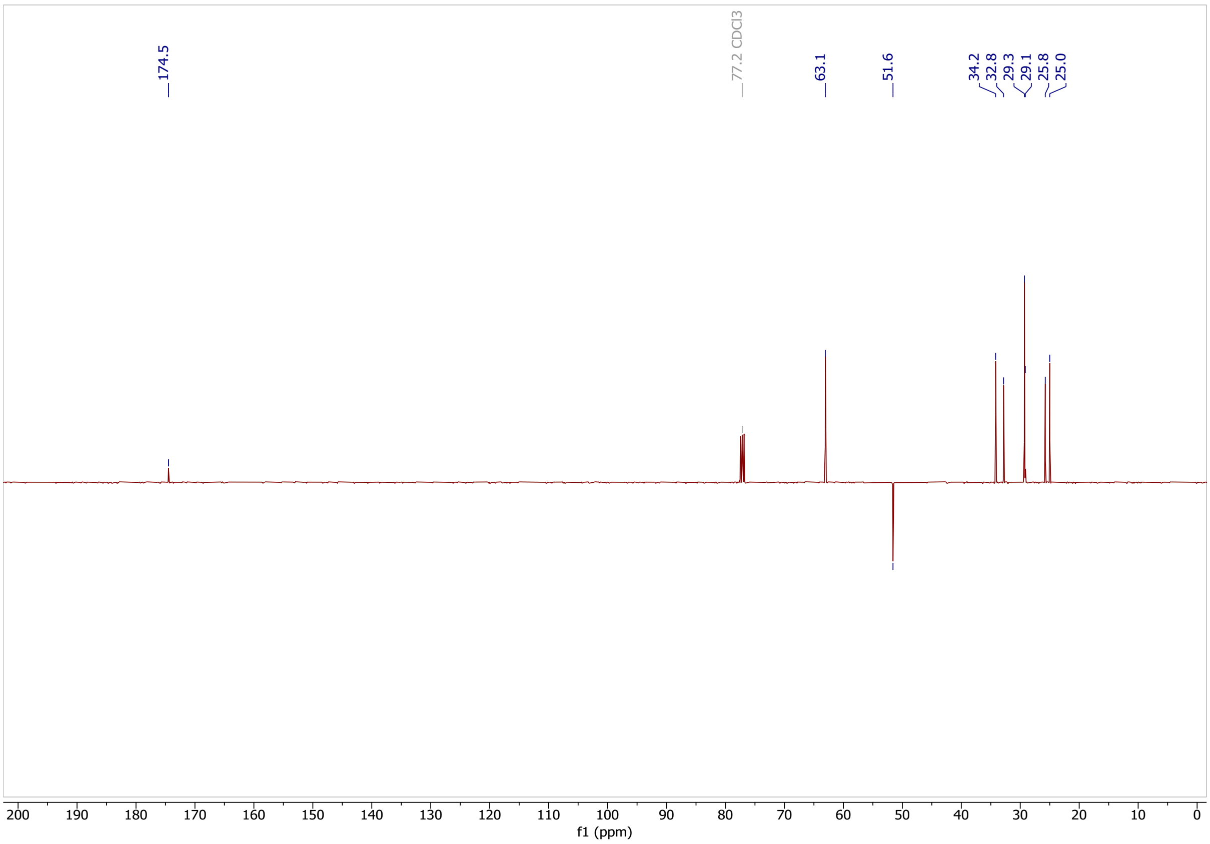

Classified as Internal | Intern

**methyl 9-oxononanoate (8)**

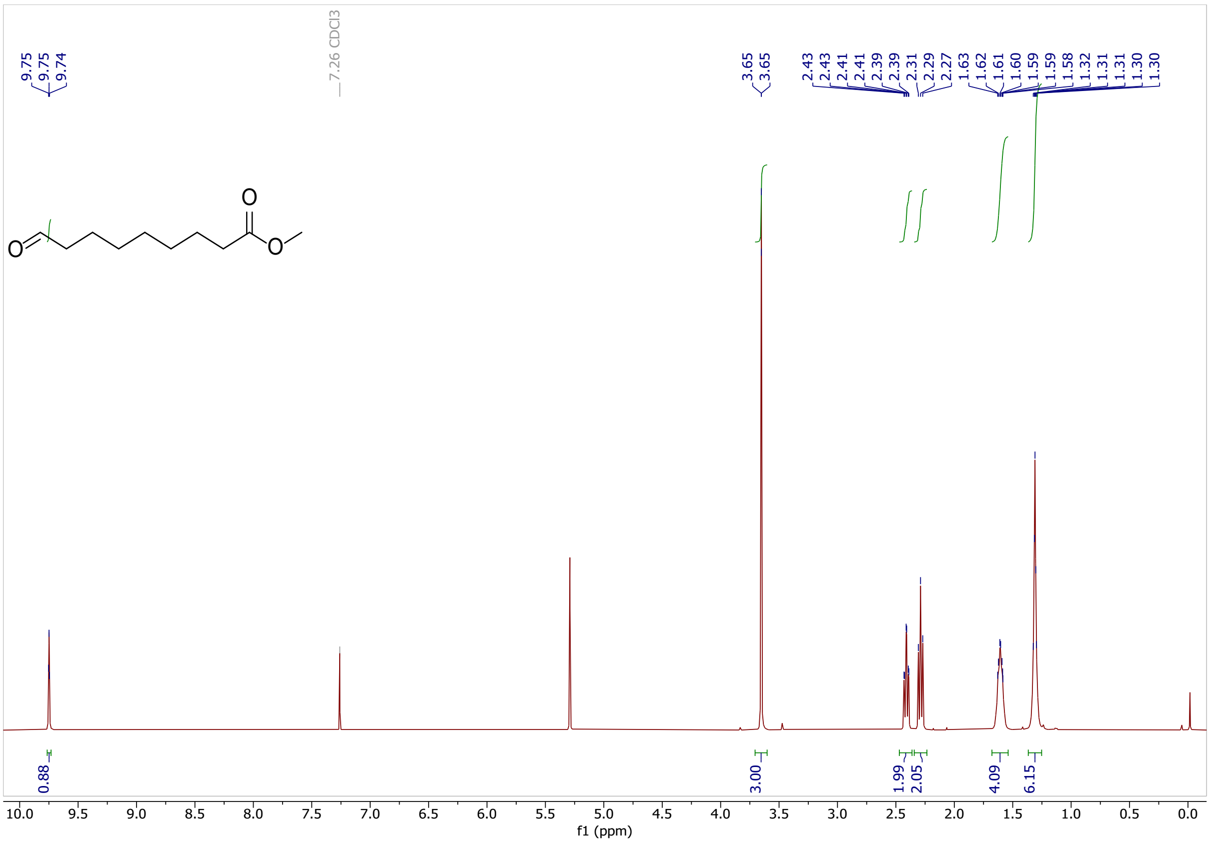

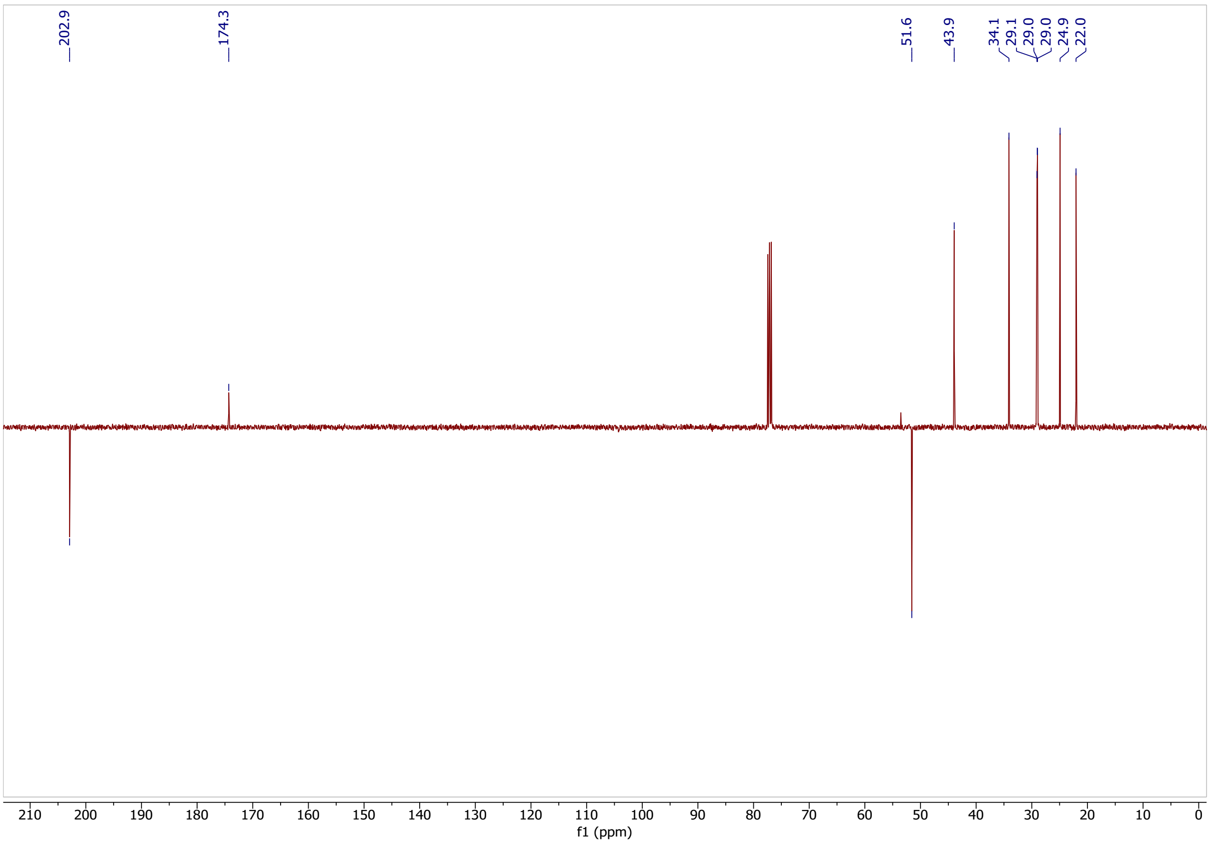

Classified as Internal | Intern

**methyl (Z)-11-(2-pentylcycloprop-1-en-1-yl)undec-9-enoate (18)**

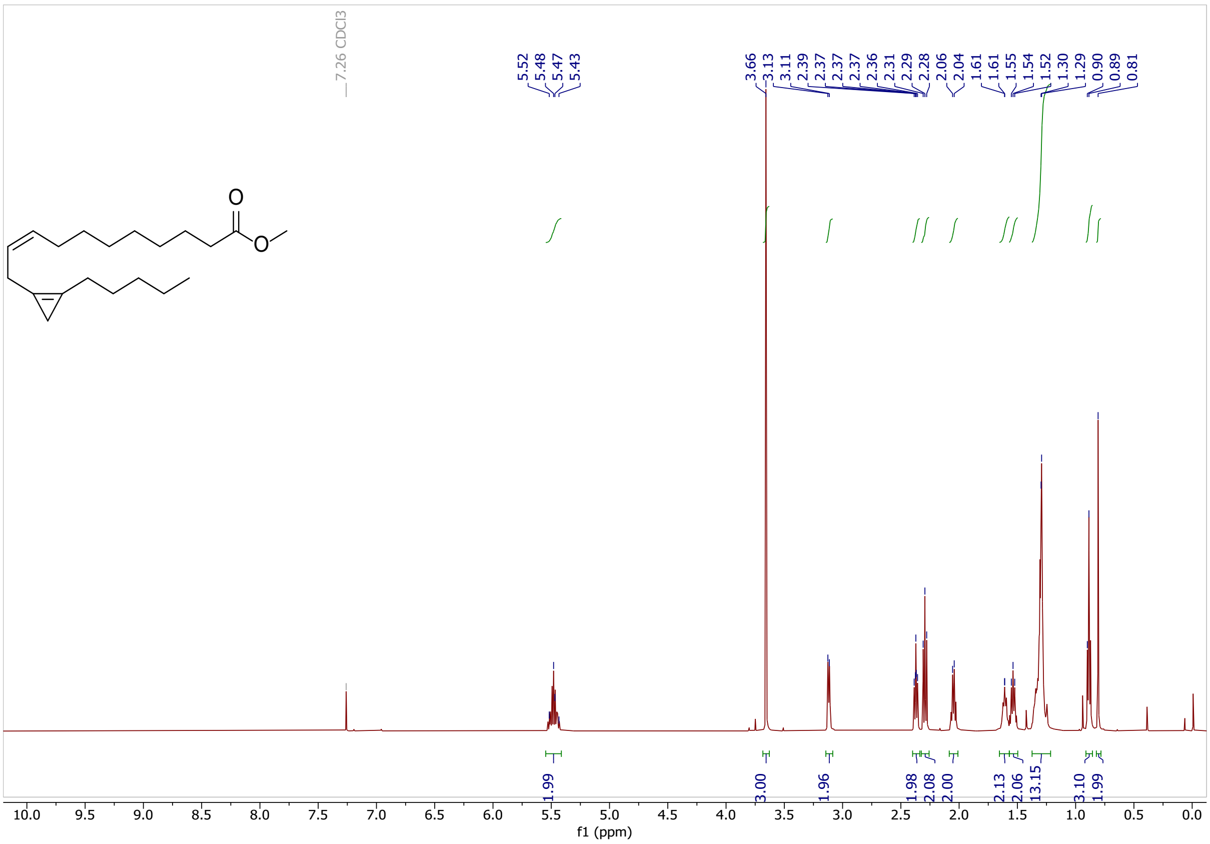

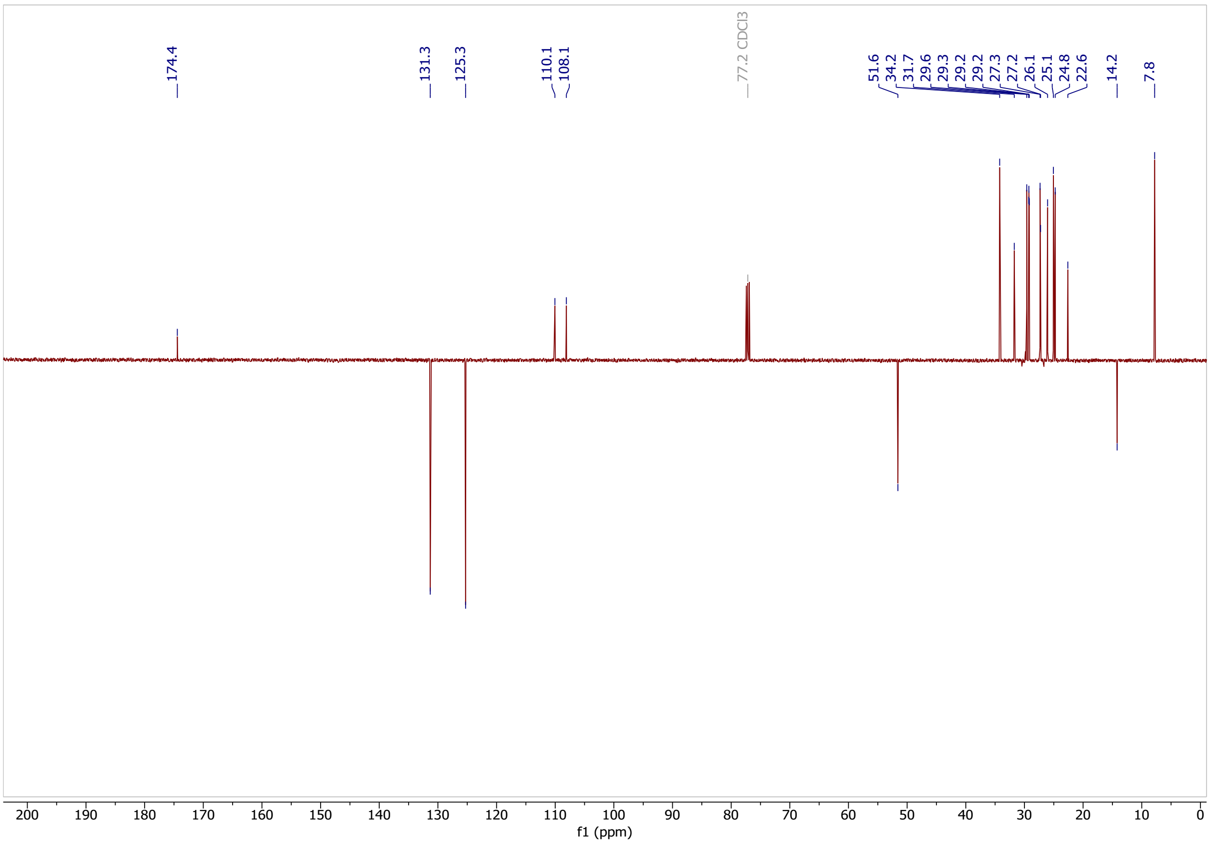

Classified as Internal | Intern

### (Z)-11-(2-pentylcycloprop-1-en-1-yl)undec-9-enoic acid (5)

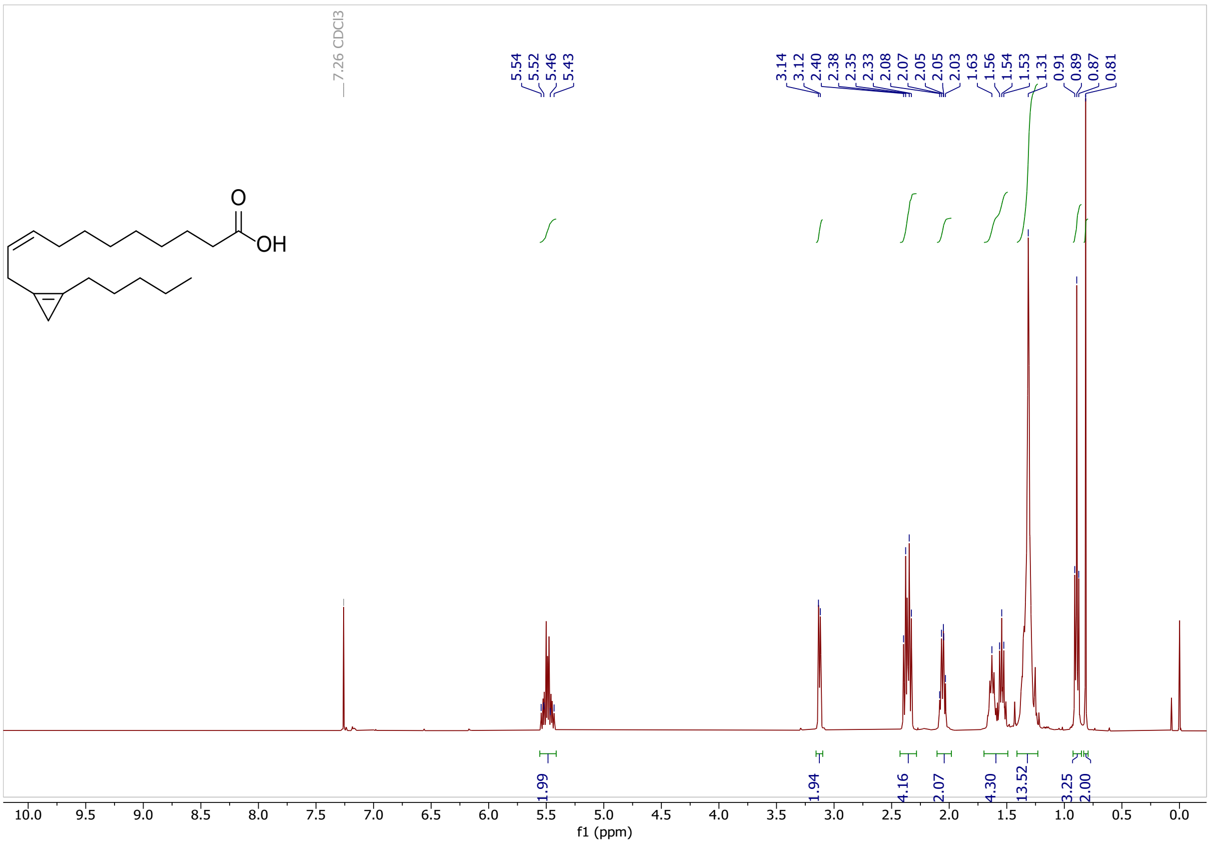

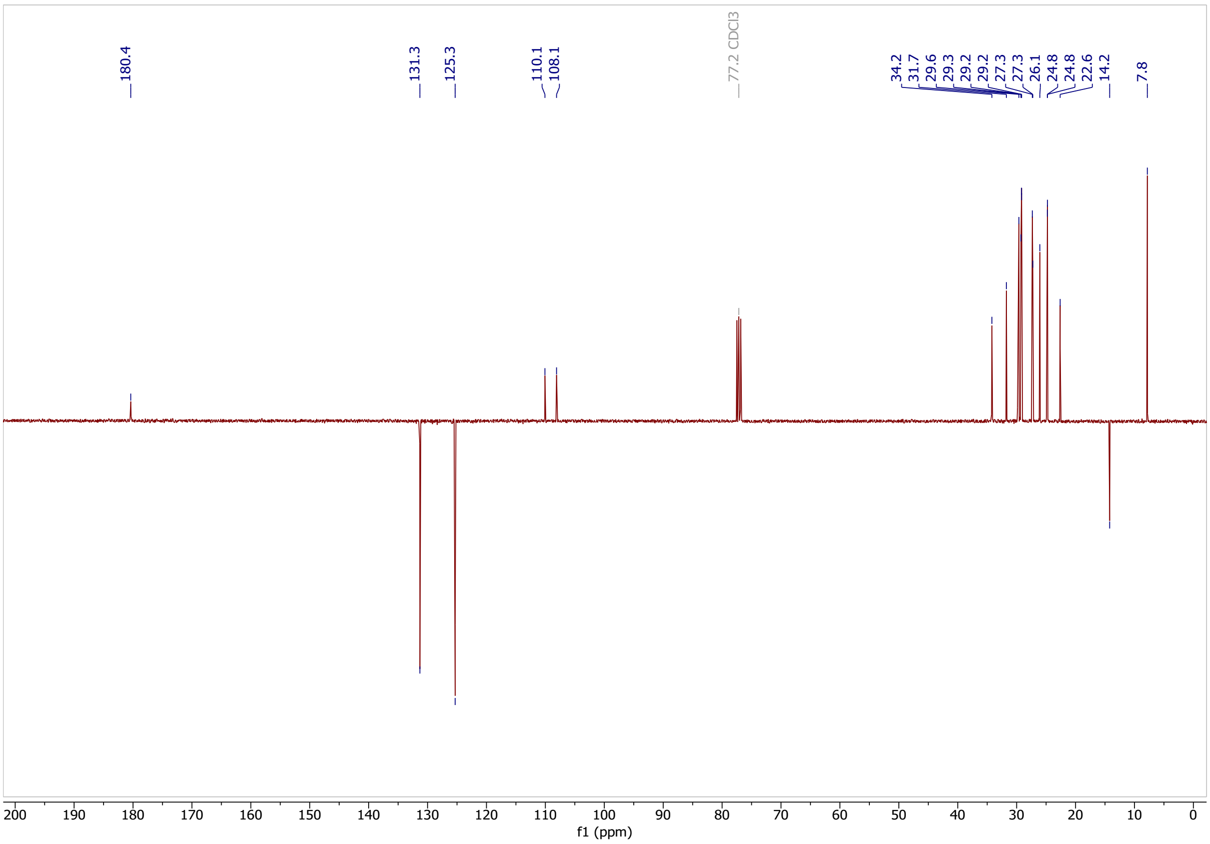

Classified as Internal | Intern

**methyl 7-hydroxyhept-5-ynoate (19)**

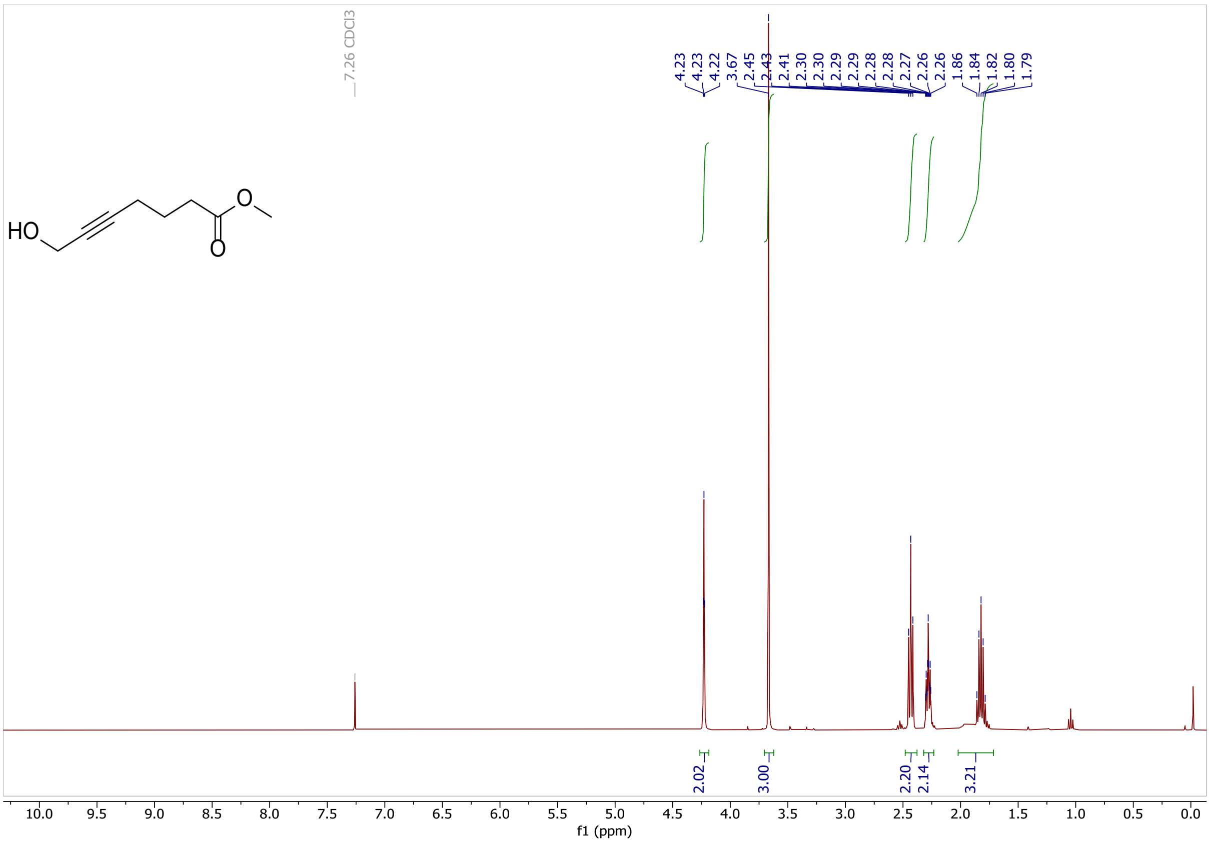

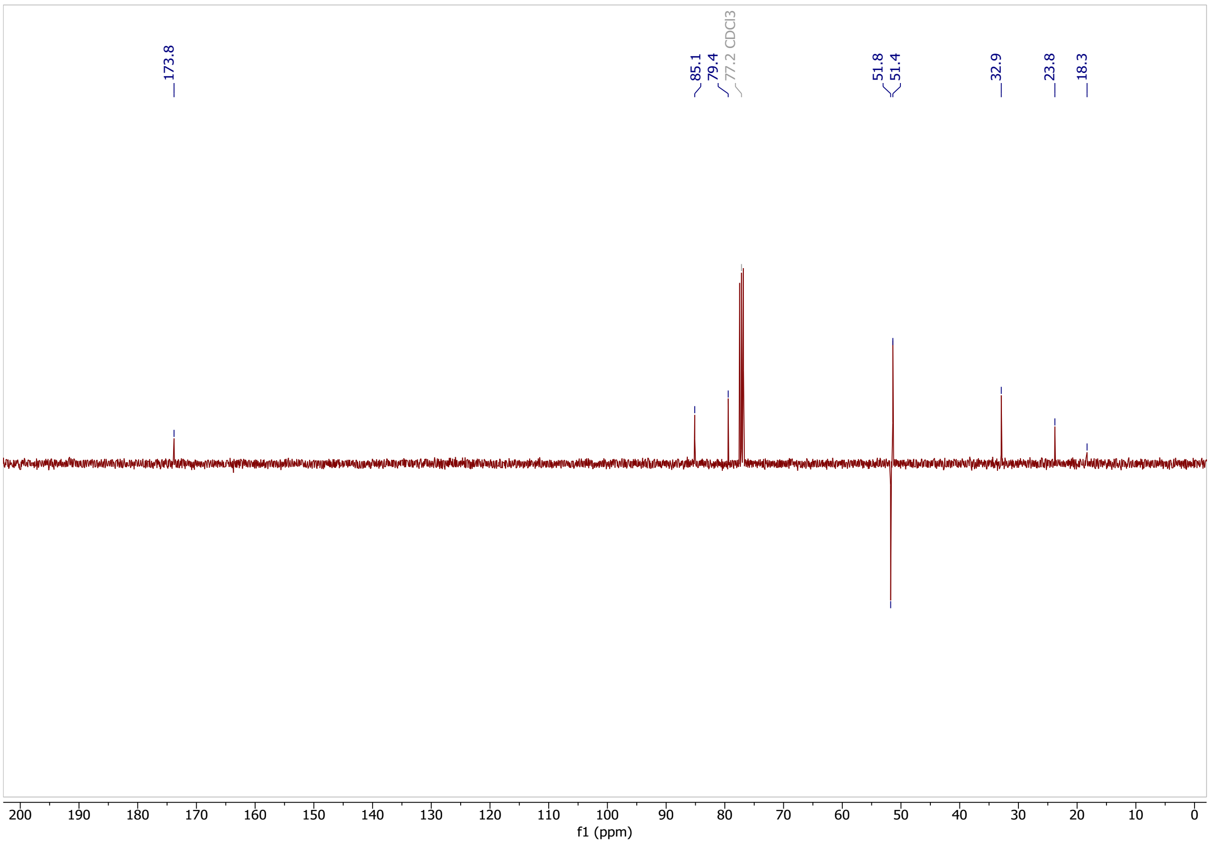

Classified as Internal | Intern

**methyl 7-bromohept-5-ynoate (20)**

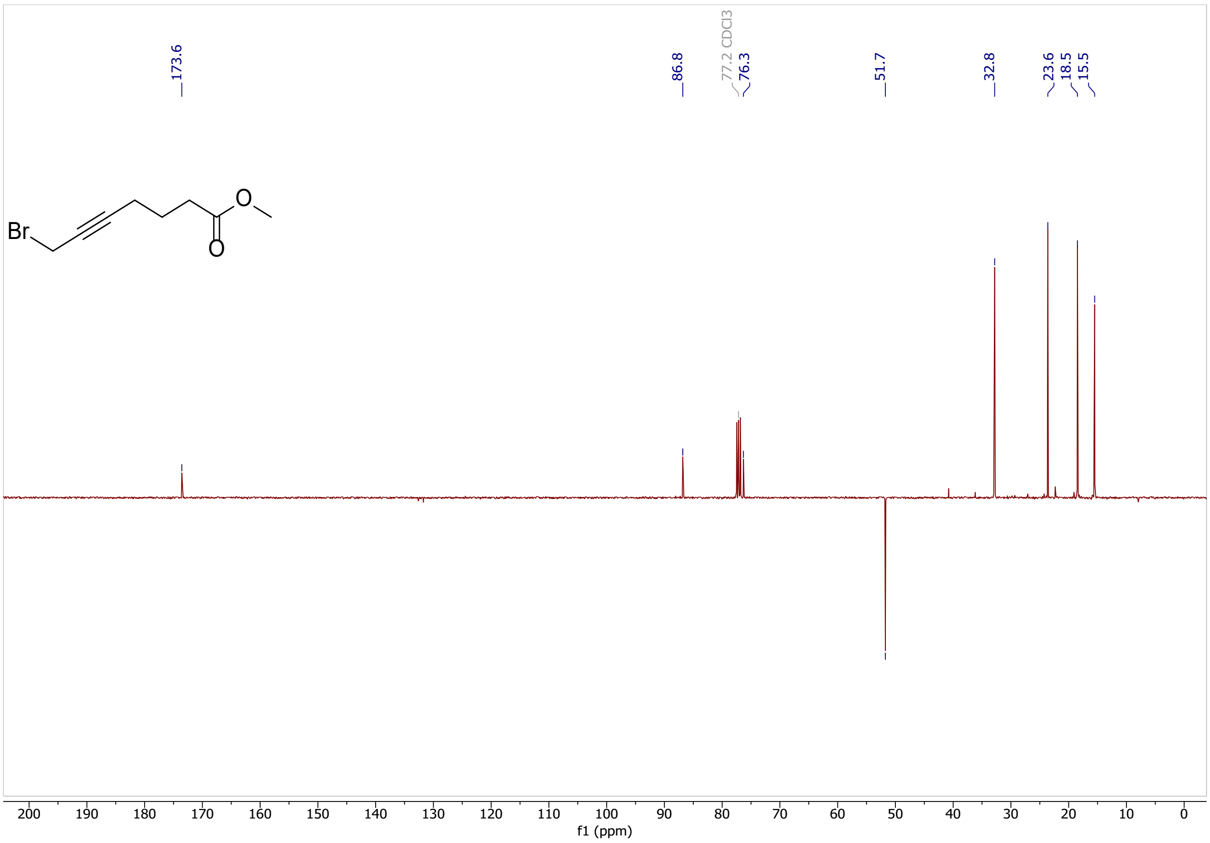

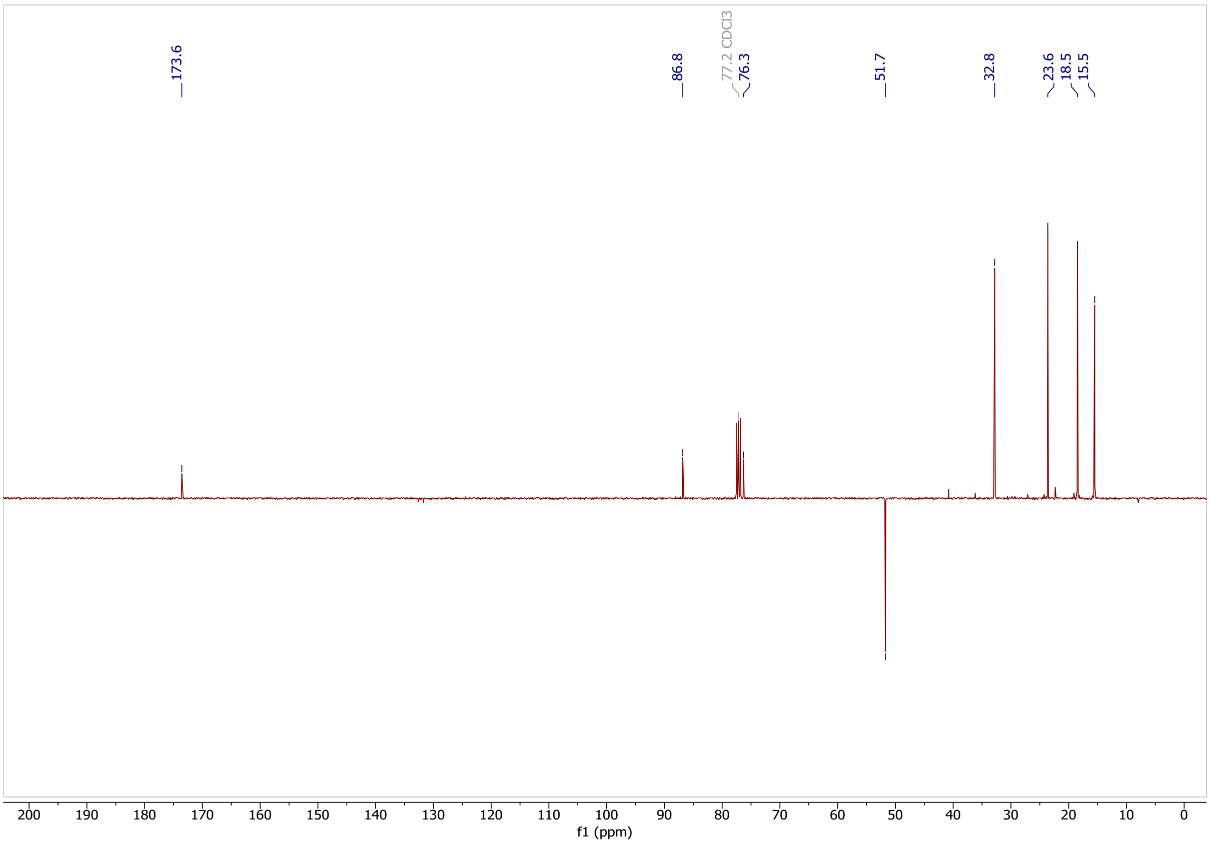

Classified as Internal | Intern

### methyl 11-hydroxyundeca-5,8-diynoate (10)

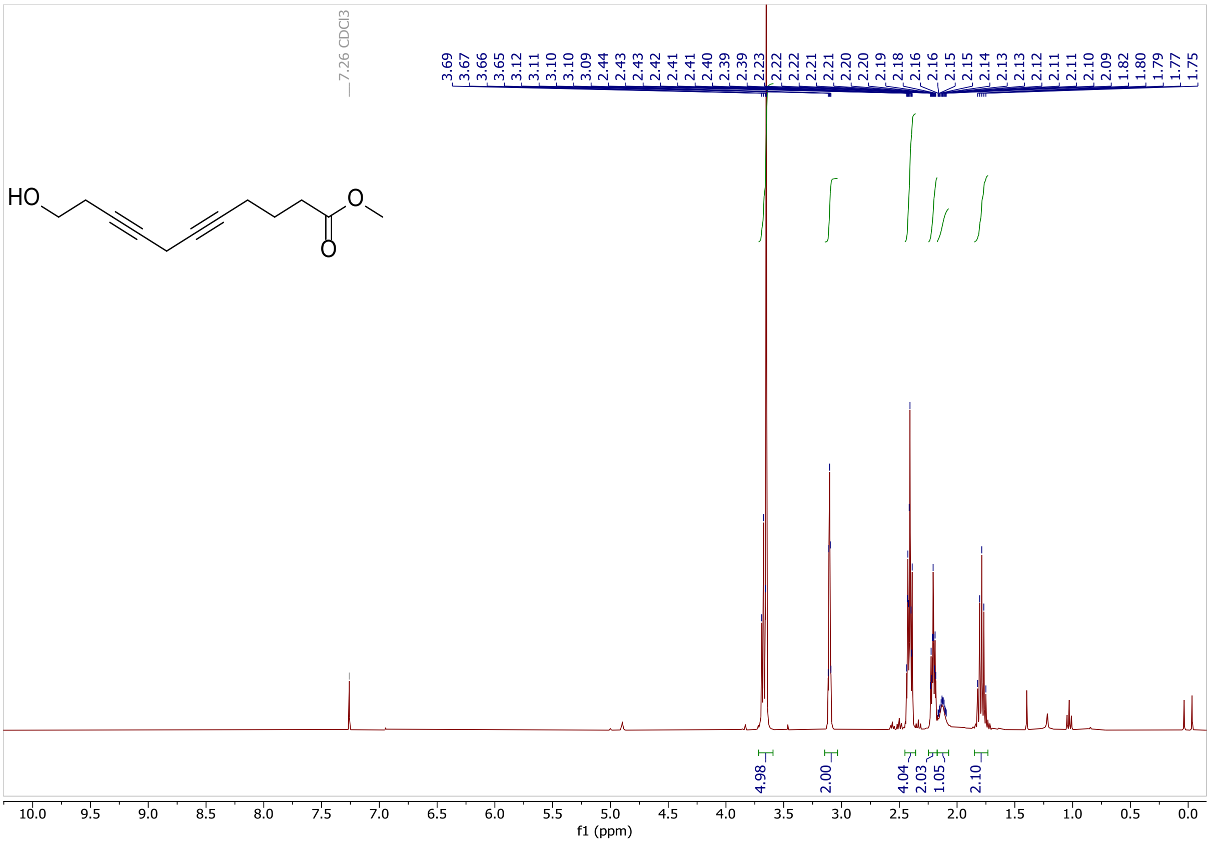

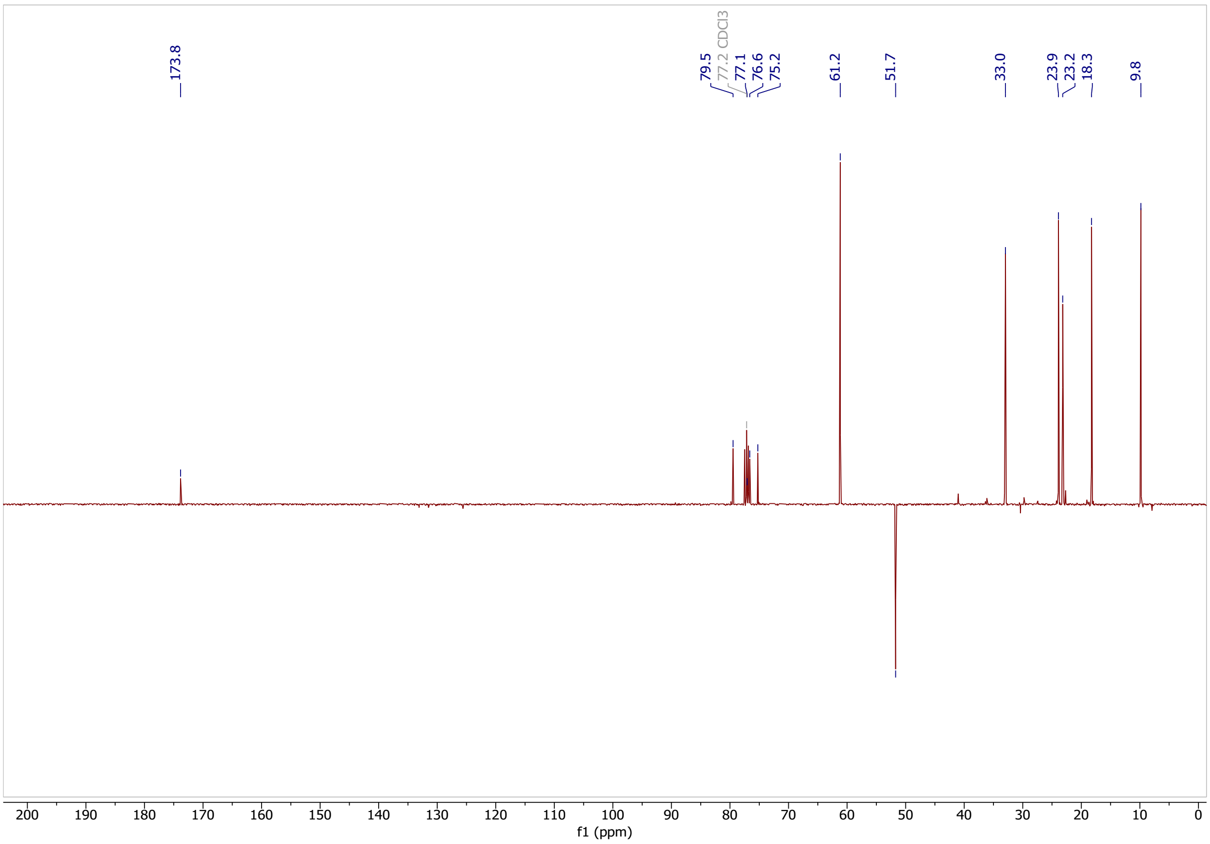

Classified as Internal | Intern

**methyl (5Z,8Z)-11-hydroxyundeca-5,8-dienoate (21)**

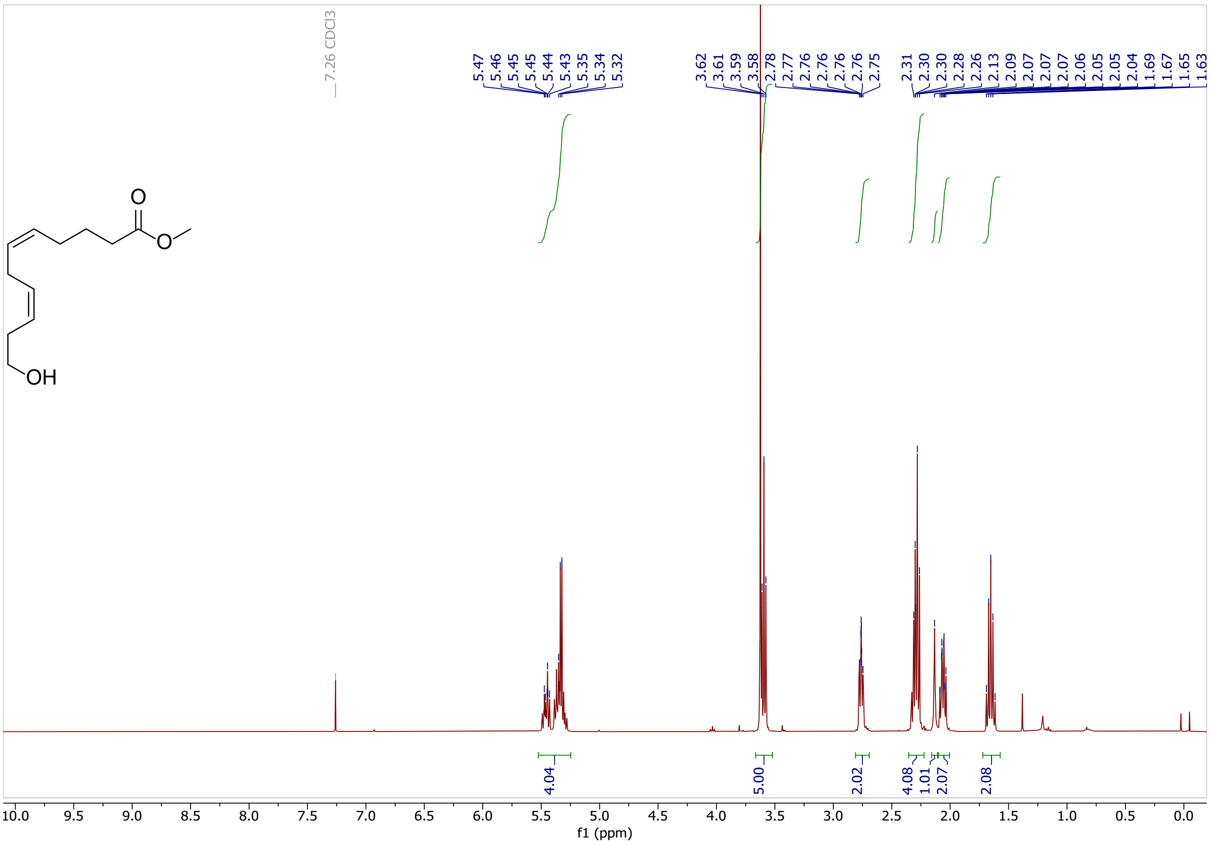

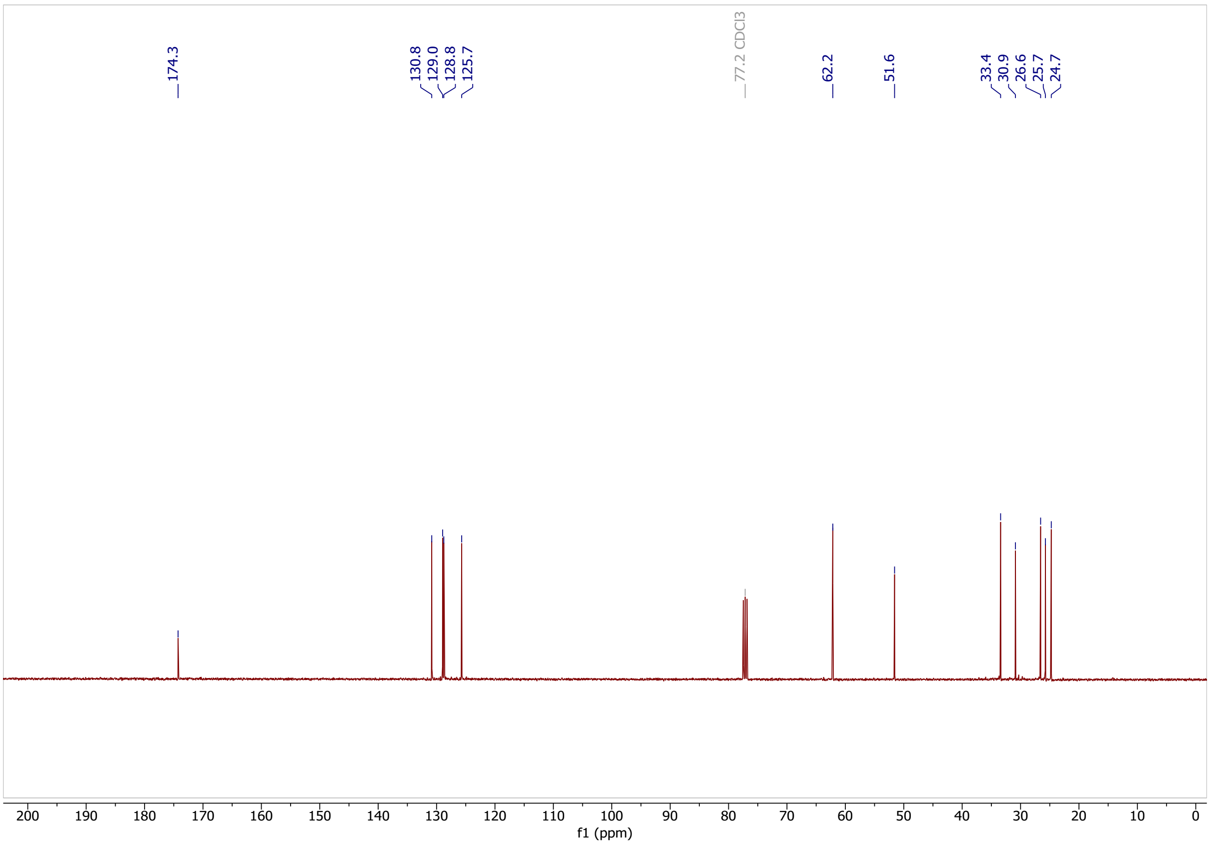

Classified as Internal | Intern

### methyl (5Z,8Z)-11-oxoundeca-5,8-dienoate (9)

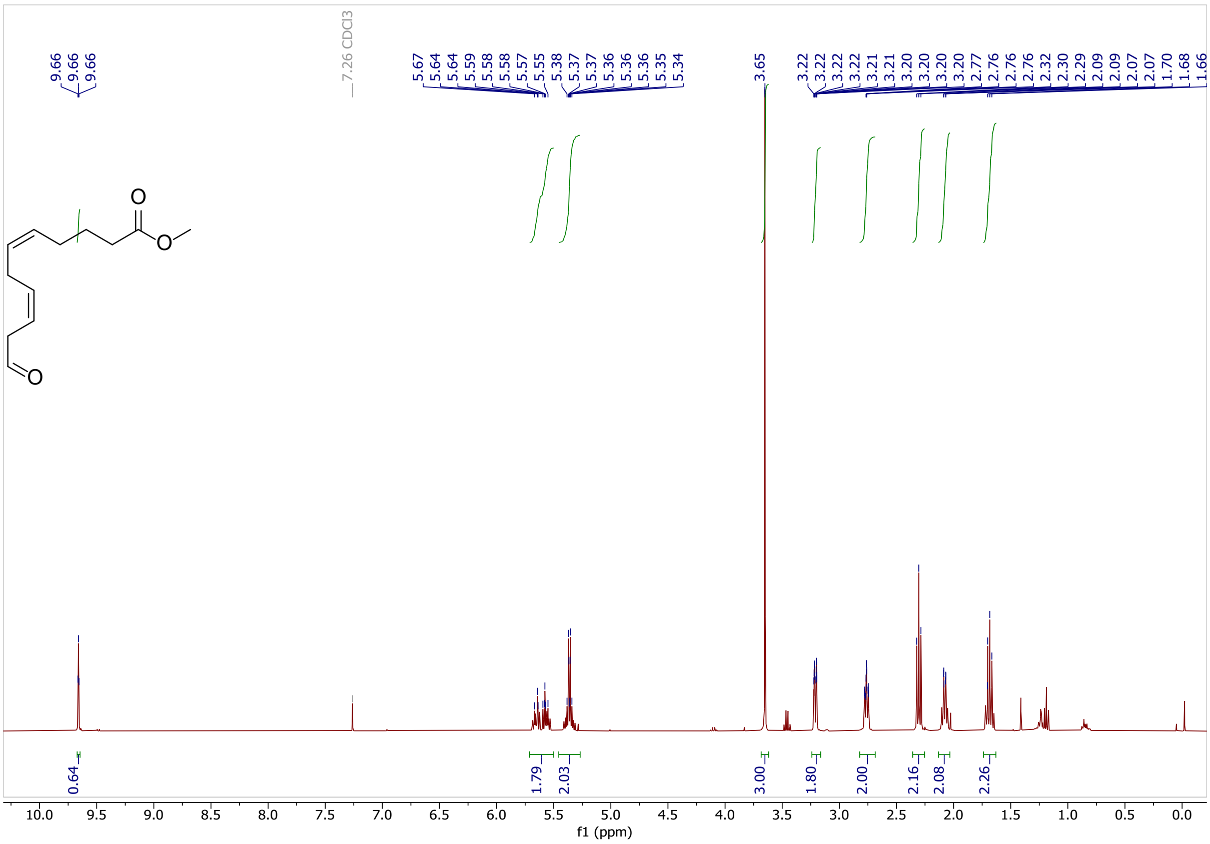

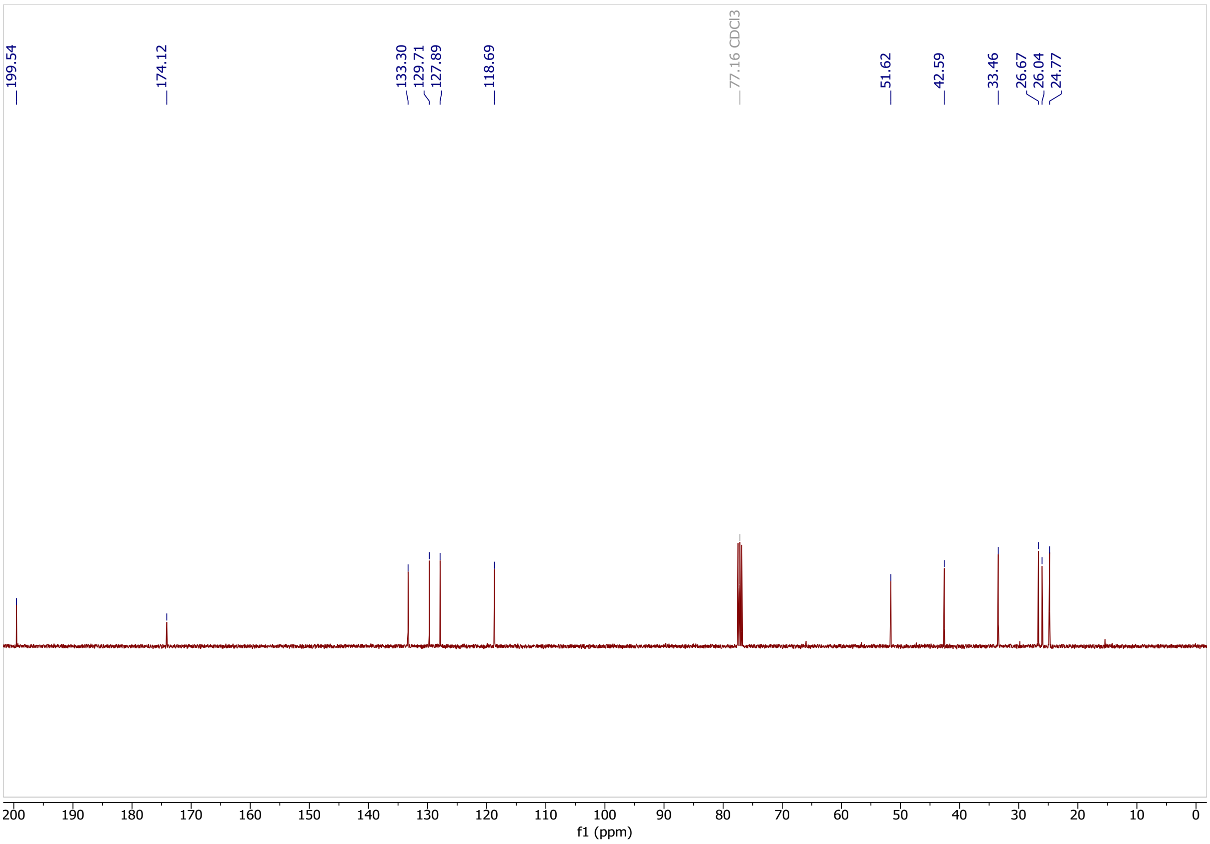

Classified as Internal | Intern

**methyl (5Z,8Z,11Z)-13-(2-pentylcycloprop-1-en-1-yl)trideca-5,8,11-trienoate (22)**

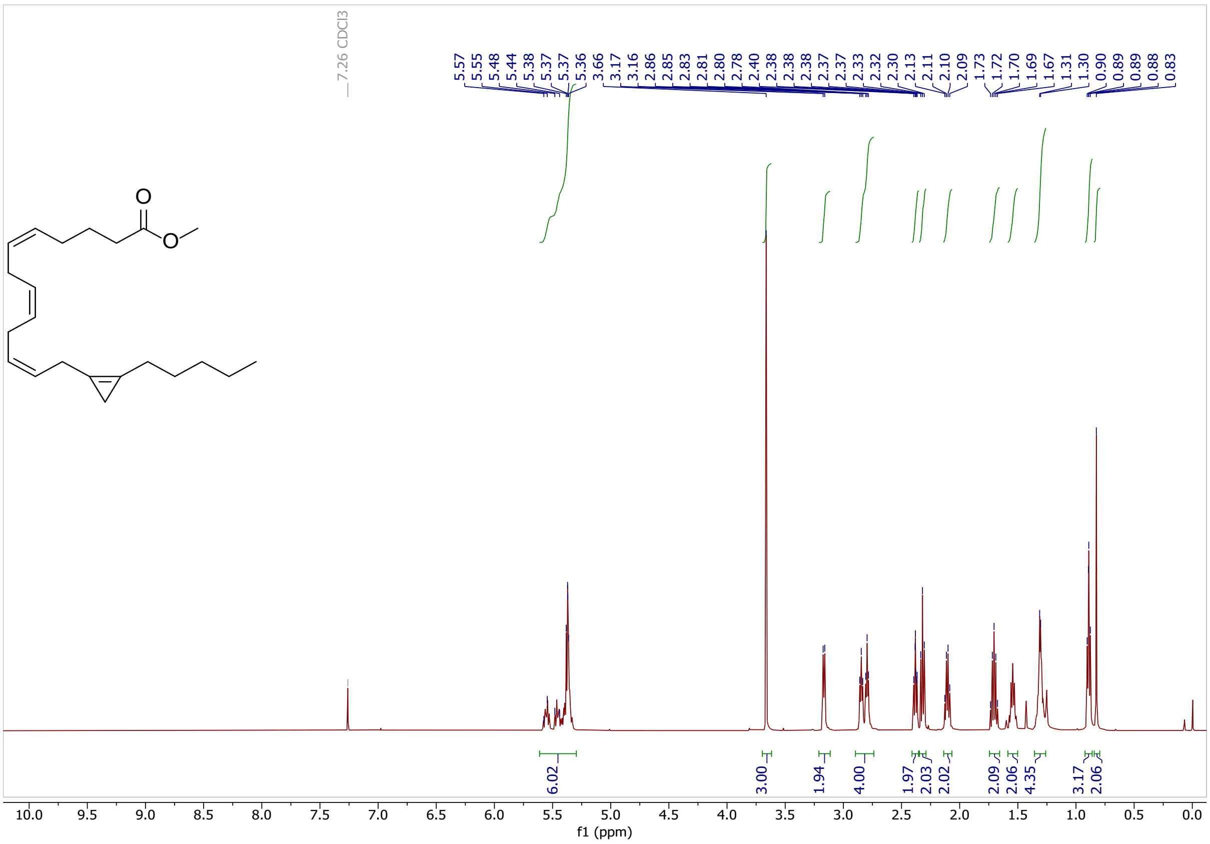

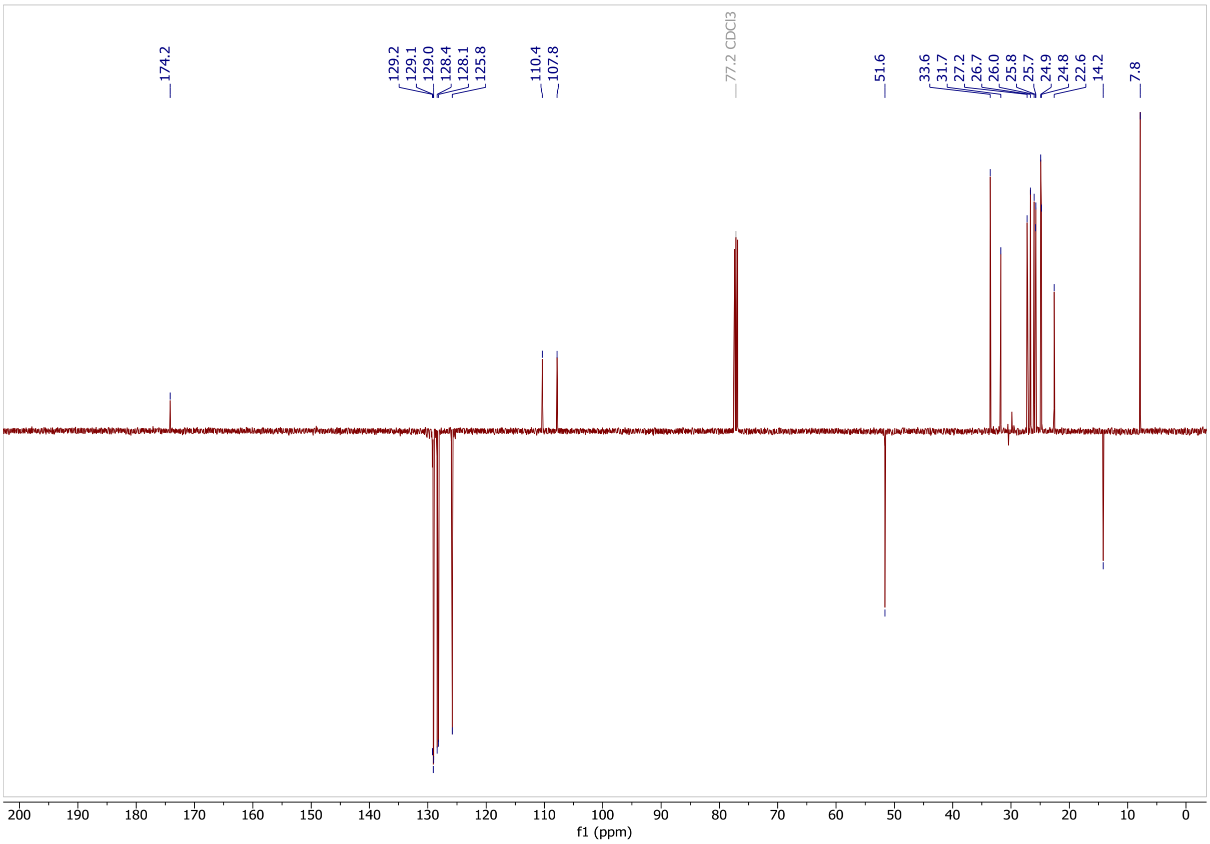

Classified as Internal | Intern

### (5Z,8Z,11Z)-13-(2-pentylcycloprop-1-en-1-yl)trideca-5,8,11-trienoic acid (6)

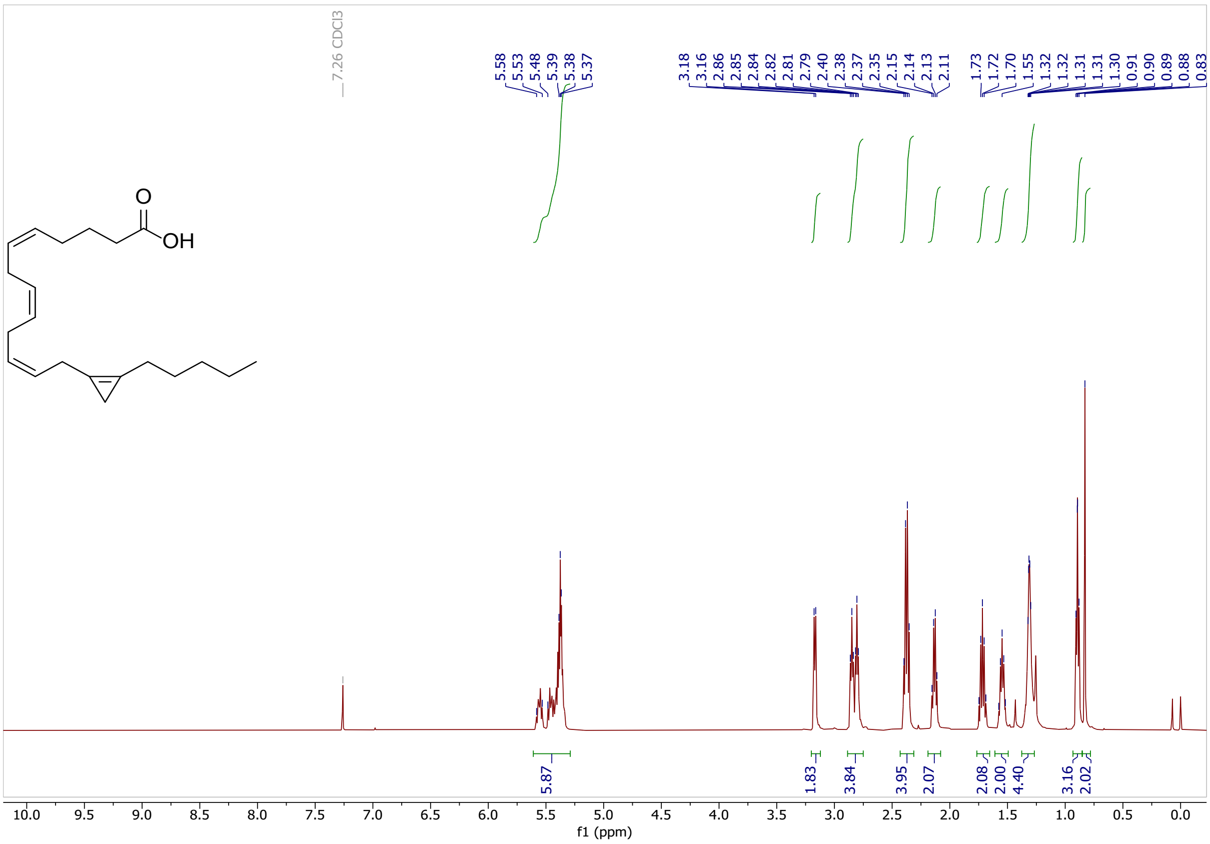

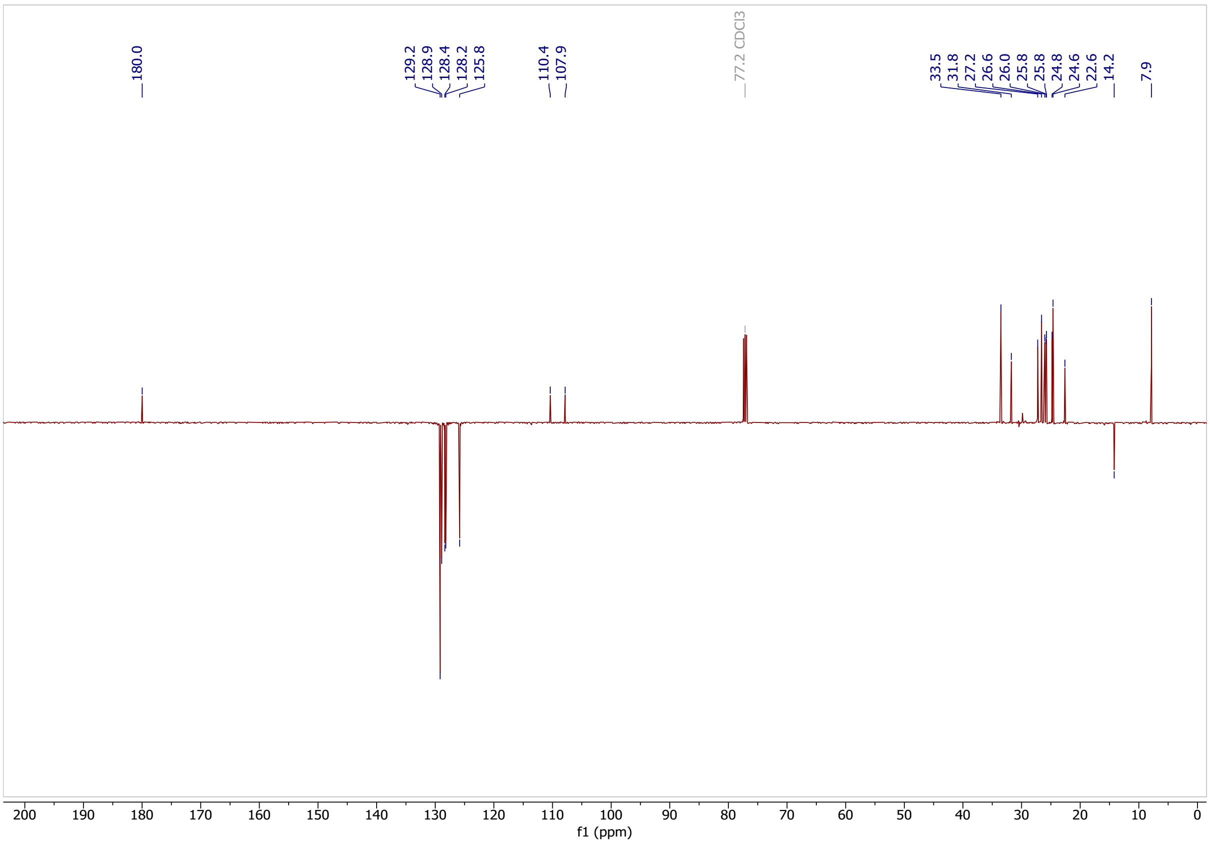

Classified as Internal | Intern

### 8-(2-hexylcycloprop-1-en-1-yl)octanoic acid (3)

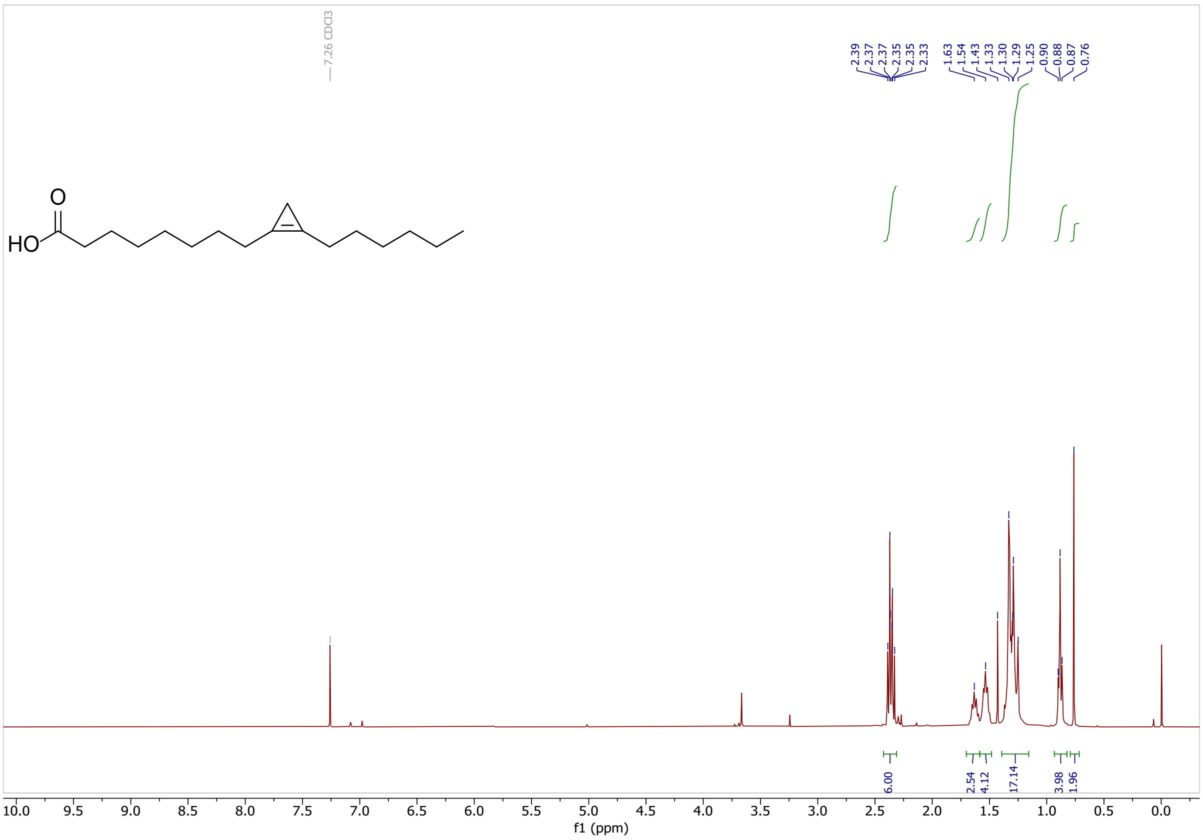

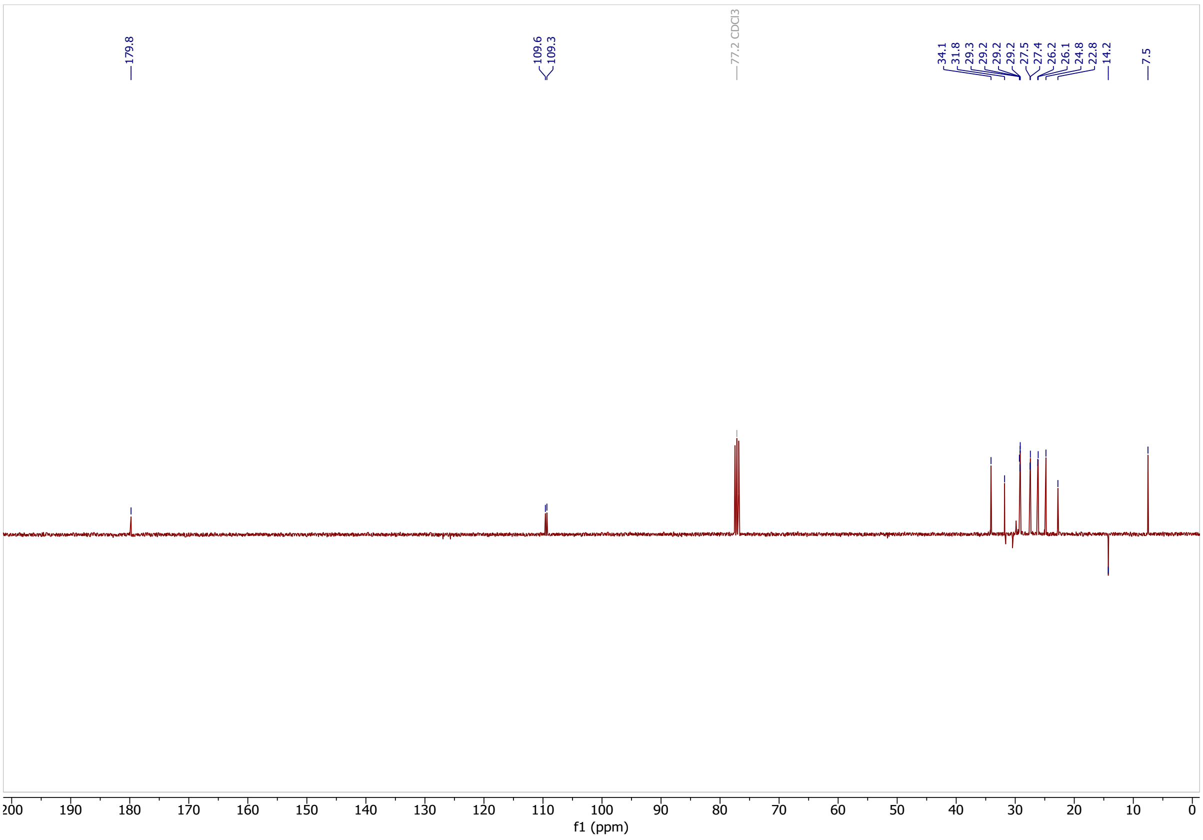

Classified as Internal | Intern

**1,1,2-tribromo-2-methylcyclopropane (25)**

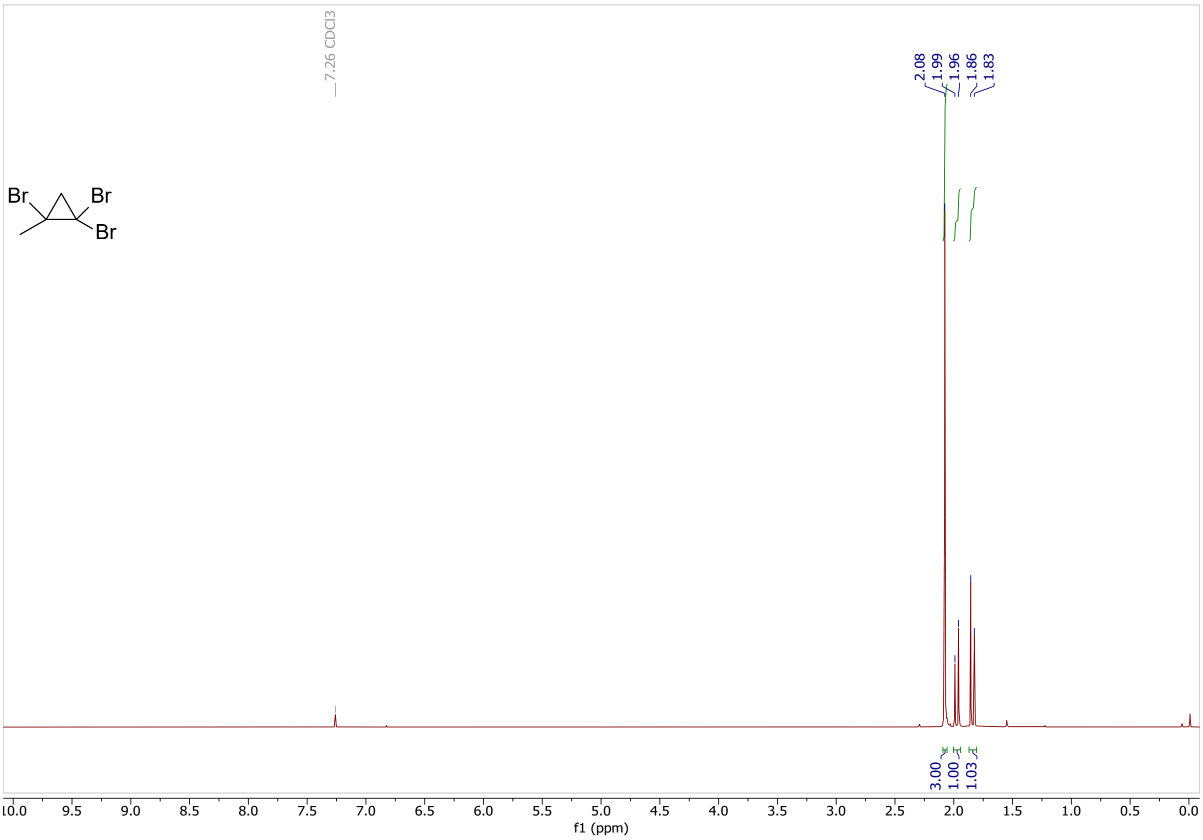

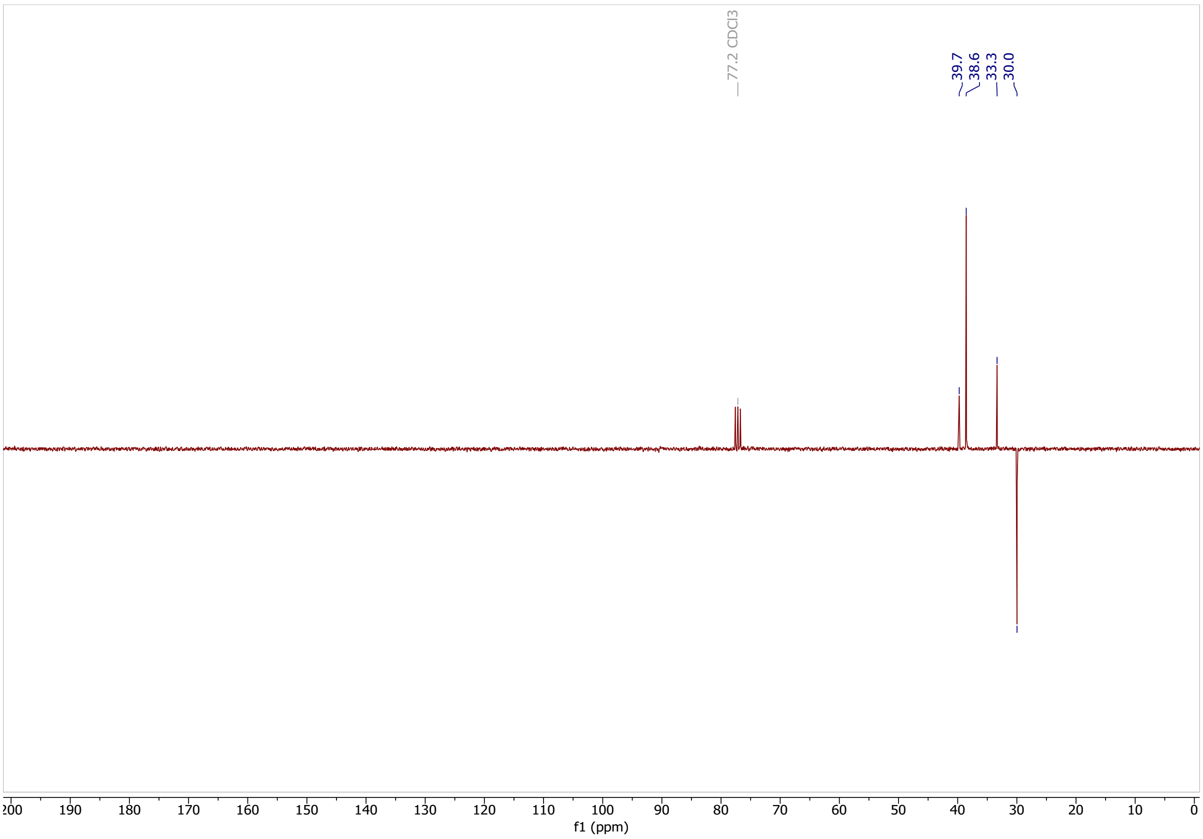

Classified as Internal | Intern

### 1-(12-iodododecyl)-2-methylcycloprop-1-ene (27)

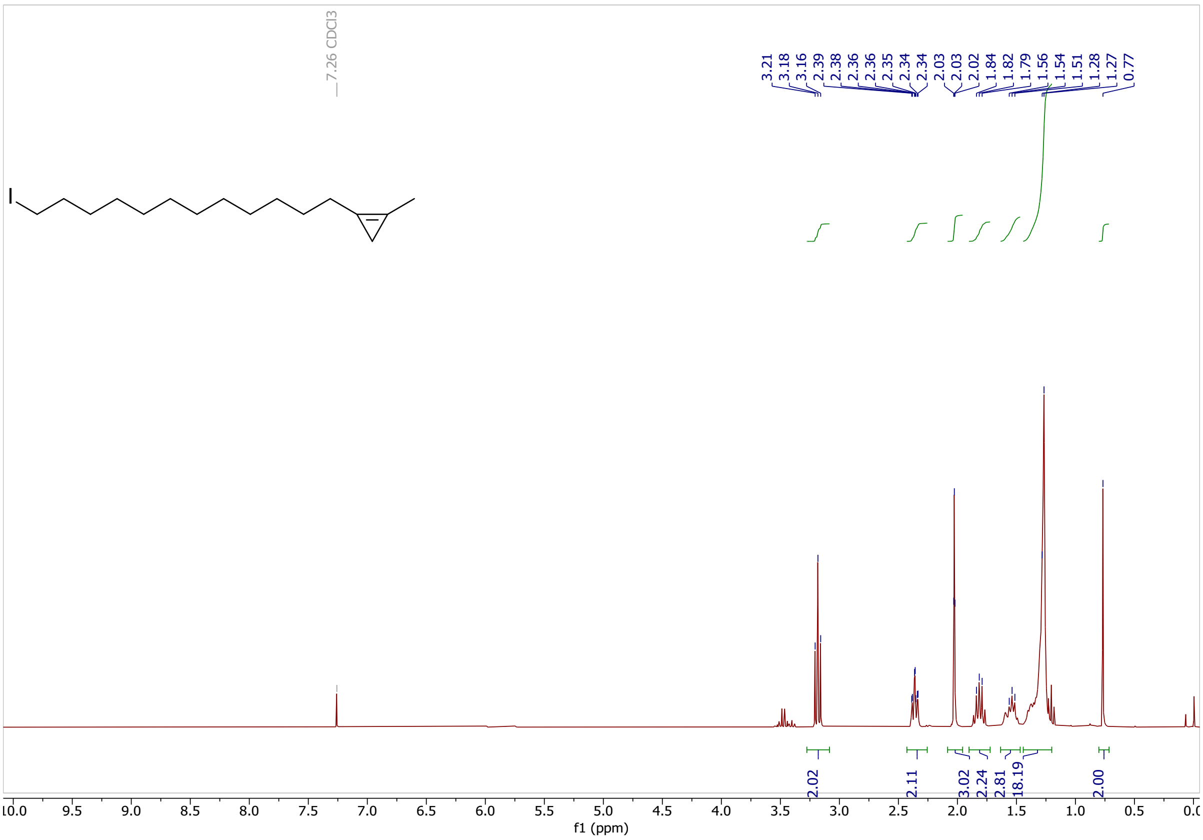

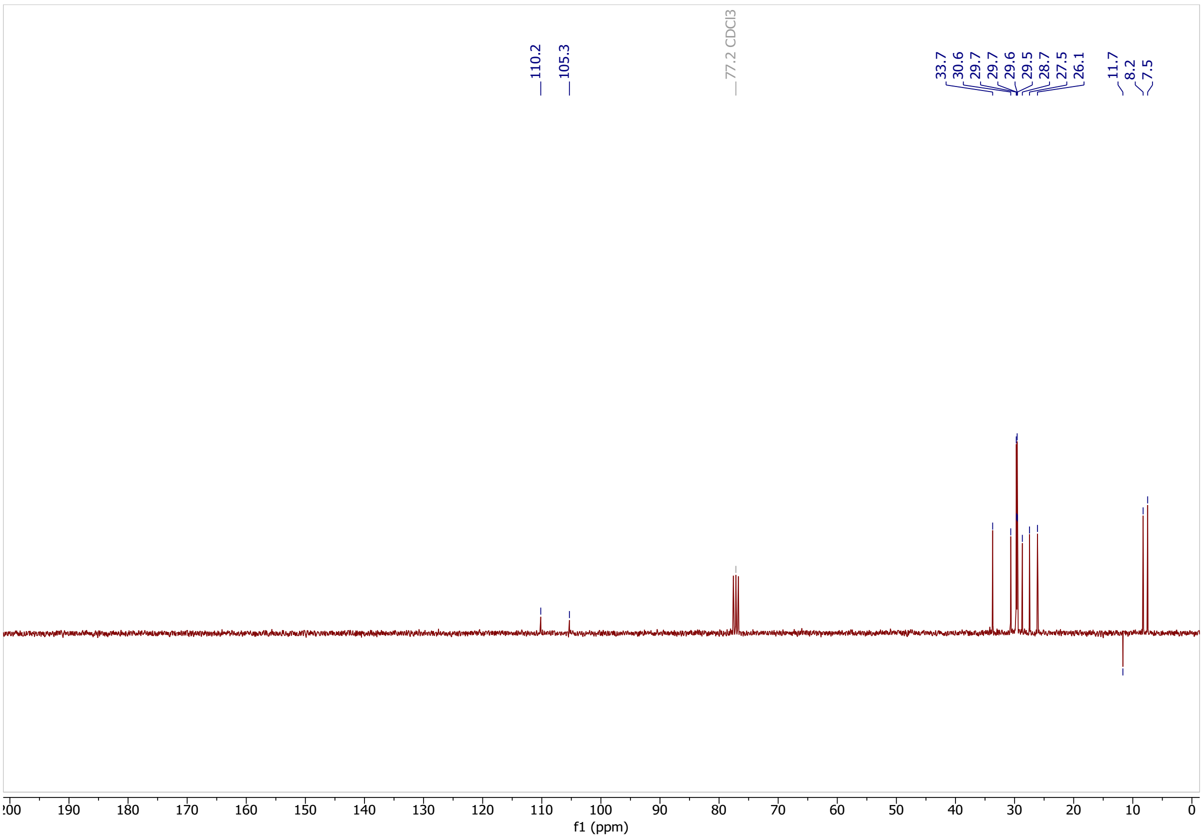

Classified as Internal | Intern

**13-(2-methylcycloprop-1-en-1-yl)tridecanenitrile (30)**

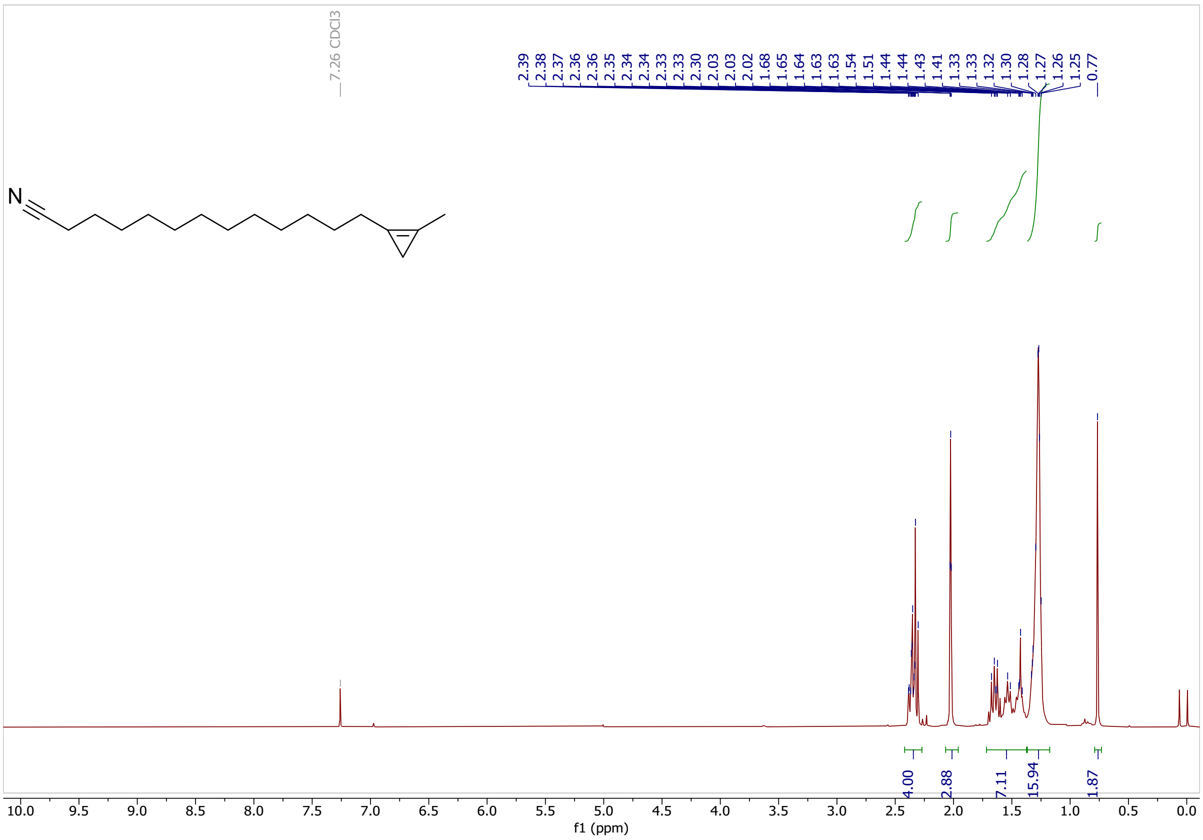

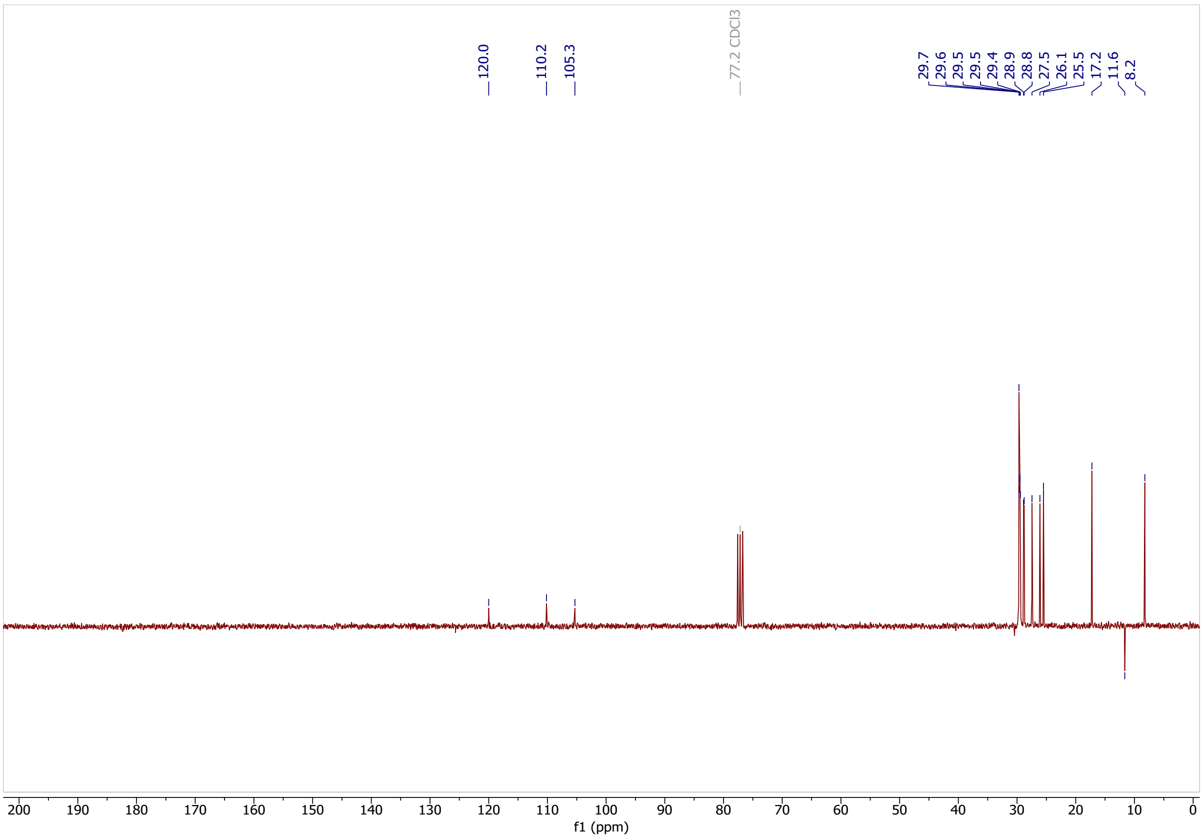

Classified as Internal | Intern

### 13-(2-methylcycloprop-1-en-1-yl)tridecanoic acid (1)

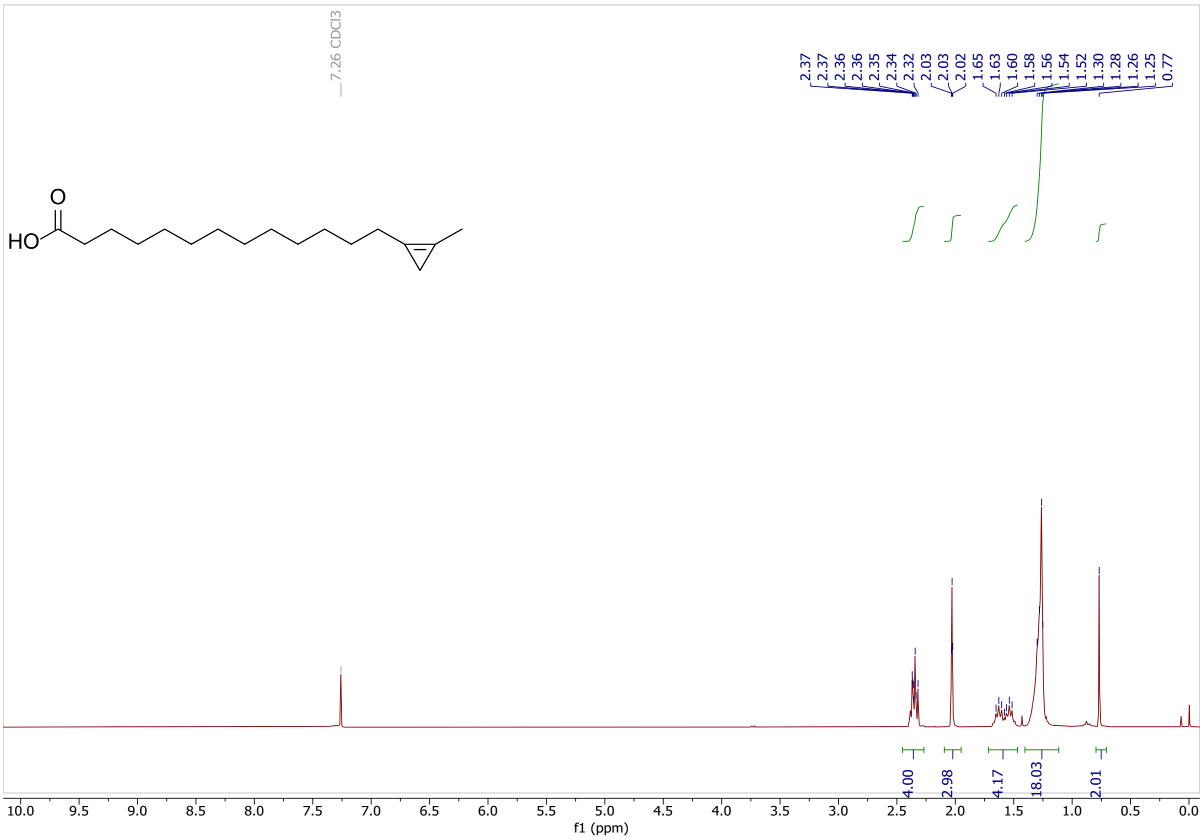

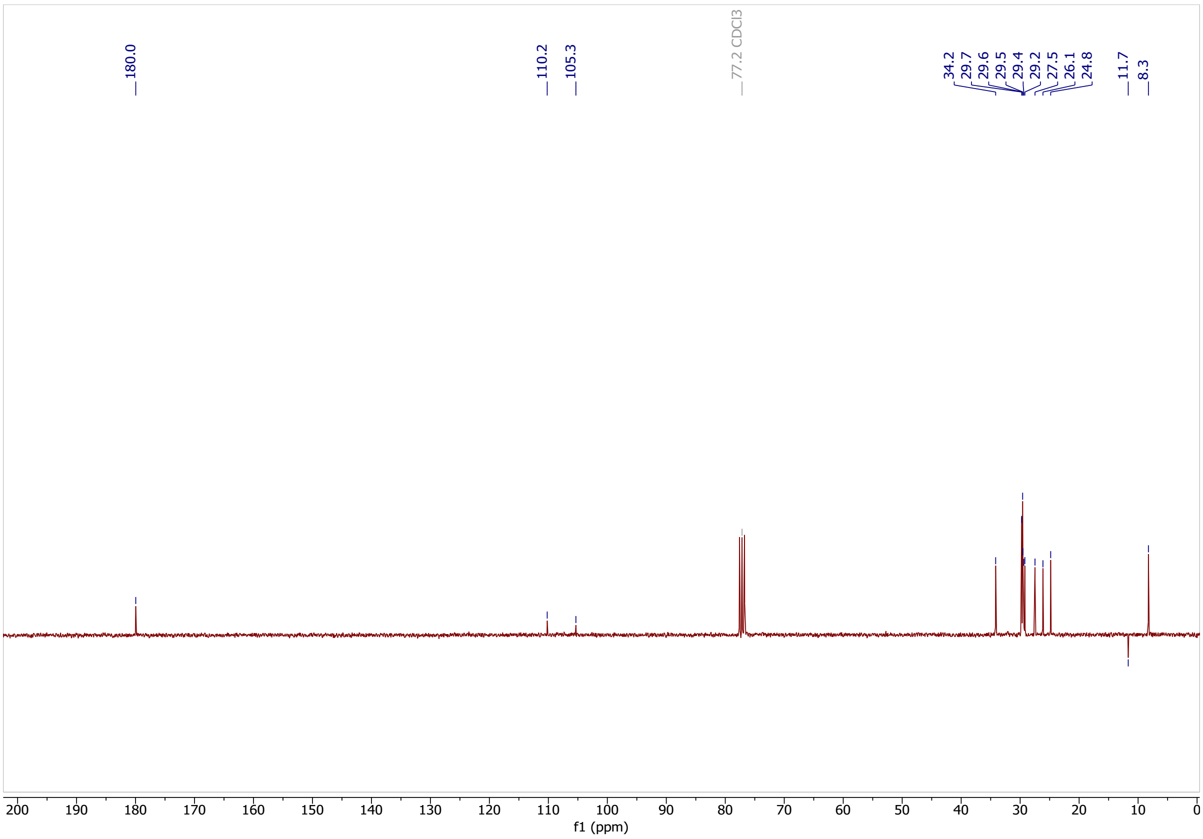

Classified as Internal | Intern

**1-(14-iodotetradecyl)-2-methylcycloprop-1-ene (28)**

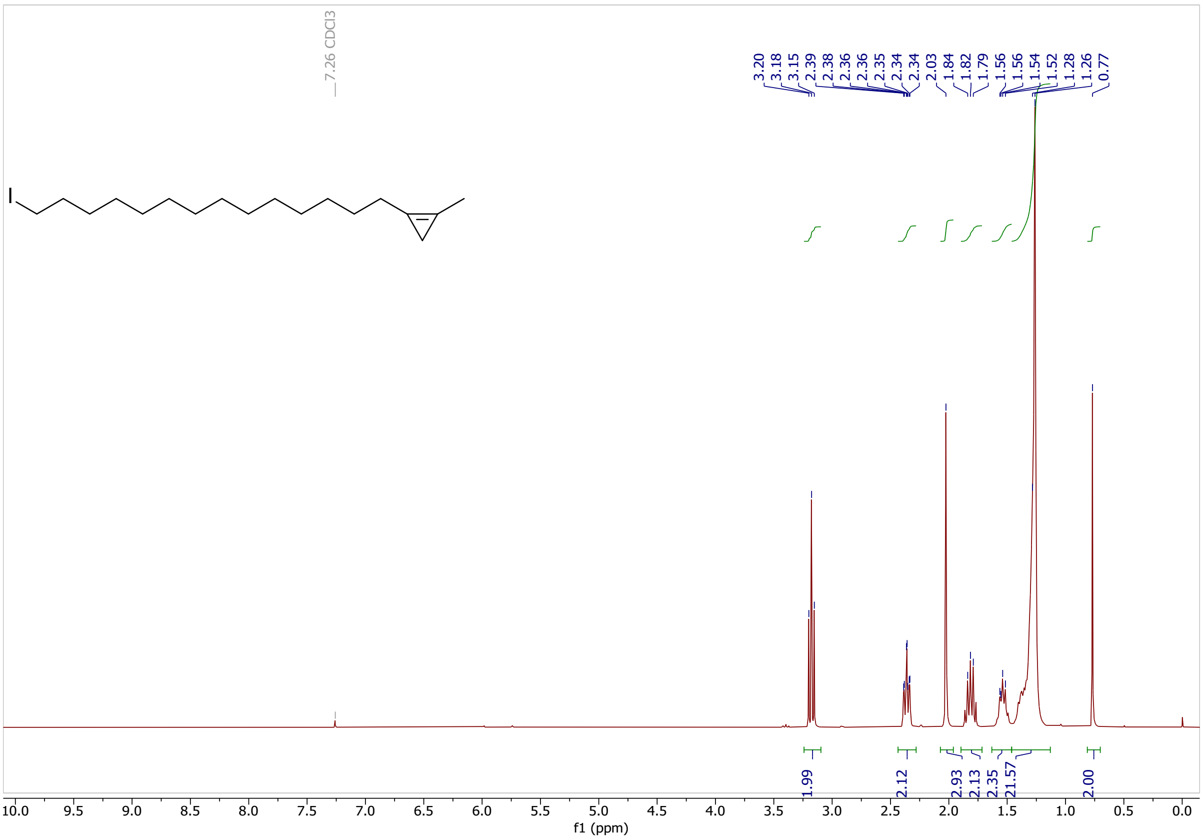

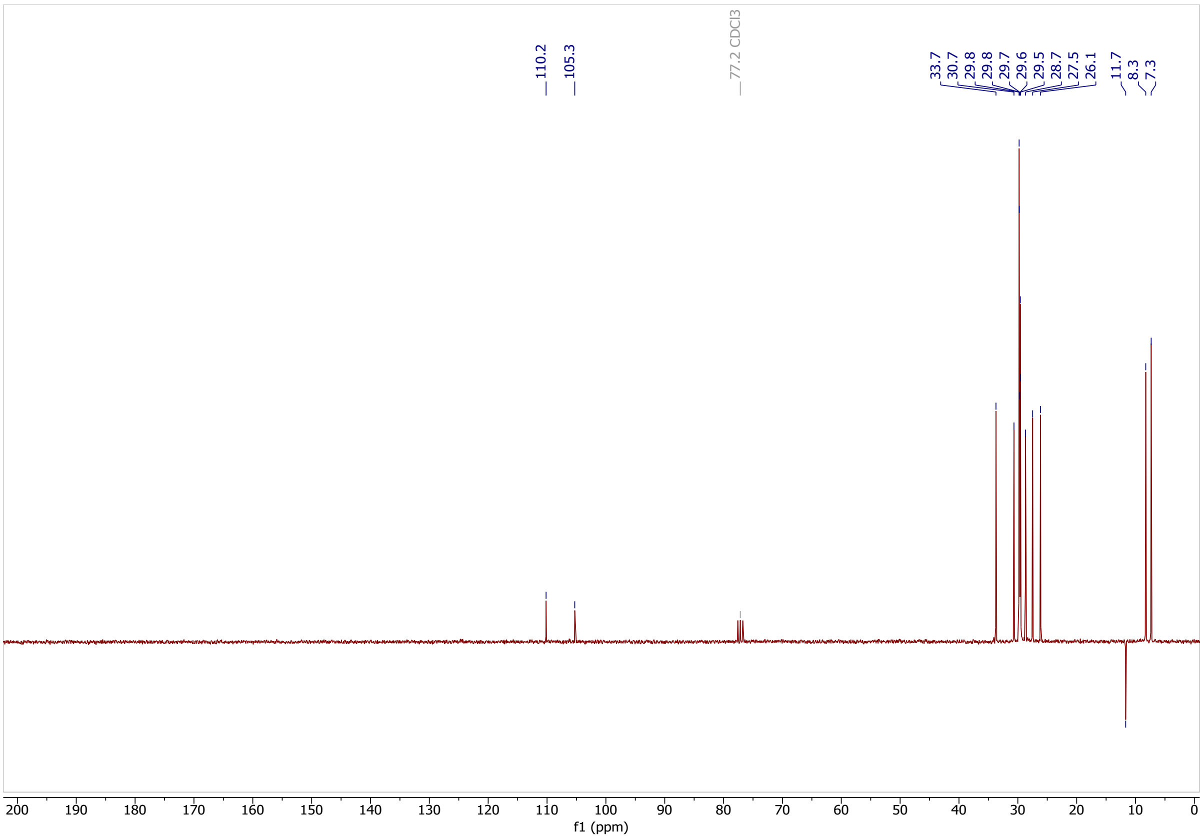

Classified as Internal | Intern

**15-(2-methylcycloprop-1-en-1-yl)pentadecanenitrile (31)**

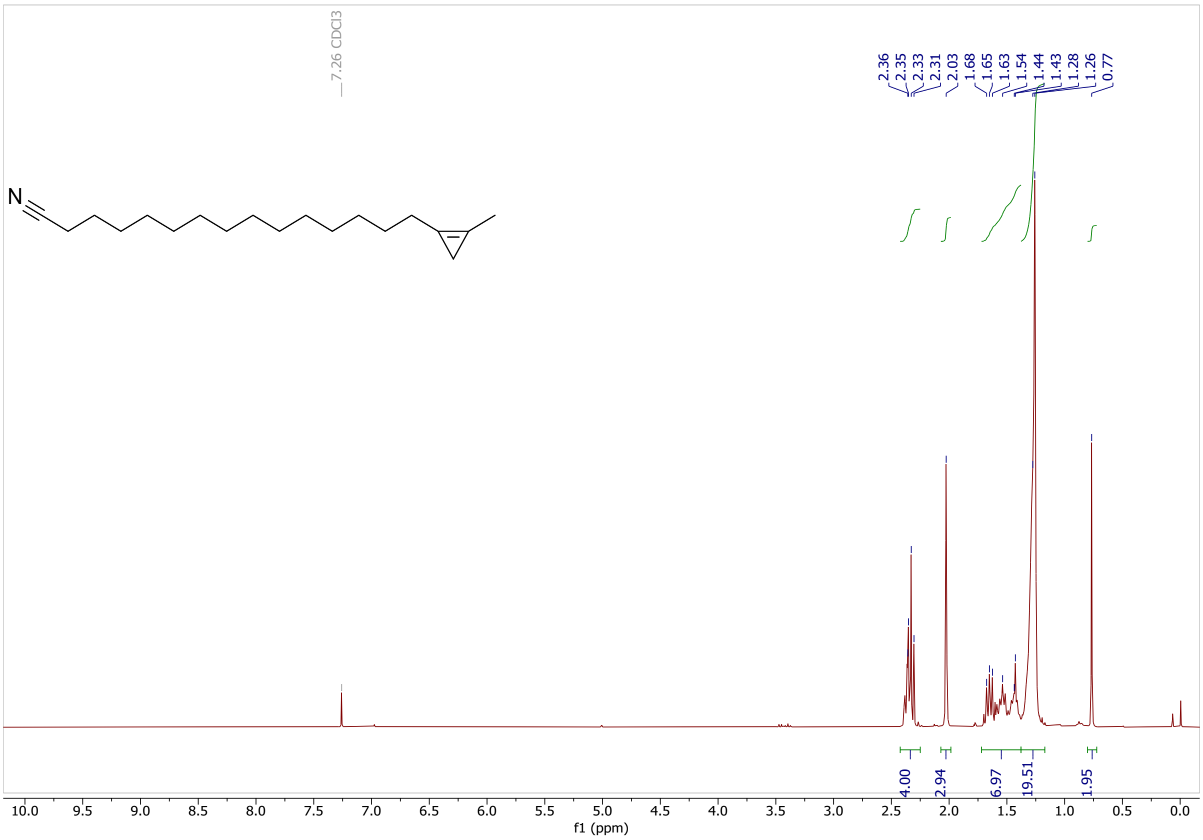

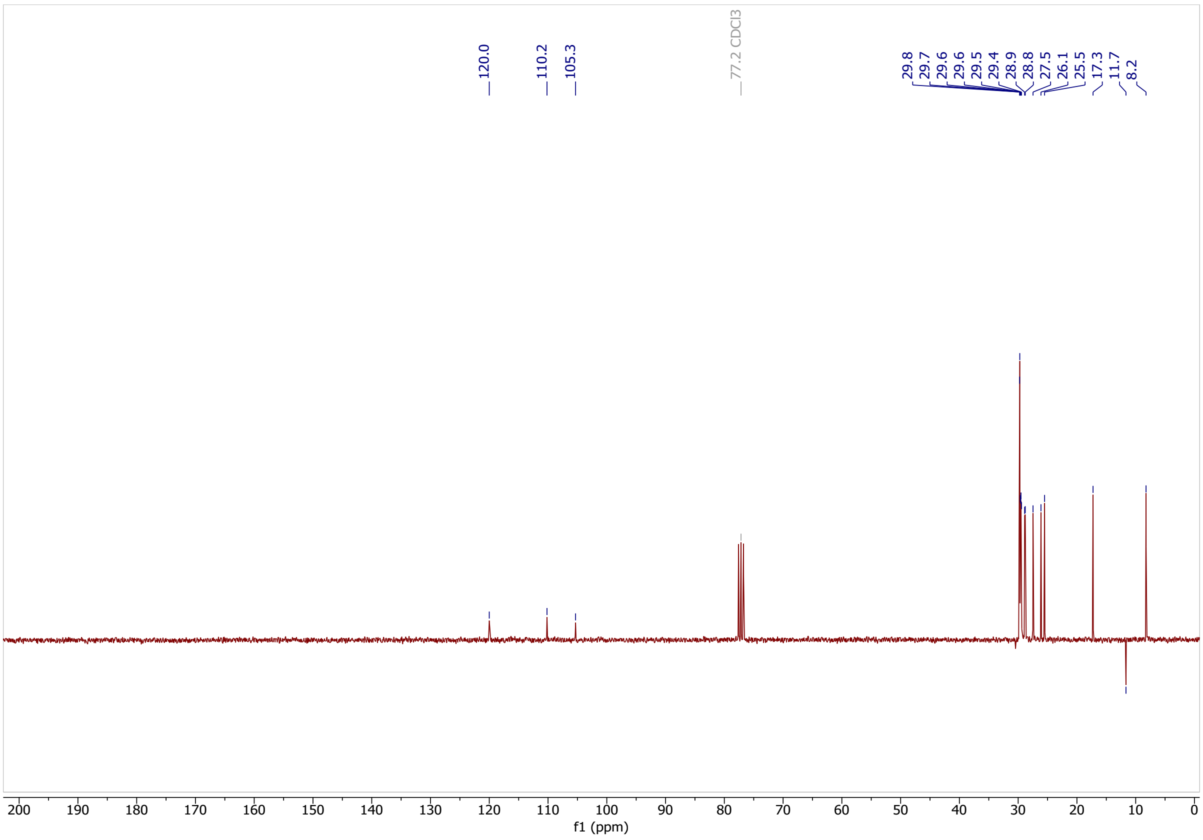

Classified as Internal | Intern

### 15-(2-methylcycloprop-1-en-1-yl)pentadecanoic acid (2)

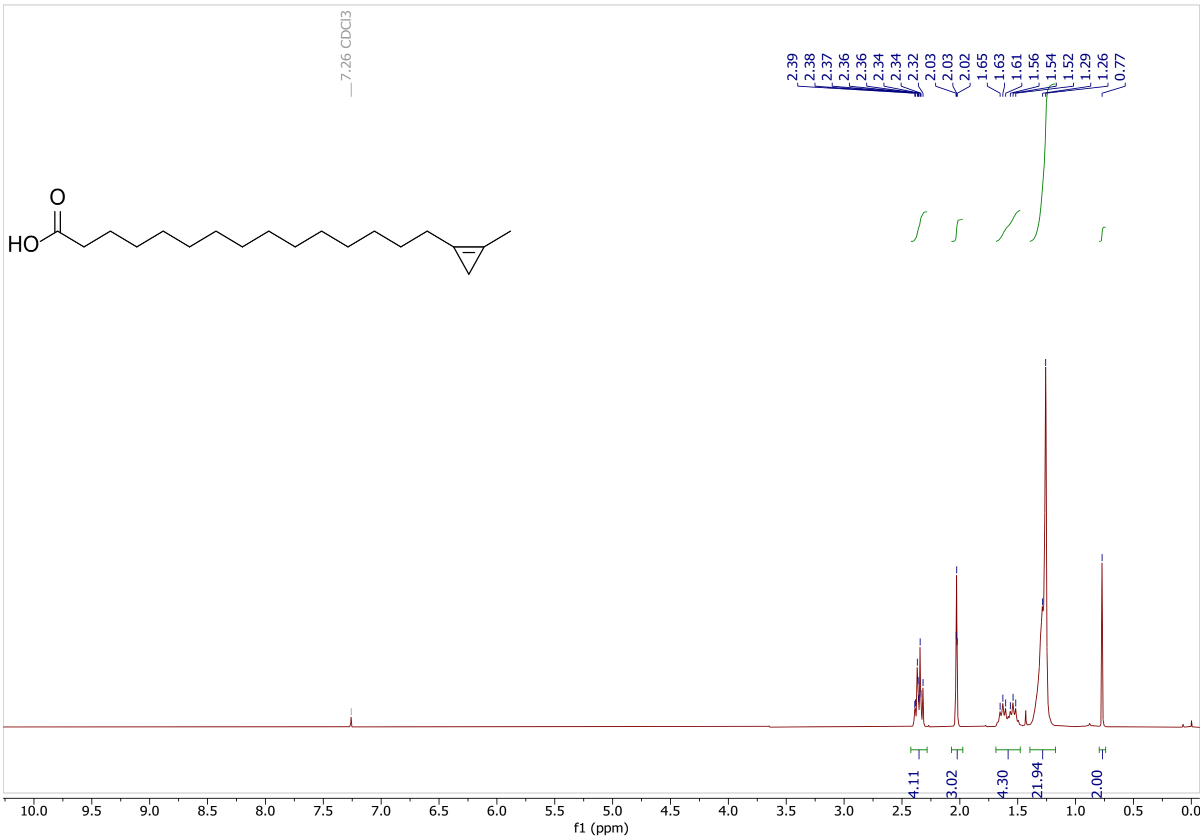

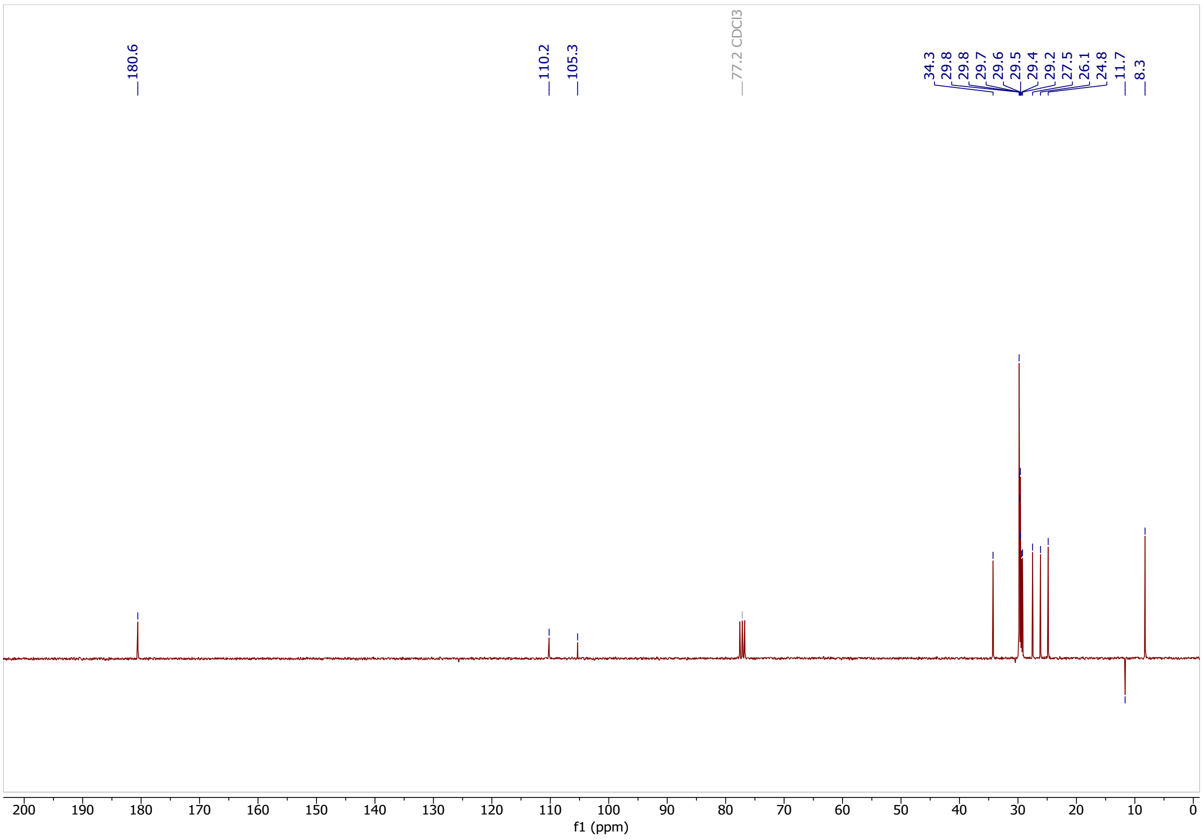

Classified as Internal | Intern

**2-bromodec-1-ene (24)**

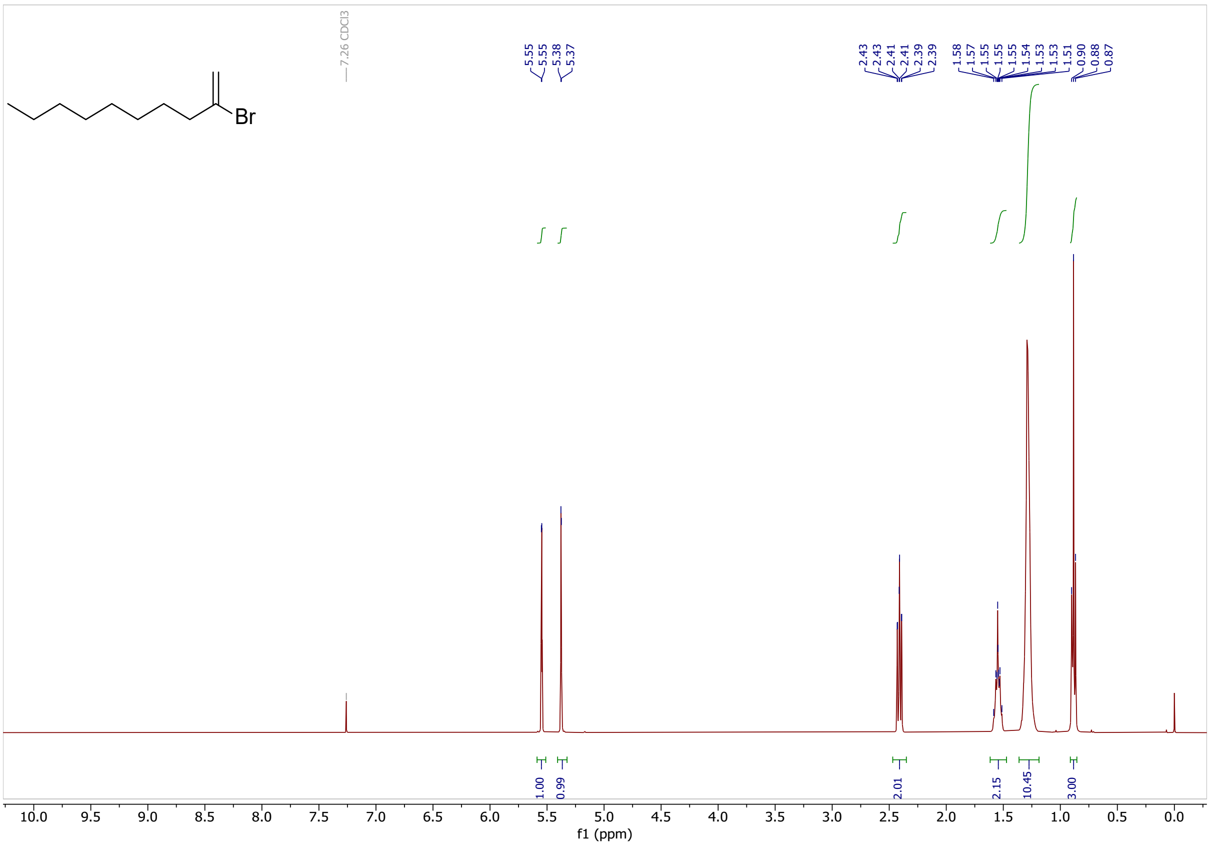

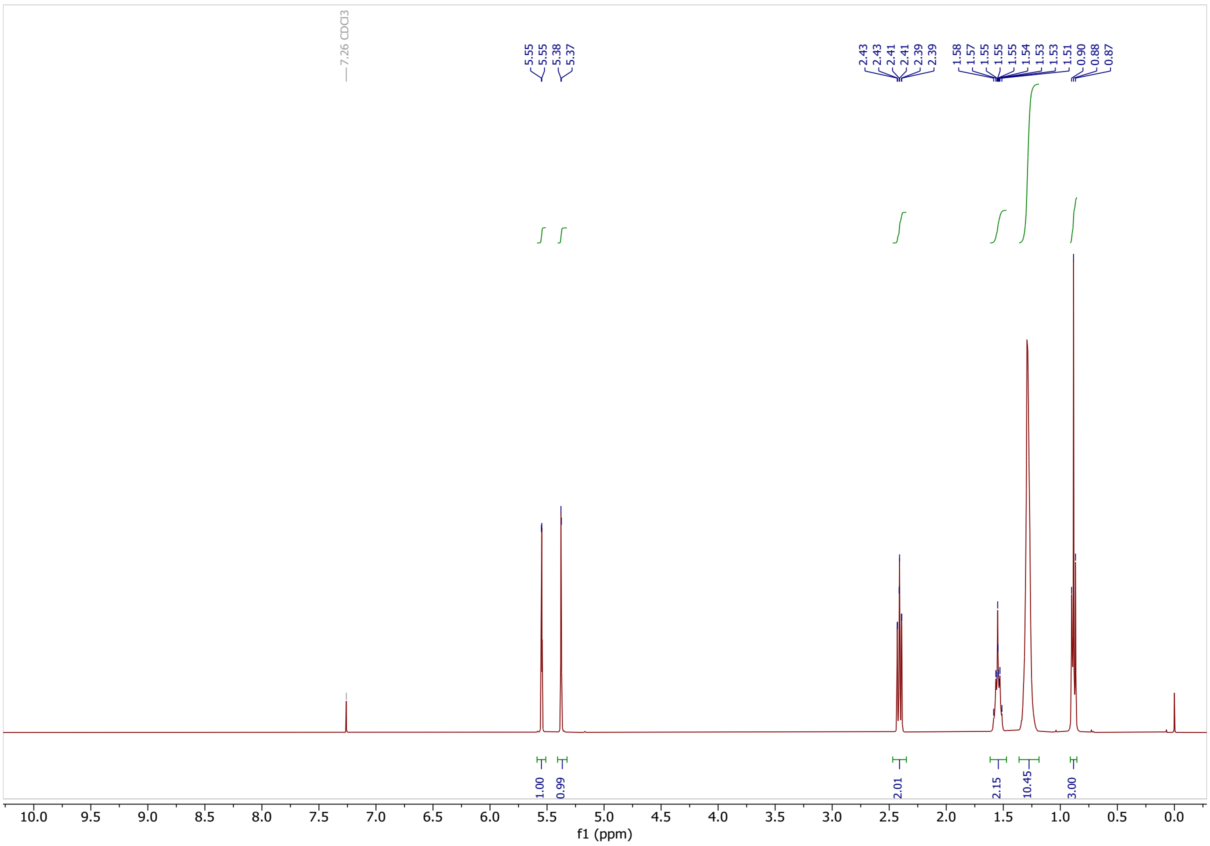

Classified as Internal | Intern

### 1,1,2-tribromo-2-octylcyclopropane (26)

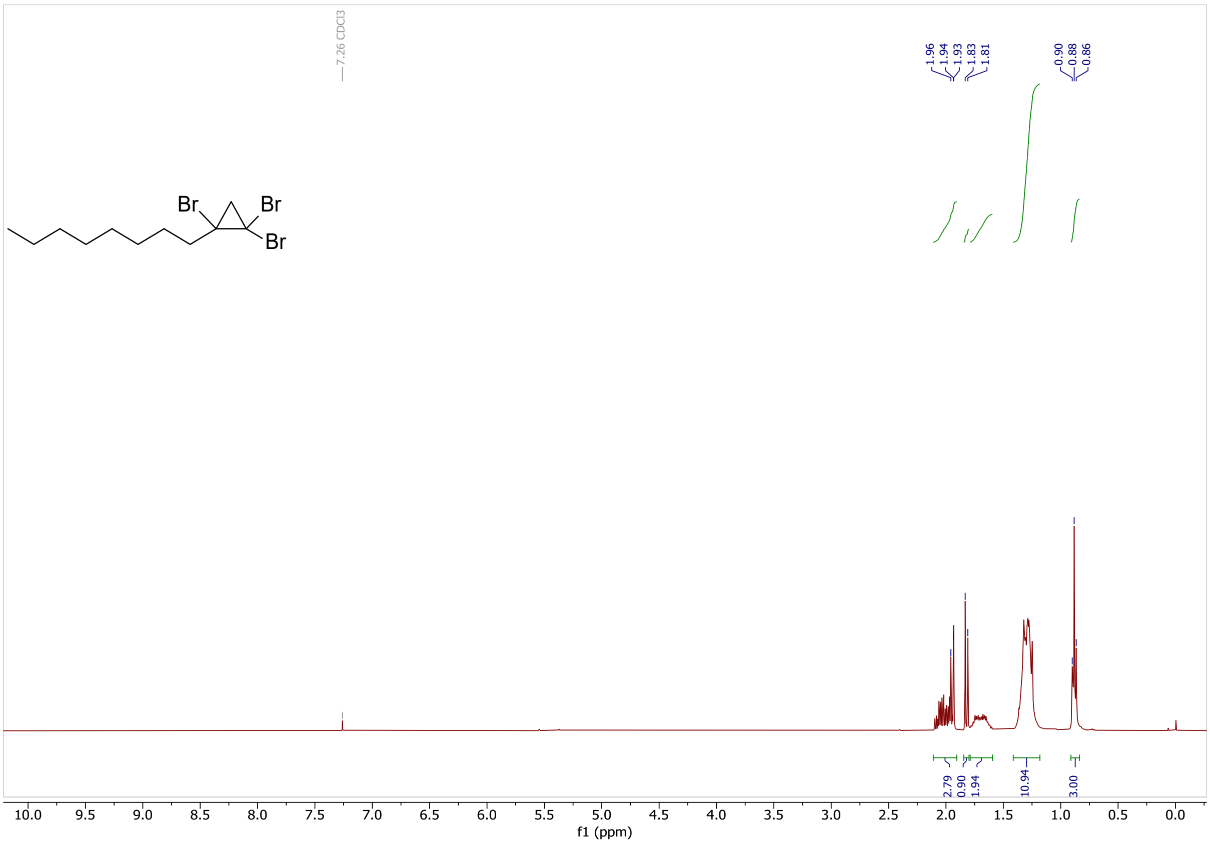

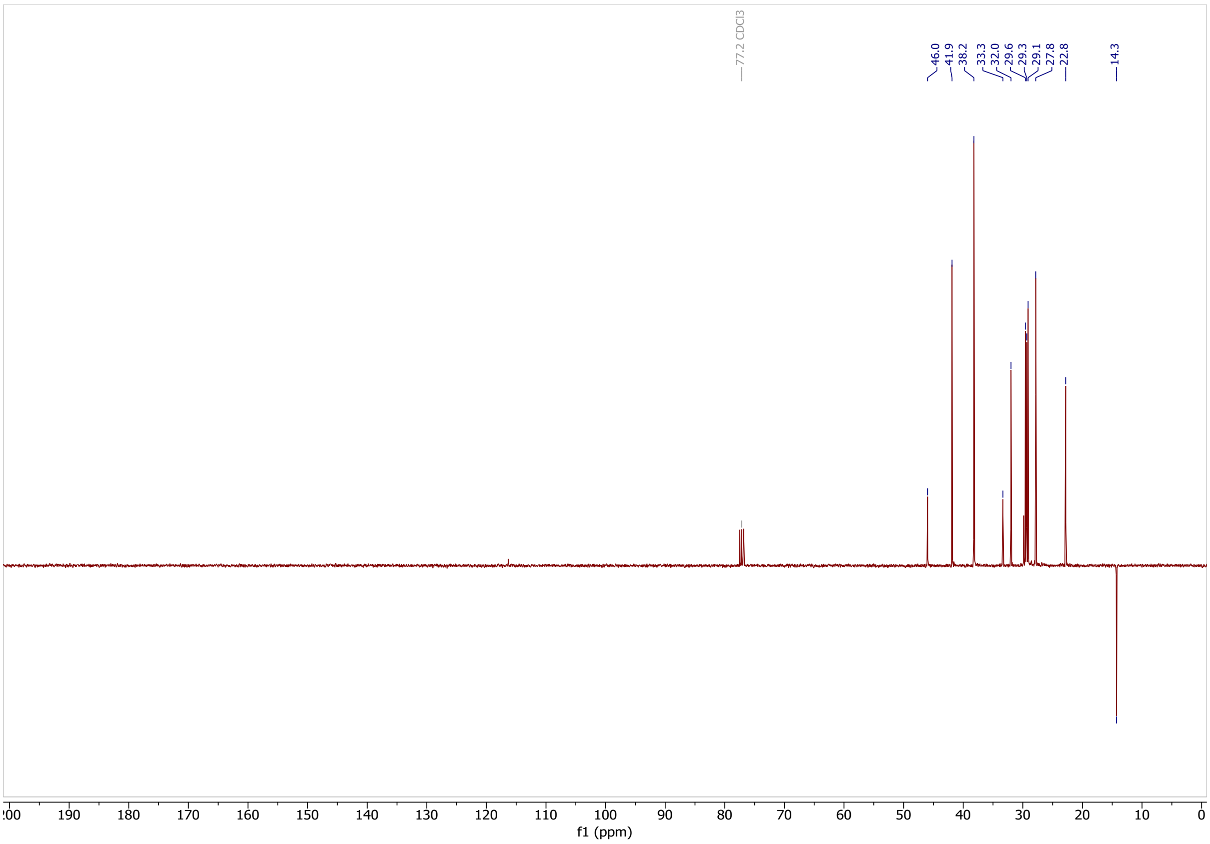

Classified as Internal | Intern

### 1-(7-iodoheptyl)-2-octylcycloprop-1-ene (29)

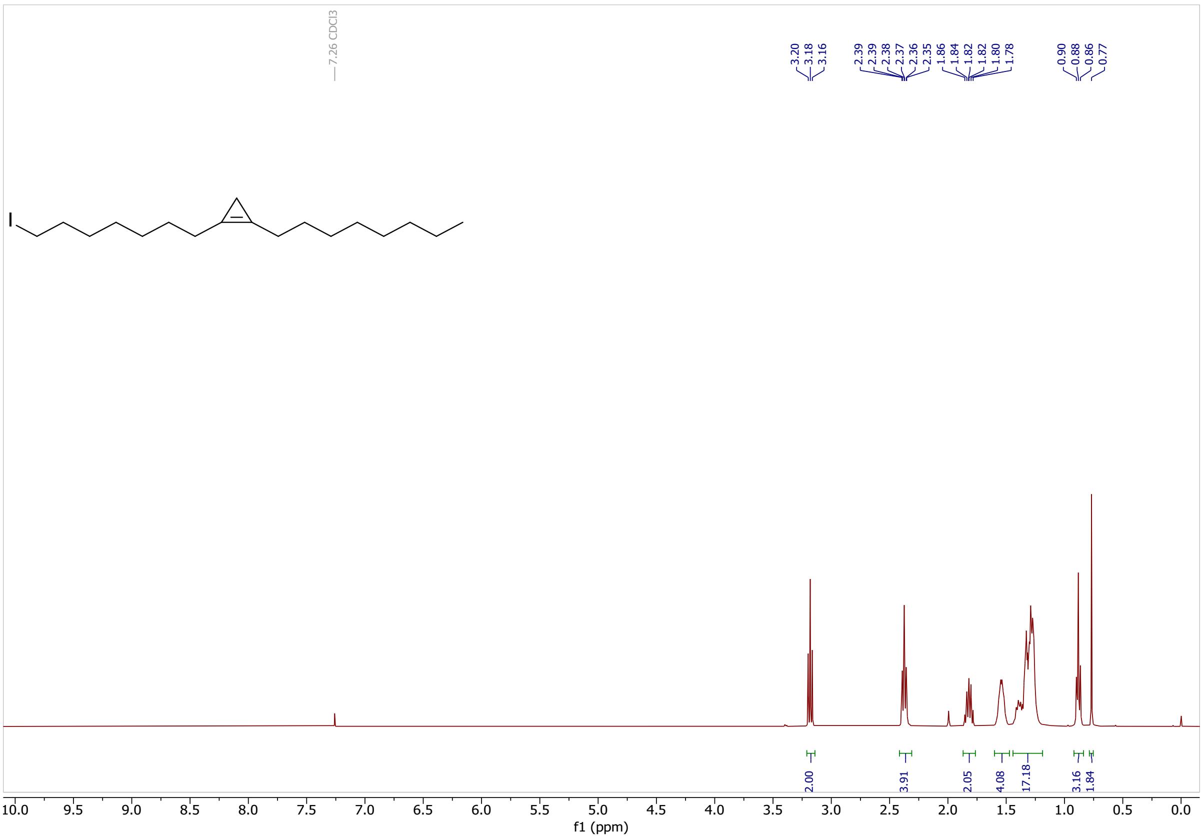

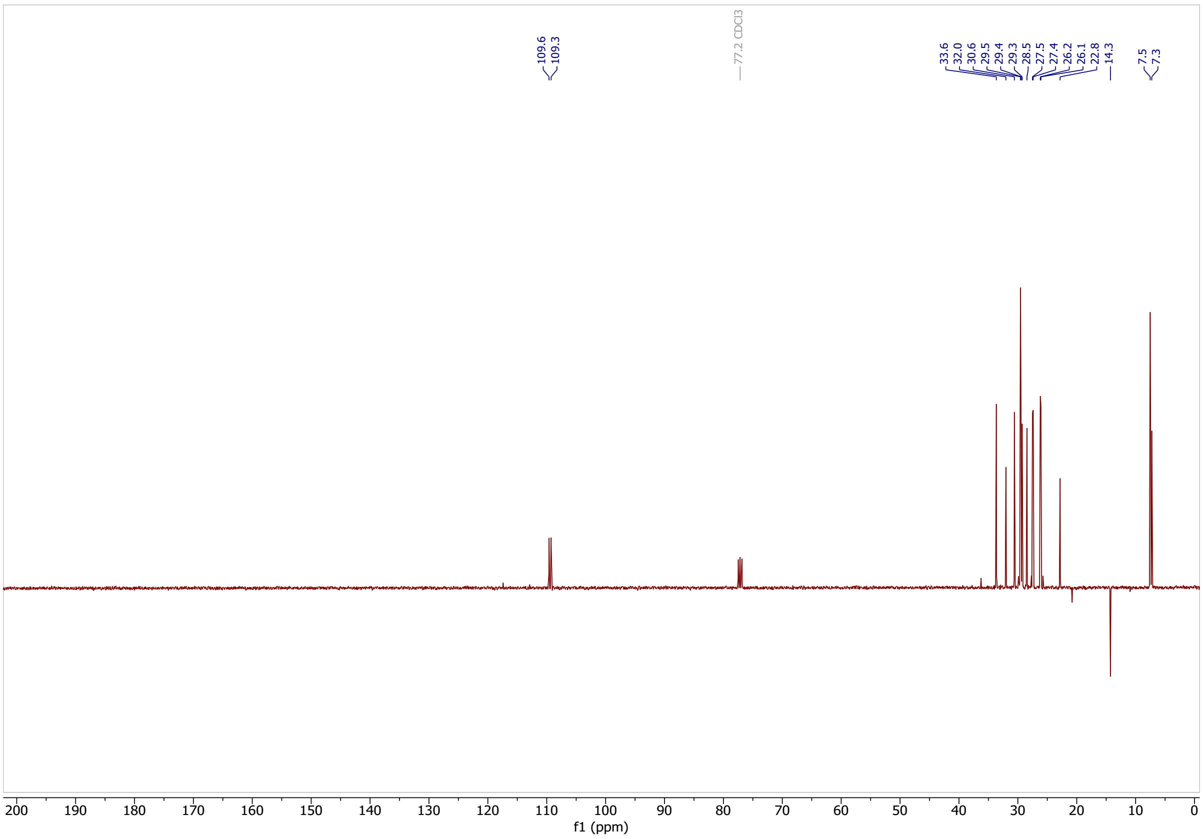

Classified as Internal | Intern

**8-(2-octylcycloprop-1-en-1-yl)octanenitrile (32)**

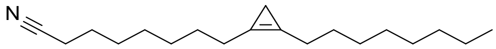

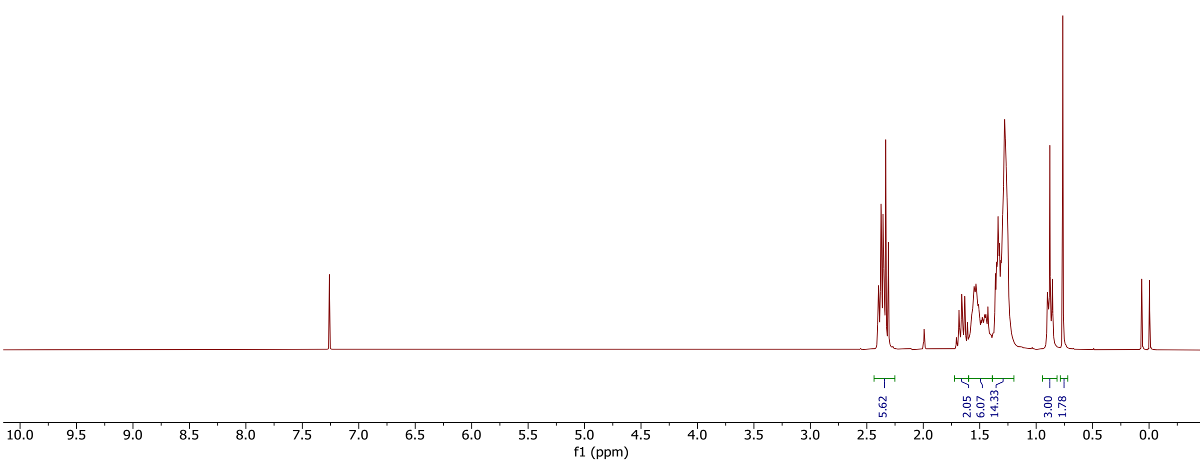

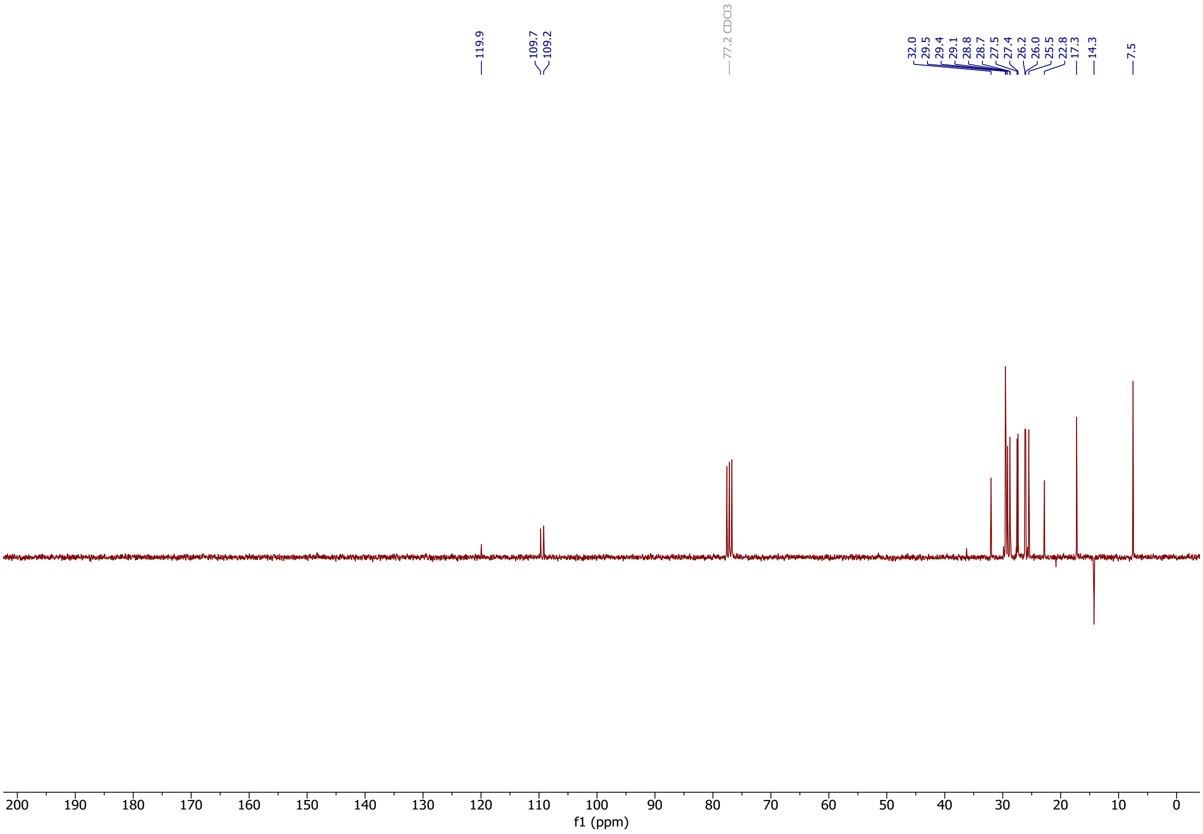

Classified as Internal | Intern

**8-(2-octylcycloprop-1-en-1-yl)octanoic acid (4)**

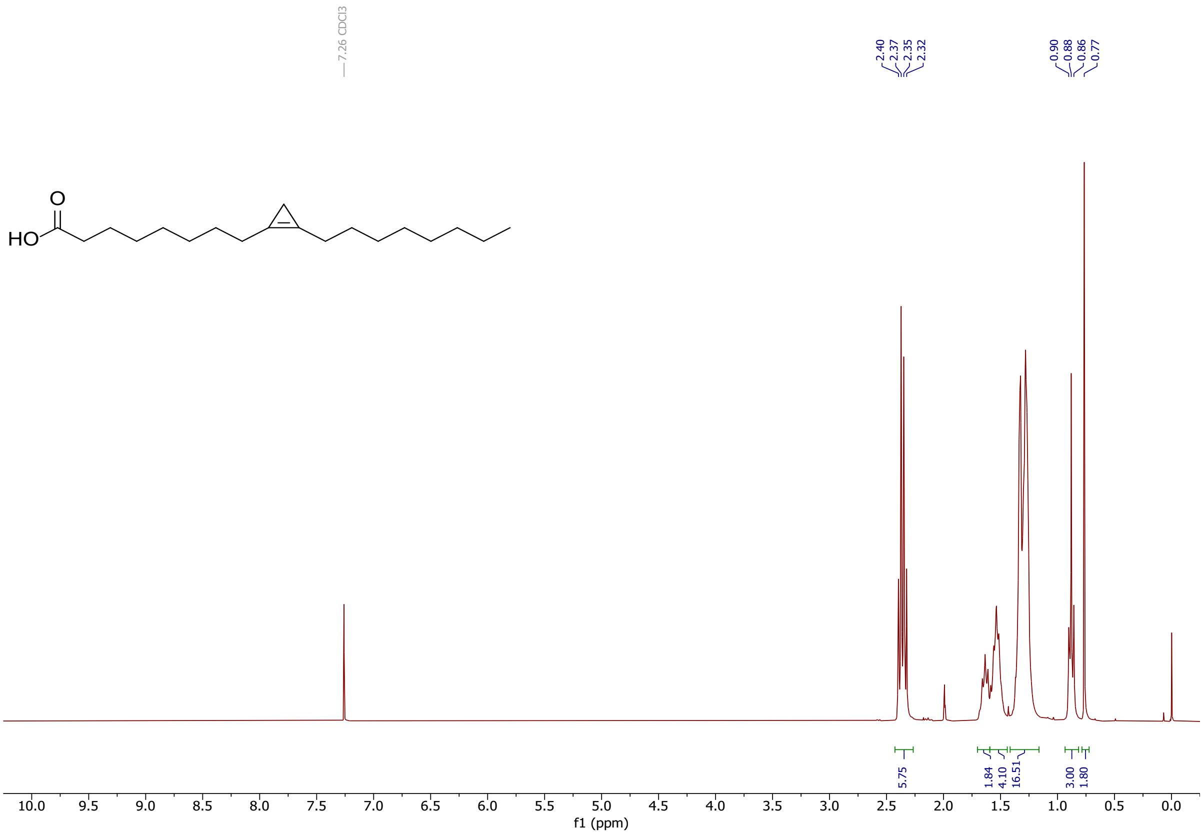

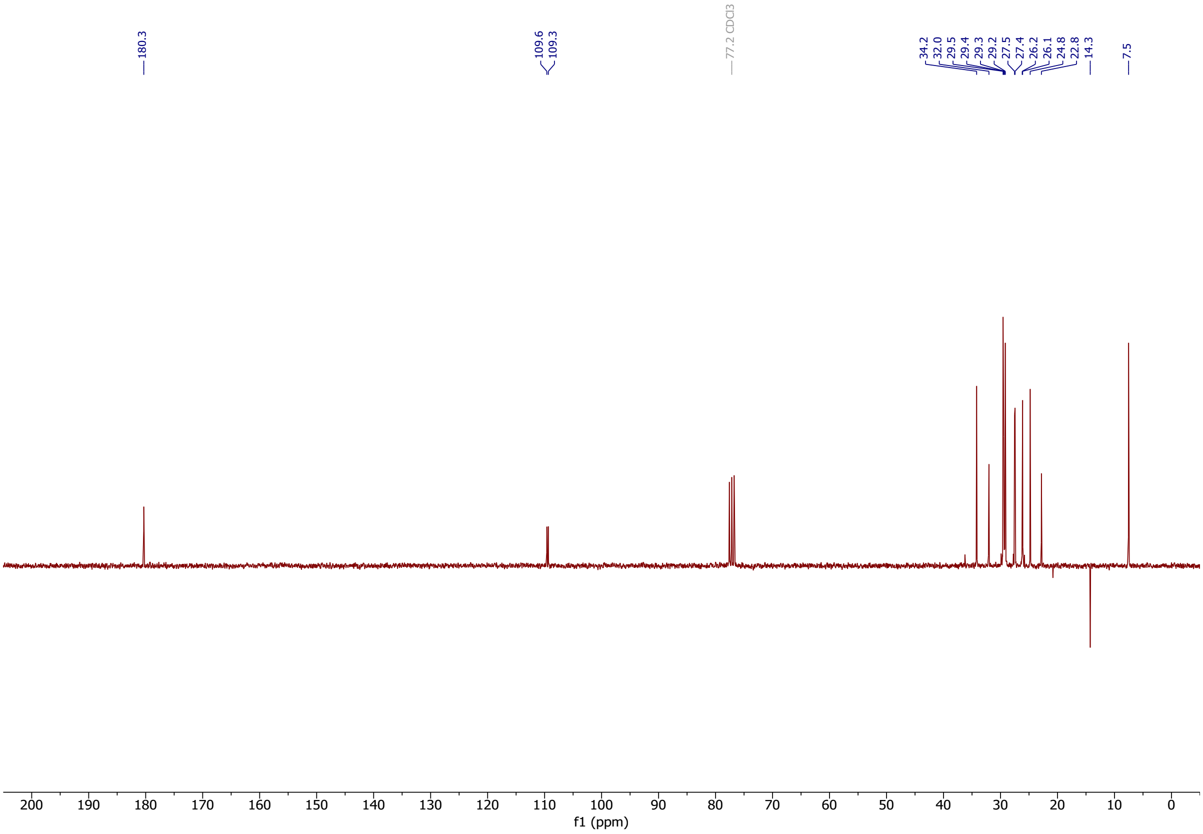

Classified as Internal | Intern

### 9-bromononanoic acid (s4)

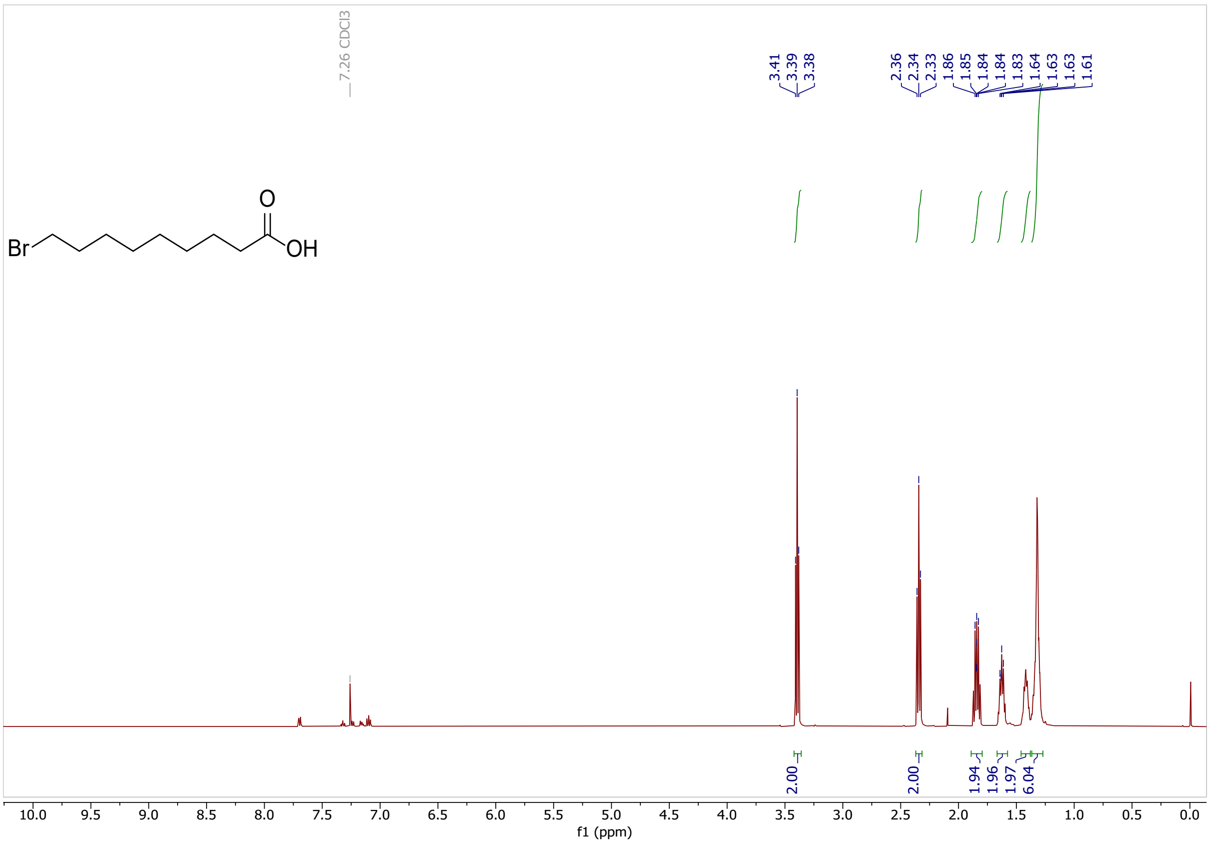

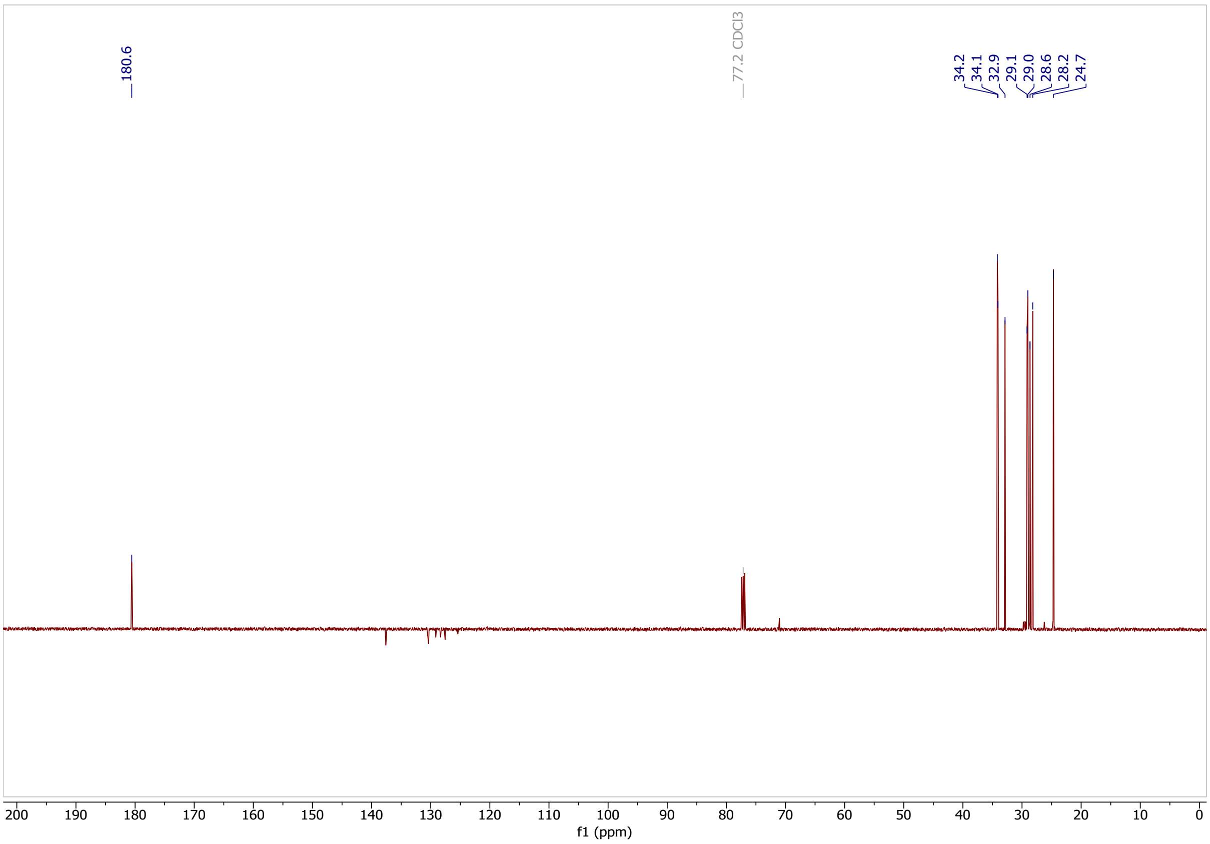

Classified as Internal | Intern

**8-Carboxyoctyl)triphenylphosphonium bromide (s5)**

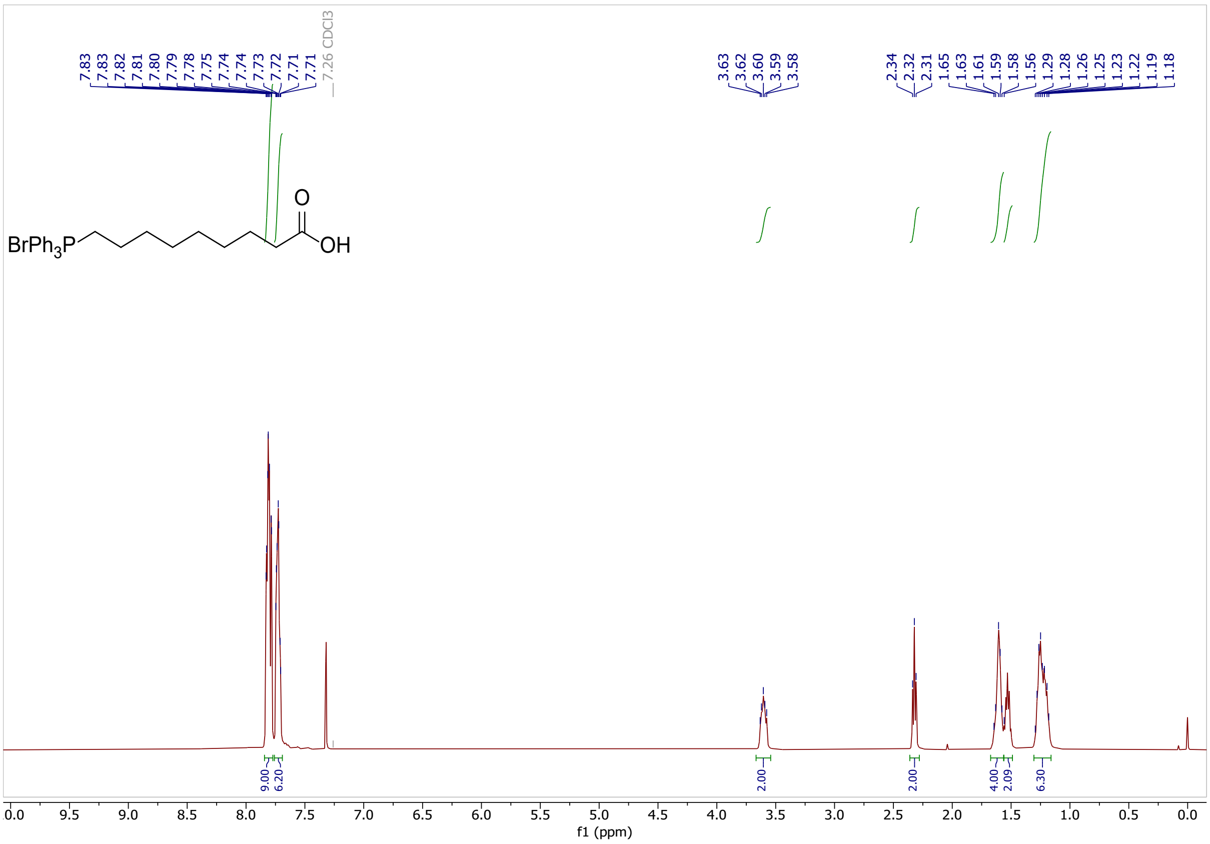

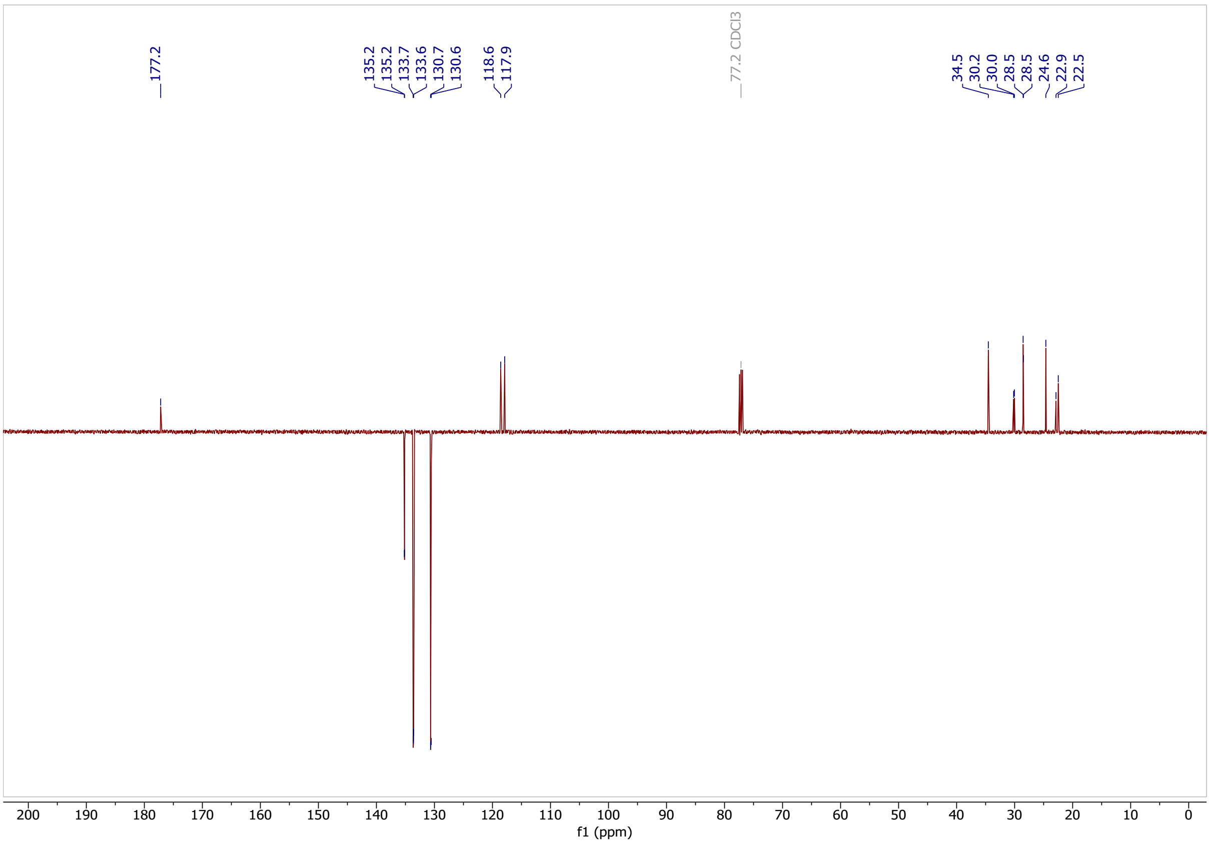

Classified as Internal | Intern

**(2-(1,3-dioxan-2-yl)ethyl)triphenylphosphonium bromide (s6)**

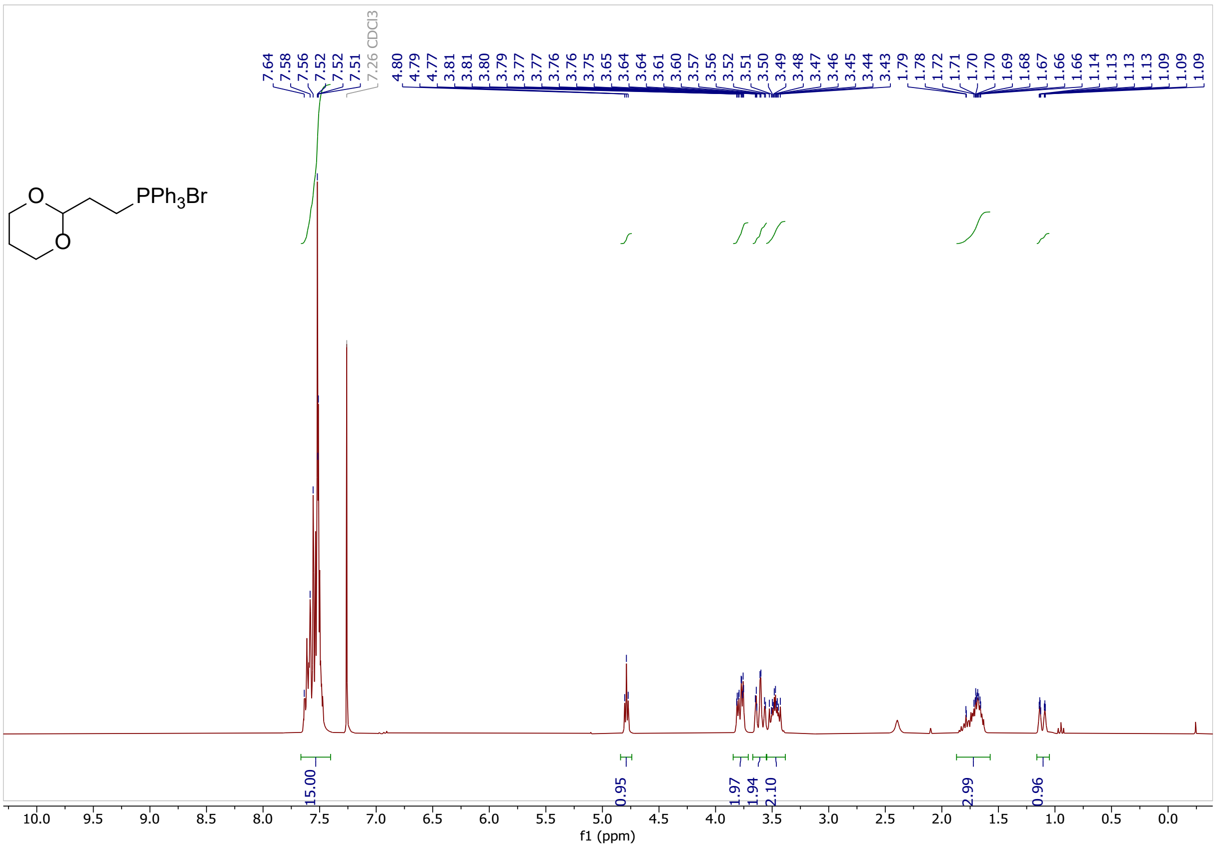

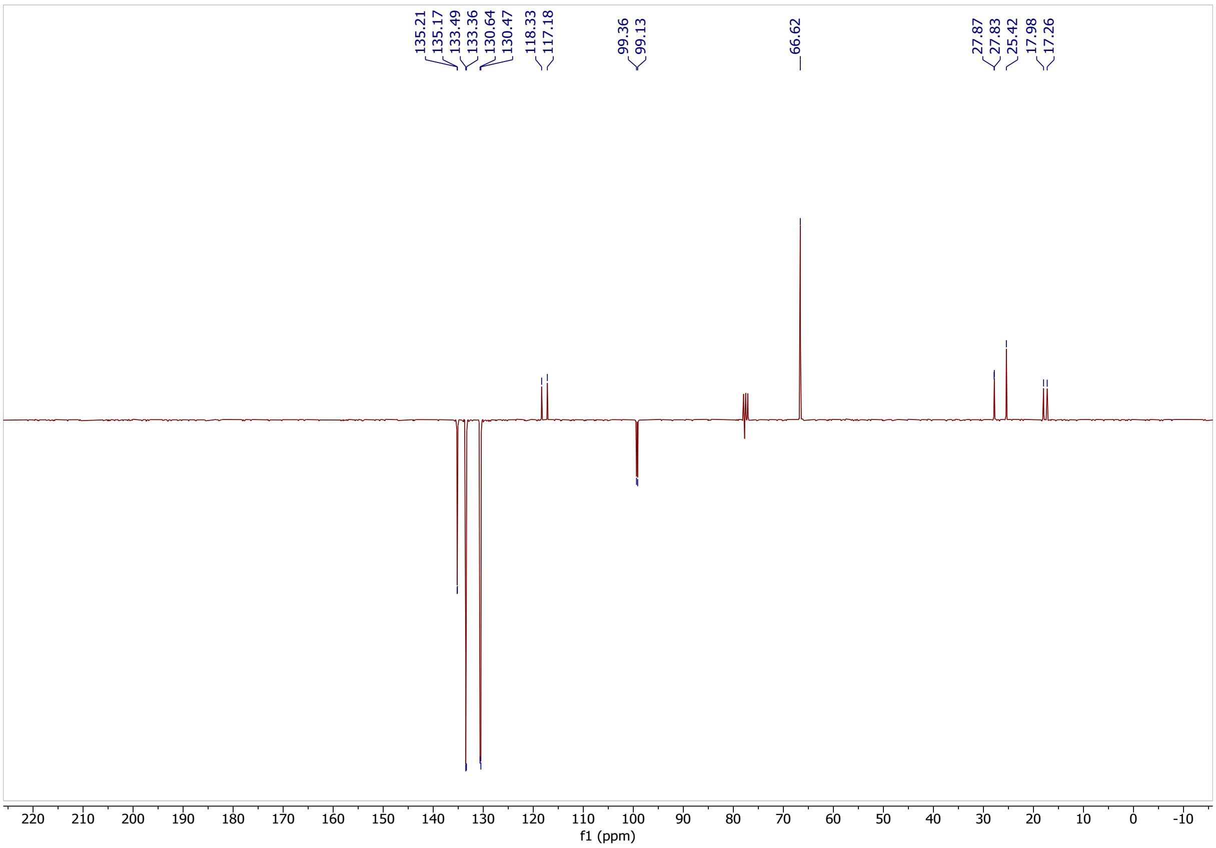

Classified as Internal | Intern

### methyl 5-oxopentanoate (s7)

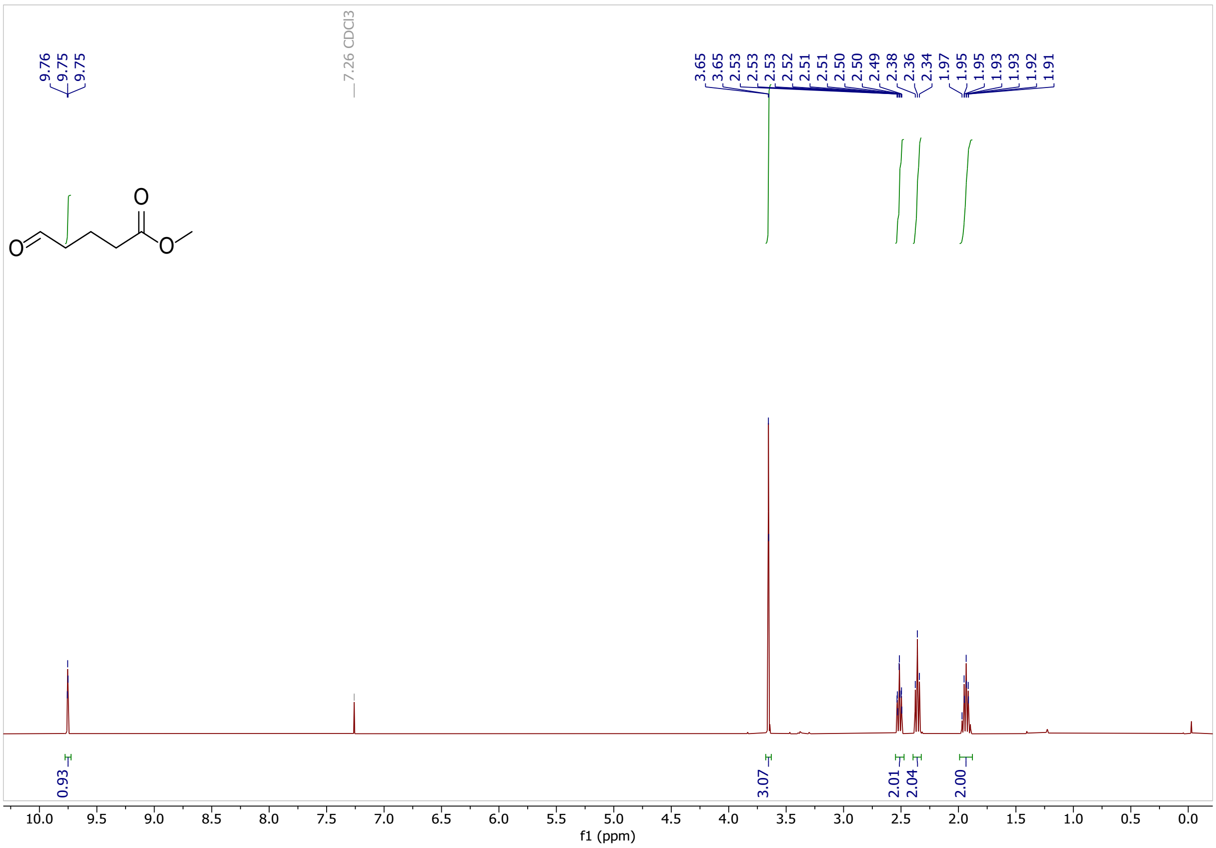

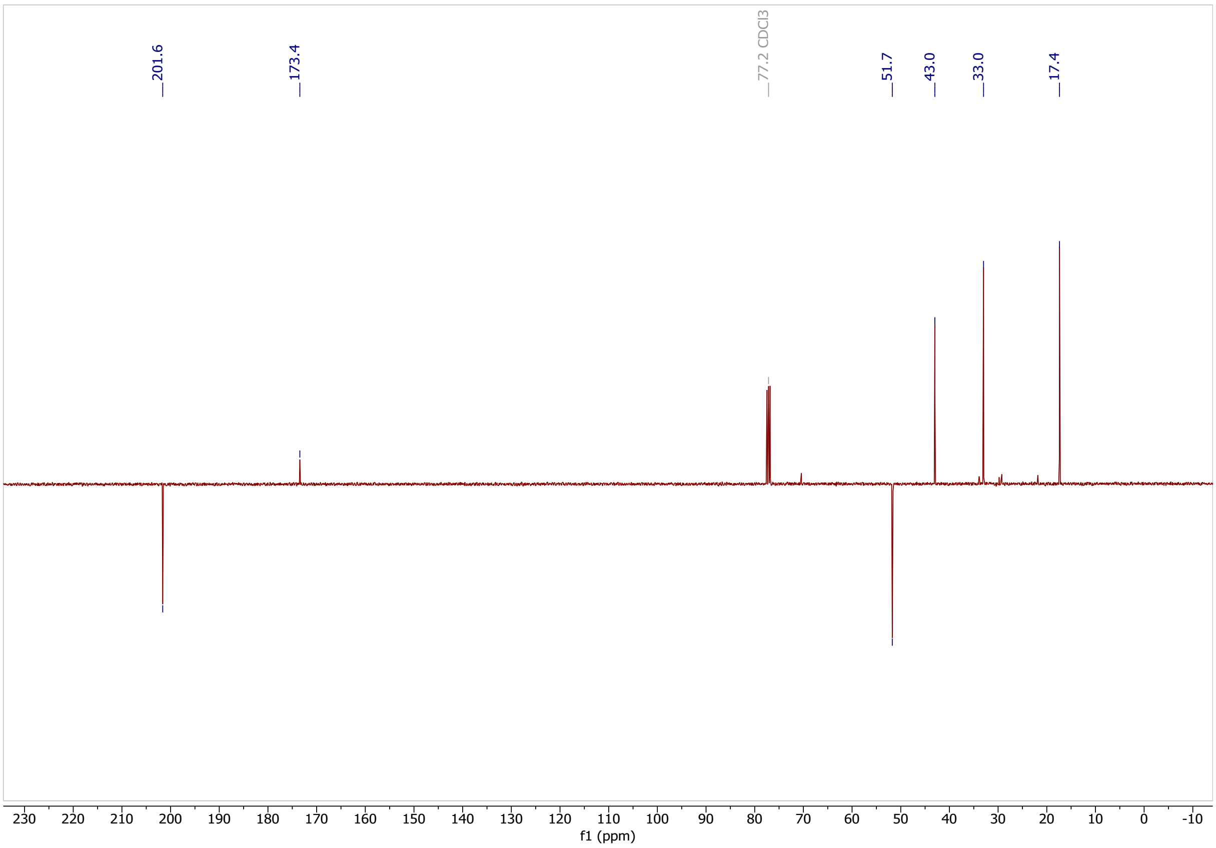

Classified as Internal | Intern

**methyl (Z)-7-(1,3-dioxan-2-yl)hept-5-enoate (s8)**

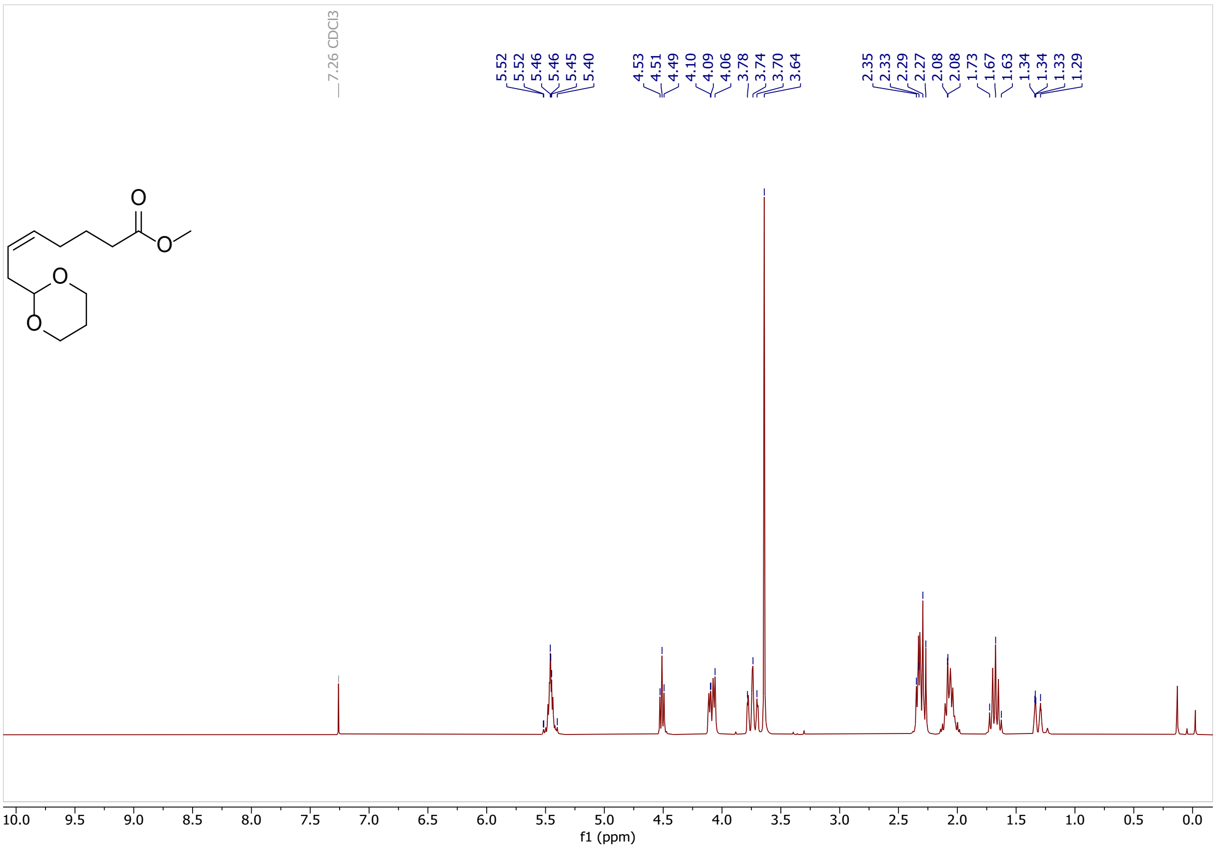

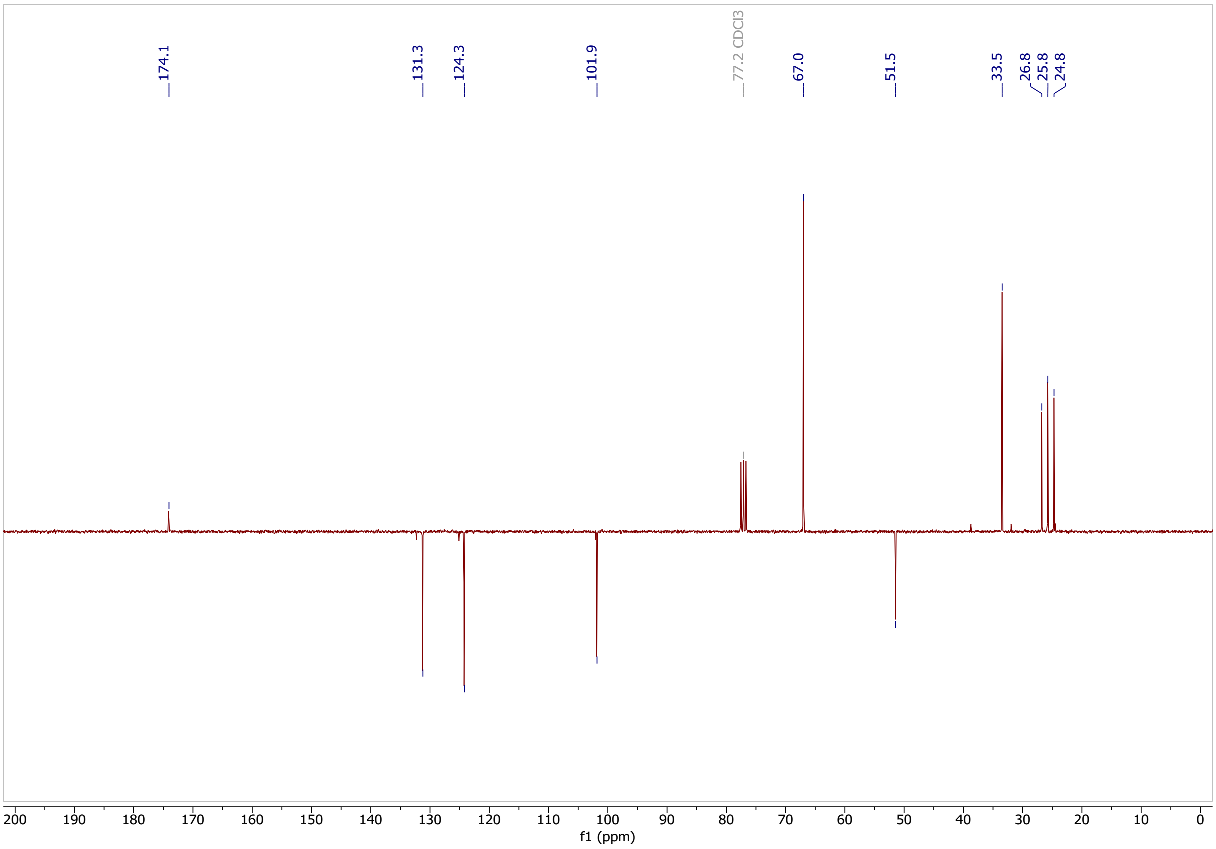

Classified as Internal | Intern

**hexadec-9-yn-1-ol (S9)**

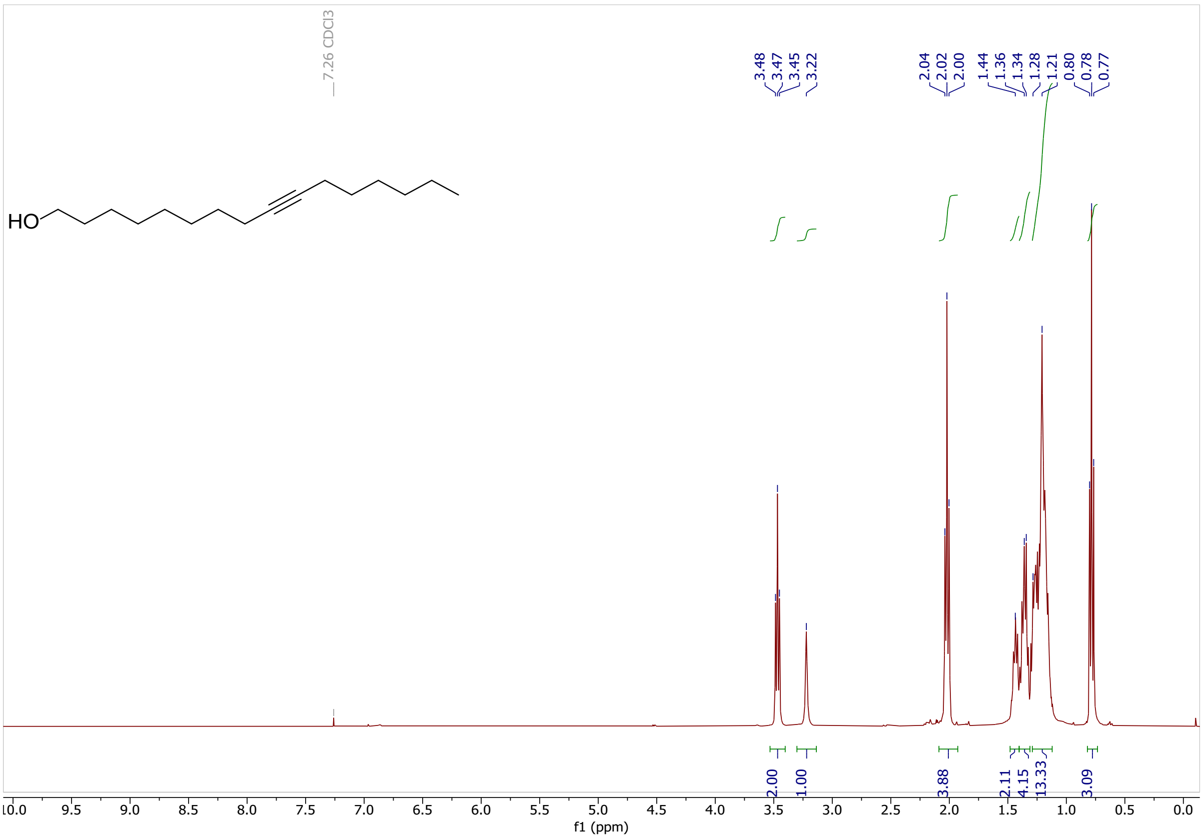

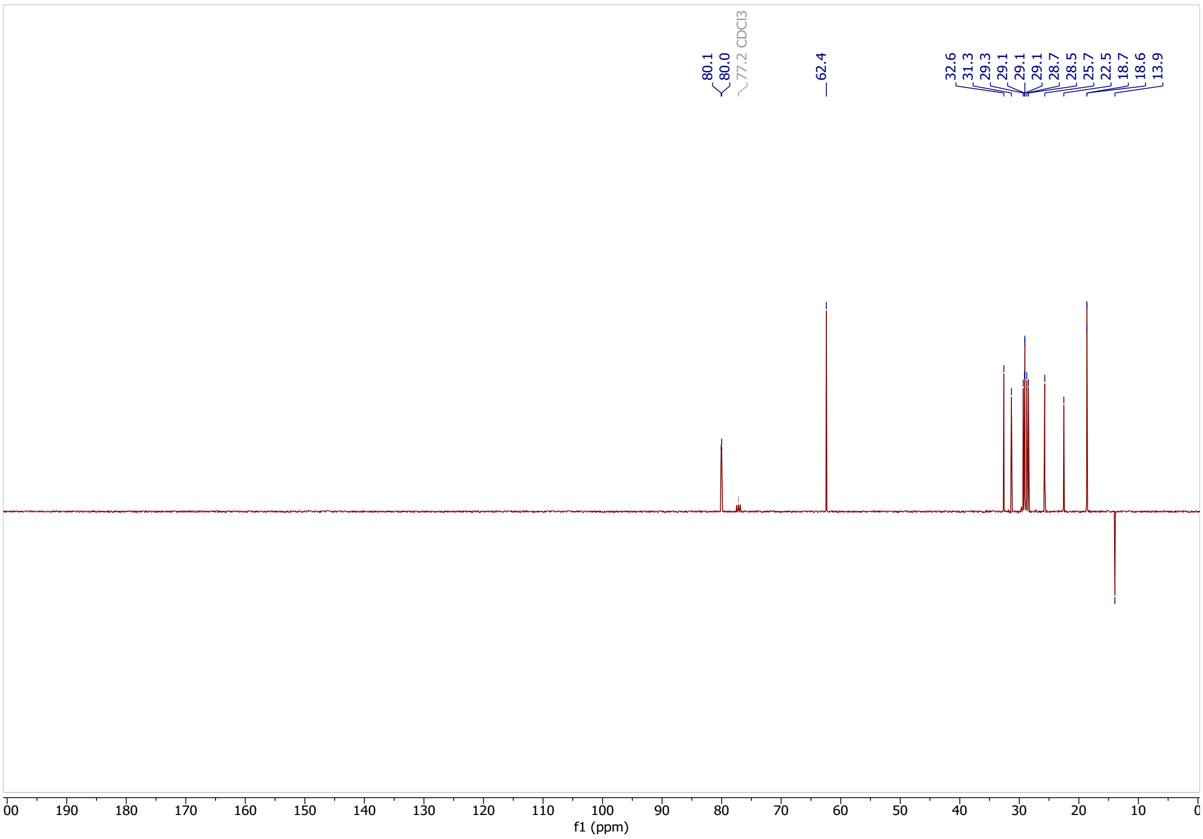

Classified as Internal | Intern

**hexadec-9-ynoic acid (S10)**

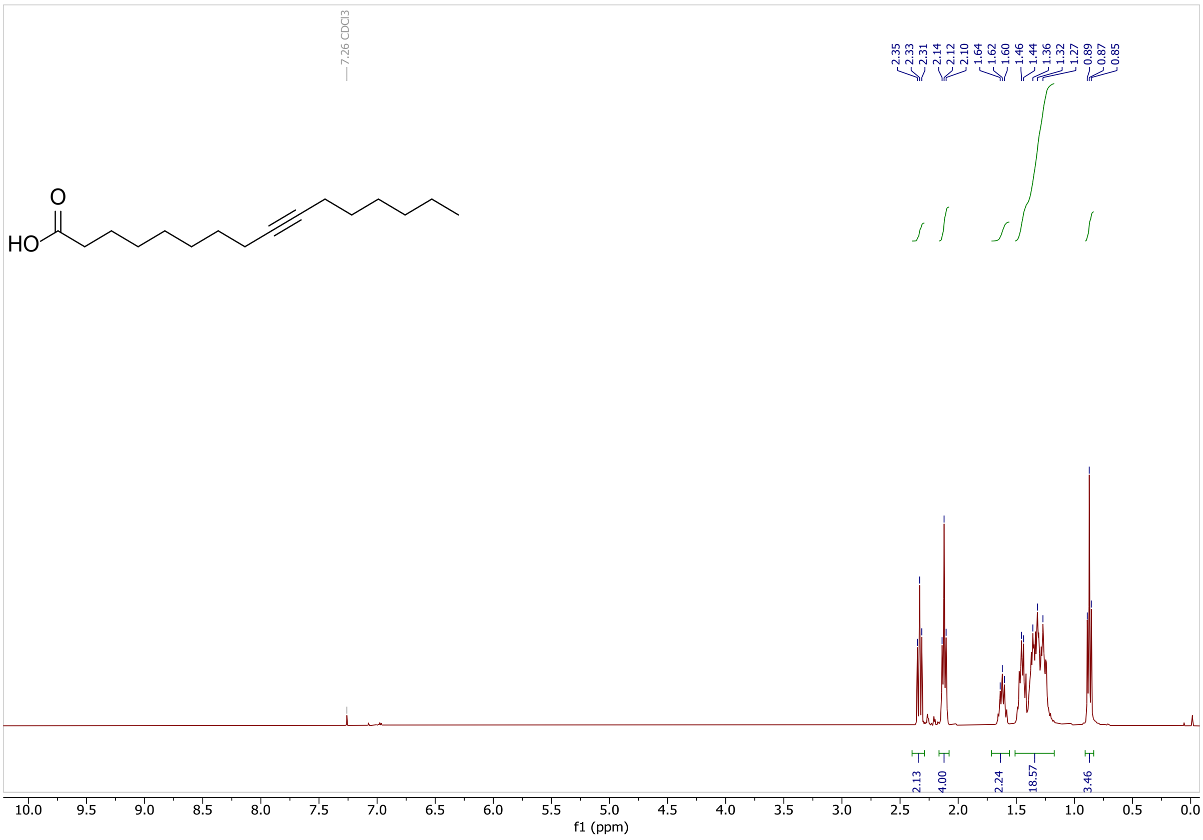

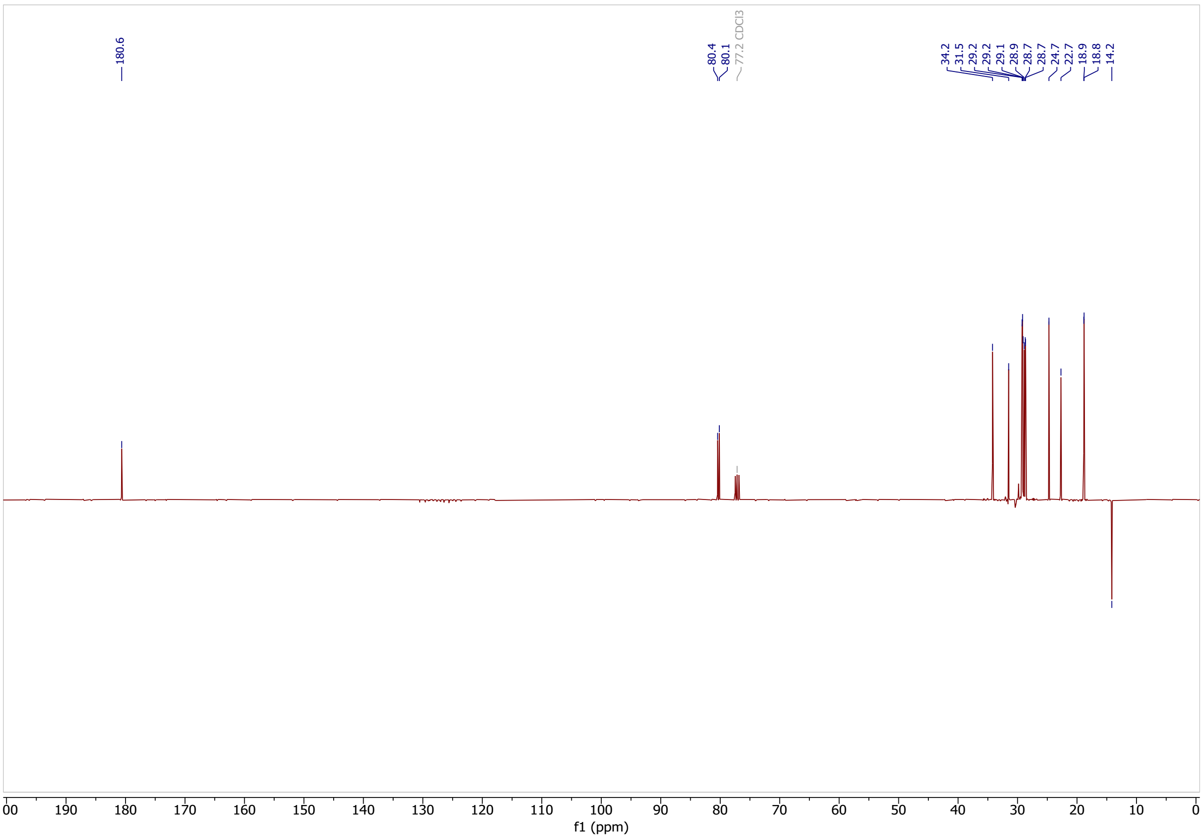

Classified as Internal | Intern

**methyl hexadec-9-ynoate (S11)**

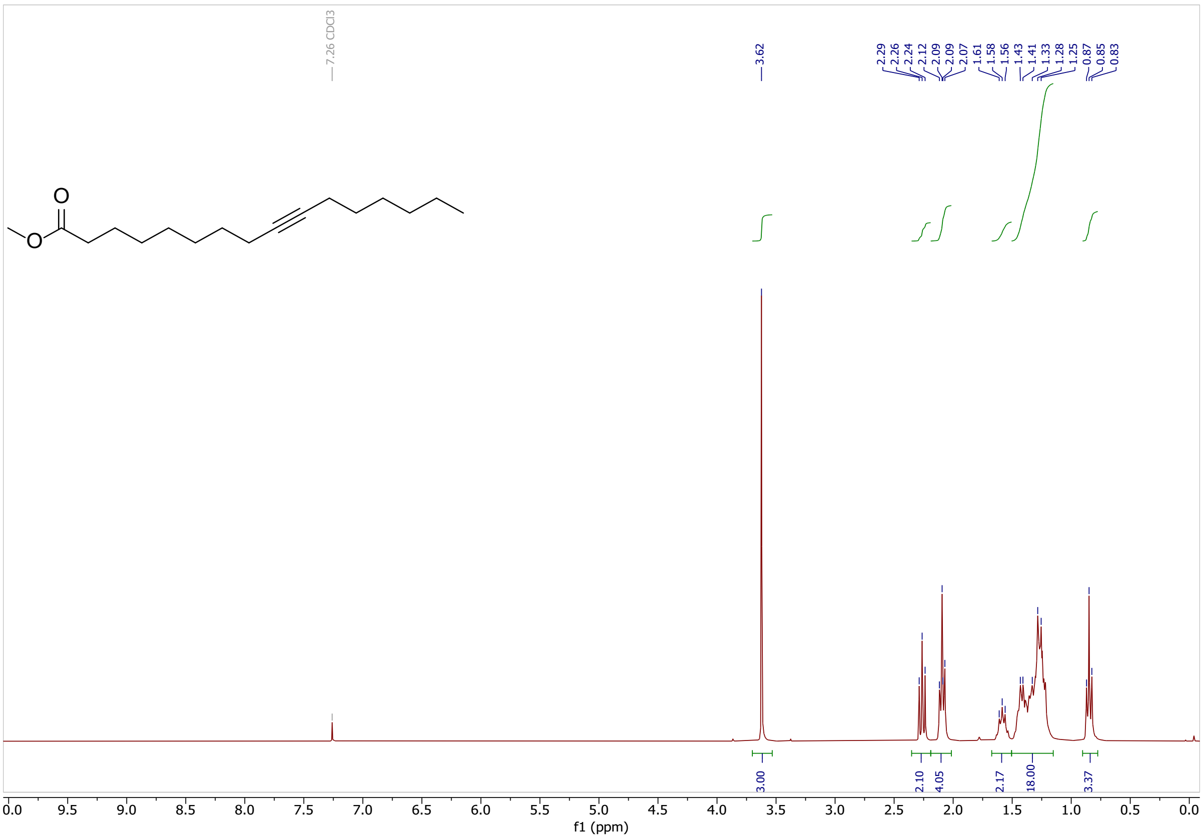

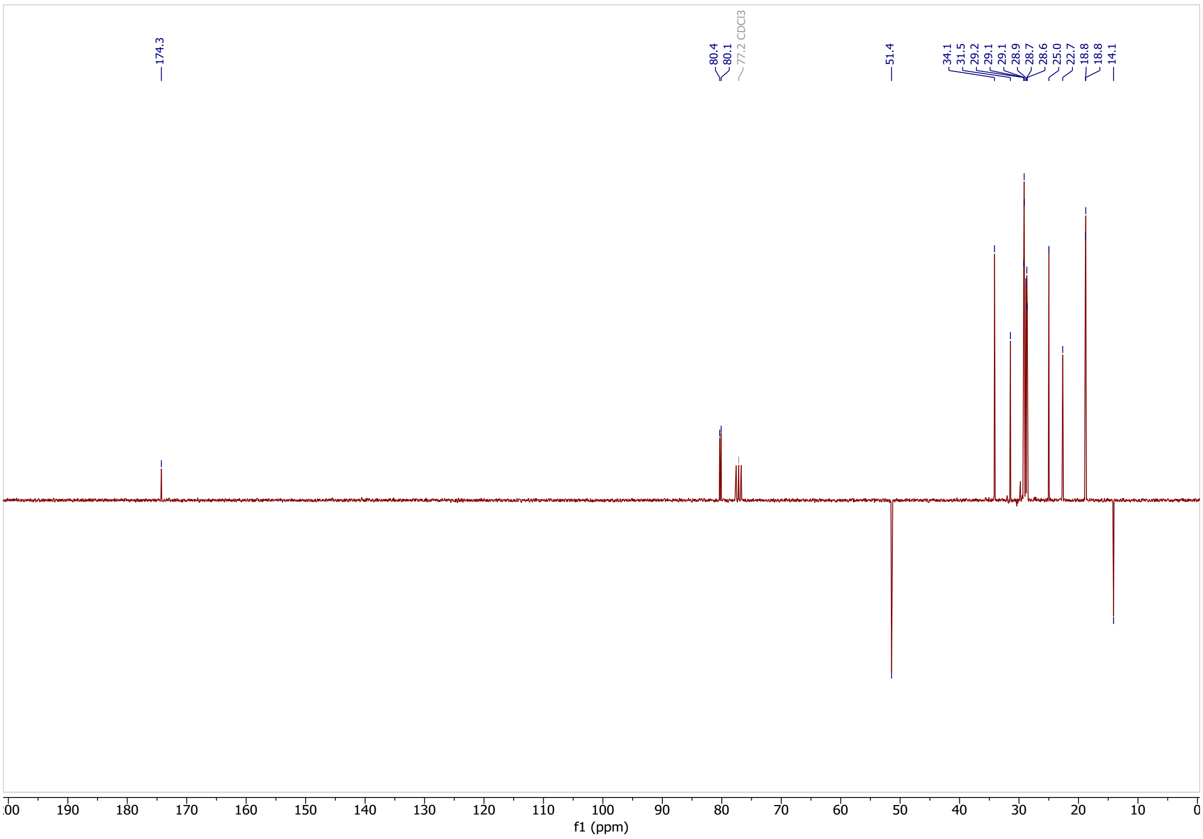

Classified as Internal | Intern

**ethyl 2-hexyl-3-(8-methoxy-8-oxooctyl)cycloprop-2-ene-1-carboxylate (S12)**

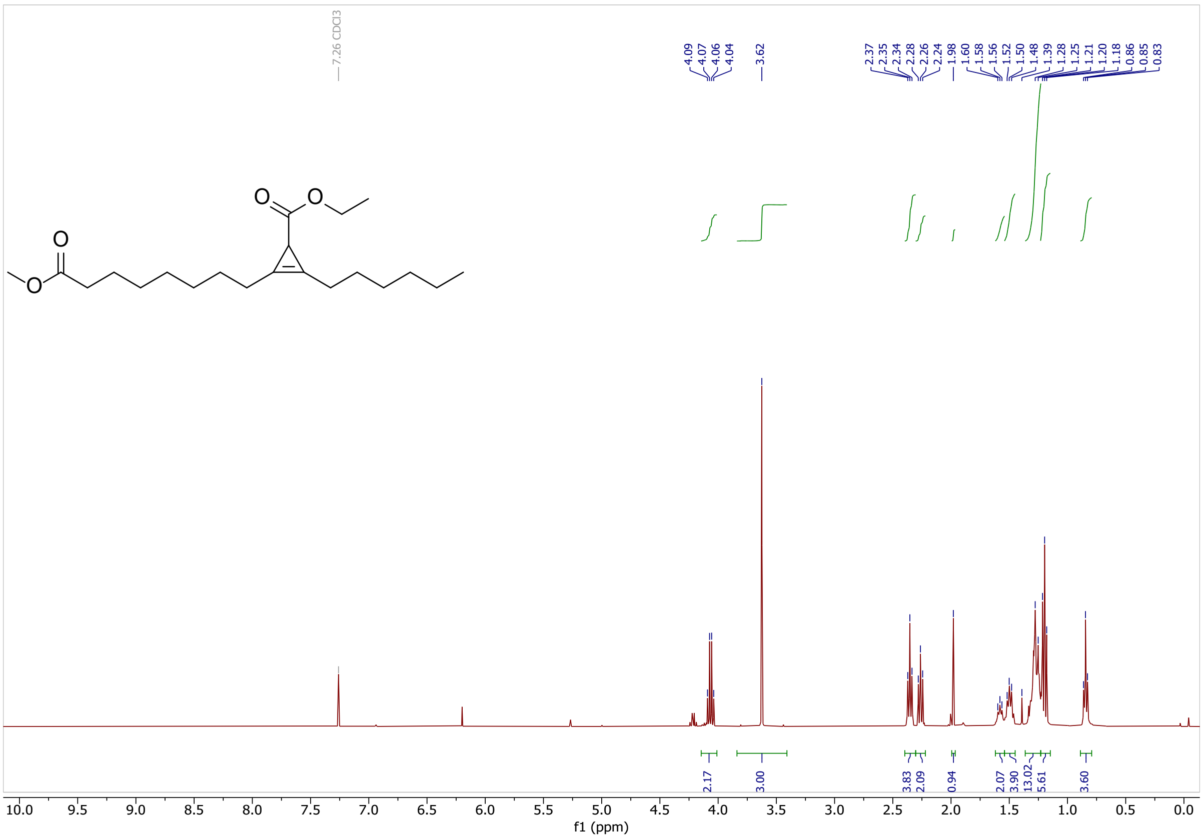

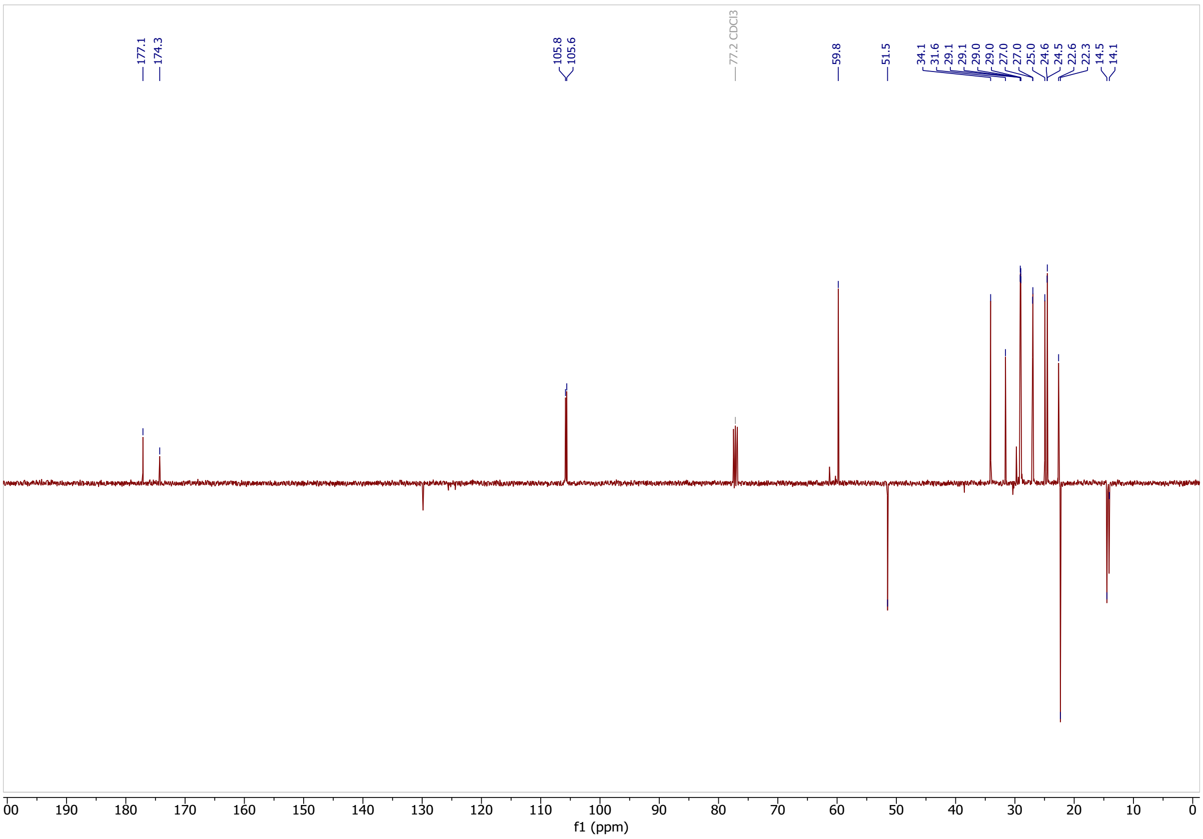

Classified as Internal | Intern

### 2-(7-carboxyheptyl)-3-hexylcycloprop-2-ene-1-carboxylic acid (S13)

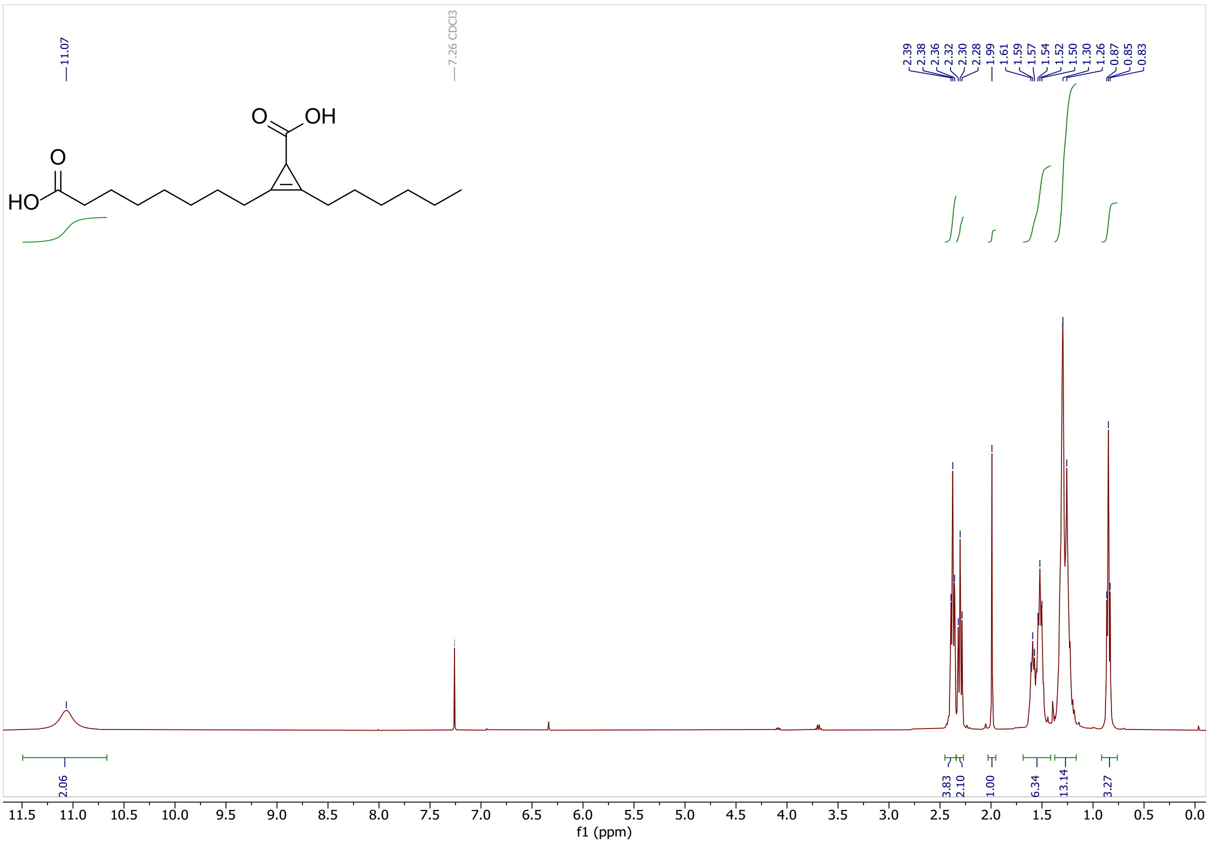

Classified as Internal | Intern

Classified as Internal | Intern
